# Divergent functions of three Kunitz trypsin inhibitor (KTI) proteins in herbivore defense in poplar

**DOI:** 10.1101/2025.09.21.677572

**Authors:** Ishani S. Das, Qianqian Shi, Steven Dreischhoff, Andrea Polle

**Author notes:** Institute of Plant Sciences, Cologne Biocenter, University of Cologne, Zülpicherstr. 47a, 50674 Cologne, Germany (Ishani S. Das). College of Landscape Architecture and Arts, Northwest Agricultural & Forestry University, Yangling, Shaanxi, 712100, China (Qianqian Shi).

## Abstract

**Background:** Climate warming promotes the expansion of insect pests. Among the inducible defense responses activated by attacked plants, Kunitz trypsin protease inhibitors (KTIs) play an outstanding role. KTIs affect food digestion and thereby control the fitness of herbivorous insects. Poplars contain an expanded family of KTIs, whose distinct intrinsic functions are under investigation. Here, we set out to identify KTIs with anti-herbivore activity and assessed the potential growth trade-off incurred by high KTI expression levels.

**Results:** We identified 28 KTIs in the haploid genome of *Populus* x *canescens*, 21 of them were responsive to herbivory. The greatest induction was observed for *KTI_400, KTI_600* and *KTI_0882* (*P. trichocarpa* orthologues Potri.019G124400, Potri.019G124600, Potri.019G088200), whereas a moderate response was found for *KTI_53200* (Potri017G153200 orthologue), a protein mainly localized in the xylem sap. Mechanical wounding and methyl-jasmonate resulted in fast and strong induction of *KTI_400* and *KTI_600* and moderate or lacking responses in *KTI_0882* and *KTI_53200*. Increased *KTI* expression levels were associated with upregulation of *ALLENE OXIDE SYNTHASE* (AOS), whereas exposure to compounds eliciting ethylene or salicylic acid signaling did not affect *KTI*s. We generated stable CRISPR/Cas12a-mediated knock-out and *p35S*-mediated overexpression lines of *KTI_400, KTI_600* and *KTI_53200* in *Populus* x *canescens.* Among the wildtype and transgenic lines, only *kti_400+kti_600* double knock-out lines produced greater biomass. Larvae of *Helicoverpa armigera,* a pest expanding in Europe due to a warmer climate, were allowed to feed on wildtype and transgenic poplar lines. Transgenic poplars overexpressing *KTI_400* or *KTI_600* resulted in reduced and their double knock-out lines in increased weight gain of the larvae. In contrast, overexpressing or knock-out lines of *KTI_53200* had no effect on larval weight gain compared with controls.

**Conclusion:** KTI_400 and KTI_600 are potent, natural *in-planta* anti-herbivorous agents. Their expression is associated with larval growth reductions. Modulation of KTI_53200 levels had no direct effects on the fitness of leaf-feeding *H. armigera* or on plant growth. This study sheds light on the potential application of *KTI* in plant defenses and biocontrol against herbivores in trees and presents new options to investigate growth-defense theories.

## 1. Background

Forests are facing an escalating array of anthropogenic-induced changes to their environment. Rising global temperatures, driven by human activities, have led to extended breeding seasons and increased generation cycles for insect herbivores (Pureswaran et al., 2018). Herbivory and pathogens pose significant threats to the growth and recruitment of poplars in northern biomes (Pureswaran et al., 2018). As key stone species, poplars play a crucial role in supporting associated organisms (Rogers et al., 2020), and their industrial relevance as a feedstock for biofuels and bioproducts only adds to their importance (Stanton et al., 2021). Consequently, a deeper understanding of how poplars respond to and adapt to environmental cues is a pressing need.

Poplars possess a range of constitutive and inducible defenses, including metabolite- and protein-based mechanisms (Constabel and L. Lindroth, 2010; Divekar et al., 2022). Among these defenses, proteinase inhibitors are of particular interest due to their role in protecting against herbivory (Grosse-Holz and van der Hoorn, 2016). Proteinase inhibitors are widespread throughout the plant body (Ma et al., 2011; Major and Constabel, 2008; Philippe et al., 2009). When ingested by leaf-feeding herbivores, these proteins are absorbed and inhibit the activity of digestive enzymes in the insect’s gut (Pureswaran et al., 2018). This inhibition leads to impaired food digestion, reduced dietary assimilation, and, in some cases, mortality of the herbivore (do Amaral et al., 2022; Arnaiz et al., 2018; Mehmood et al., 2024; Oliveira et al., 2009).

Protease inhibitors interact with different classes of proteases (Clemente et al., 2019; Fear et al., 2007; Rustgi et al., 2017). Kunitz Trypsin Inhibitor (KTI) proteins are part of the serine protease inhibitor family, known as “serpins”, and target serine proteases such as trypsin and chymotrypsin (do Amaral et al., 2022; Mehmood et al., 2024; Oliveira et al., 2009). KTIs have a molecular mass of approximately 21 kDa, with approximately 120 amino acids and four cysteine residues. These cysteine residues form two disulfide bridges, which contribute to stabilize the reactive site. The reactive loop of KTIs with the P1-P1’ site binds to the serine protease (Bendre et al., 2018; Bonturi et al., 2022; Oliva et al., 2010). Serine proteases, characterized by a Ser/His/Asp catalytic triad, reversibly interact with KTI via hydrogen bonds at this active site (Agback and Agback, 2018; Clemente et al., 2019; Grosse-Holz and van der Hoorn, 2016; Joshi et al., 2013, 2013; Katz et al., 2001; Oliva et al., 2010; Patel, 2017). Several mechanisms exist for how proteinase inhibitors interact with their targets, resulting in bifunctional activities (Grosse-Holz and van der Hoorn, 2016). For example, they block alpha-amylases from barley grains (Abdul-Hussain and Paulsen, 1989) and inhibit proteases from the pathogenic fungus *Fusarium culmorum* (Pekkarinen et al., 2007). KTIs are implicated in various physiological functions such as nitrogen recycling during senescence (Havé et al., 2017) and protein processing during development (Rustgi et al., 2017), in addition to mediating herbivore interactions (Kaling et al., 2018; Major and Constabel, 2008, 2007; McCormick et al., 2016; Philippe et al., 2009; Philippe and Bohlmann, 2007).

The interaction of plants with insects is regulated by phytohormones of the jasmonate (JA) pathway (Erb et al., 2012; Lortzing and Steppuhn, 2016). Oral secretions of insects contain elicitors like volicitin, inceptin, and caeliferins, which can induce JA-specific responses (Alborn et al., 1997; Schäfer et al., 2011; Schmelz et al., 2006). Furthermore, JAs (jasmonate-derivates) are produced in response to wounding (Conconi et al., 1996; Feussner and Wasternack, 2002; Qi et al., 2011; Vick and Zimmerman, 1983). Feeding of gypsy moth on poplar leaves and treatment with JA causes partially overlapping responses, including *KTI* induction (Babst et al., 2009). However, insect feeding also induces other signaling pathways such as ethylene or abscisic acid (ABA) signaling, in parallel with KTIs (Babst et al., 2009; Kaling et al., 2018).

In poplar, *KTI*s form a large gene family with more than 30 members (Bradshaw et al., 1990; Eberl et al., 2021; Guo et al., 2025; Major and Constabel, 2008; Philippe et al., 2009). Transcriptomic analysis revealed that specific *KTIs* of *P. trichocarpa* x *deltoides* show distinct profiles across different tissue types impacted by herbivore attack (*Malacosoma disstria*) (Philippe et al., 2009). It was also observed that *KTI*s in *P. nigra* are transcriptionally regulated in a herbivore-specific manner (Eberl et al., 2021). Given the diversity of the transcriptional responses, it is not surprising that distinct functions of most members of the KTI family in poplar herbivory defence remained experimentally unexplored.

This study aimed to address this gap and elucidate functions of distinct KTIs in poplar. Towards this goal, we selected candidate genes in *P.* x *canescens* by an *in-silico* strategy, based on published reports (Kaling et al., 2018; Kasper et al., 2022) of the response patterns of KTIs to insect feeding and presence in extracellular fluid. This analysis resulted in three candidate KTI genes in *P.* x *canescens* with homology to *Potri.019G124400*, *Potri.019G124600*, and *Potri.017G153200* (hereafter referred to as *KTI_400*, *KTI_600*, and *KTI_53200*, respectively). We then validated their transcriptional responses to wounding and phytohormone exposure. In the next step, we generated gene knock-out mutants using the CRISPR/Cas12a approach. We also produced constitutive over-expression lines under the *p35S* promoter to establish their role in herbivory defense and growth performance. Toward this end, we investigated the phenotypes of the transgenic poplar lines and performed herbivore feeding assays.

## 2. Methods

### 2.1 Poplar propagation and growth conditions

We used hybrid poplar *Populus tremula* x *P. alba* (syn. *Populus* x *canescens,* INRA 717-1B4) for all experiments. We cloned poplar plantlets by microcuttings on half-strength MS medium (Murashige and Skoog, 1962) with vitamins (Duchefa Biochemie B.V., Haarlem, Netherlands) as reported previously (Müller et al., 2013). The cuttings were grown under long day light conditions [16 h light, 70 - 85 µE m^-2^ s^-1^ PAR (photosynthetic active radiation), light source: L18W/840, Osram, Munich, Germany, 60 % RH (relative air humidity) at 24 °C for approximately 4 weeks. We sliced leaves and stems of the plantlets for transformation experiments and used rooted plants for growth, phytohormone treatments and bioassays.

### 2.2 Phylogenetic analysis and selection of candidate genes

For the phylogenetic analysis, putative KTI polypeptide sequences were searched in the *Populus trichocarpa* database PlantGenIE (https://plantgenie.org/; Date accessed: 28^th^ August 2023) and in sPta717 v2 *P.* x *canescens* from AspenDB (https://www.aspendb.org/downloads; Date accessed: 28^th^ August, 2023). Since *P.* x *canescens* (INRA 717-1B4) is a hybrid, gene model searches were performed for both parents, *Populus tremula* and *Populus alba.* Polypeptide sequences of these gene models were predicted by searching the Open Reading Frame (ORF) and their translation to amino acids. The prediction of ORFs and polypeptide translation, followed by multiple sequence alignment of the putative KTIs from *P. trichocarpa, P. tremula* and *P. alba,* were performed in Geneious Prime (Biomatters Ltd., Auckland, New Zealand; version: 2023.12) applying theClustal Omega 1.2.2., mBed algorithm. The Geneious Tree Builder was used for the development of the phylogenetic tree with genetic distance model, Jukes-Cantor, tree build method, unweighted pair group method with arithmetic mean (UPGMA), bootstrap of 1000 replicates, and support threshold of 100 % (Additional Figure S1).

In a previous study, we exposed *P.* x *canescens* WT to poplar leaf beetle (*Chrysomela populi*) in cages under outdoor conditions (Kaling et al., 2018). We downloaded the transcriptome of control and beetle- fed leaves of *P.* x *canescens* from this experiment (Kaling et al., 2018) and searched the putative *KTI*s by their Potri.IDs. We extracted the mean transcript abundances of significantly differentially expressed *KTI* genes and clustered them with ClustVis [(http://biit.ut.ee/custvis/, accessed 20^th^ May 2025, (Metsalu and Vilo, 2015)]. We also searched the xylem sap of *P.* x *canescens* for the presence of KTIs (Kasper et al., 2022). Based on their expression profiles and presence or absence in xylem sap, we selected three candidate genes “Potri.019G124600”, “Potri.019G124400” and “Potri.017G153200”, hereafter called KTI_400, KTI_600 and KTI_5320, respectively.

### 2.3 Prediction of signal peptides

The polypeptide sequences across the haplotypes of *P. tremula* and *P. alba* (Additional Figure S2) were used as a search query for the *in-silico* prediction of signal peptides in KTI_400, KTI_600 and KTI_53200. The databases SignalP-6.0 (https://services.healthtech.dtu.dk/services/SignalP-6.0/; accessed on 29^th^ March, 2025) and PrediSI (http://www.predisi.de/; accessed on 29^th^ March, 2025) were used for the predictions of the signal peptide. WoLF PSORT (https://www.genscript.com/wolf-psort.html; accessed on 29^th^ March, 2025) and Plant-mSubP (https://bioinfo.usu.edu/Plant-mSubP/; accessed on 29^th^ March, 2025) were used for the prediction of the sub-cellular localization.

### 2.4 Transformation of poplar

For the transformation of poplar, we designed vectors, cloned them in *E. coli*, transformed them into *Agrobacteria*, which were then used to transform poplars, adapting protocols from (Bruegmann et al., 2019) and (Amirkhosravi et al., 2025). The details of the adapted pipeline have been described in the Additional “Methods”. Briefly, the CRISPR/Cas12a gene knock-out and cloning strategies were adapted from Merker *et al*., (2020). The Gateway-compatible cloning plasmid sets, pDettLbCas12a and pEnRZ- Lb-Chimera were a gift from Prof. Dr. H. Puchta (KIT, Karlsruhe, Germany) and were used for poplar transformation. For the target site design, the PAM (Protospacer Adjacent Motif) site of the CRISPR- Cas12a system (5’-TTTV-3’) was initially searched within the exon regions of *KTI_400, KTI_600,* and *KTI_53200.* Subsequently, 24-nucleotide Target_400_600 and Target_53200 sites, lying at the 3’ site of the PAM were used for the double (*KTI_400* and *KTI_600*) and single (*KTI_53200*) knock-out sites. To ensure the specificity of the target sites, the genome of *P.* x *canescens* (version 2, *P.* x *canescens*; https://www.aspendb.org/downloads; Date accessed: 4^th^ September 2023) was searched to exclude off-targets. We also generated empty vector control lines comprising only the ubiquitin (*ubq)* promoter.

Over-expression lines of the candidate *KTIs* were generated under the *p35S* promoter, using a binary vector set pDONR201 (entry vector; Invitrogen Life technologies) and pK7WG2 (destination vector; Karimi *et al*., 2002), which are Gateway compatible. The CDSs of *KTI_400, KTI_600* and *KTI_53200* were individually cloned into the plasmid vector sets (Additional Methods). We also produced empty vector lines containing only the *p35S* promoter in the destination vector pK7WG2.

For plant transformation, we used excised, slit leaves and stems from three-week old sterile poplar plantlets (see 2.1), which were co-cultivated with *Agrobacterium tumefaciens* (GV3101) containing the desired gene construct. Calli of the co-cultured tissues were induced on a callus-inducing medium Additionaled with gentamicin (60 mg L^-1^) for CRISPR-Cas12a or kanamycin (50 mg L^-1^) for the *p35S* transformed poplars in climatized cabinets (AR-75L, Percival Scientific) at 28°C, 20 µE m^-2^ s^-1^ PAR, 60 % RH, 16 hr of light (light source: Alto 32 Watt, Philips, Amsterdam, Netherlands). The emerging shoots were transferred to a rooting medium containing either gentamicin (60 mg L^-1^) or kanamycin (50 mg L^-1^) and grown at 28 °C, 60 µE PAR with 16 hr light for 4 weeks (light source: Alto 32 Watt and LG4507.4).

To control the insertion of T-DNA and for genotypic analyses of the mutant poplar lines, DNA was extracted from 100 mg of frozen, milled leaf tissues using the innuPREP Plant DNA kit (Analytik Jena GmbH, Jena, Germany) according to the manufacturer’s instructions and analysed by PCR (see Additional “Methods”, Primers for cloning and the PCRs are presented in Additional Table S1). The gene editing patterns introduced due to the CRISPR/Cas12a systems and the insertion of the overexpression constructs were confirmed via Sanger sequencing (Microsynth seqlab, Göttingen, Germany) of the PCR amplicons, flanking the target sequence.

### 2.5 Real Time Quantitative Polymerase Chain Reaction (RT qPCR)

Frozen leaves (100 mg) were milled to a homogeneous frozen powder (MM400, Retsch GmbH, Haan, Germany) under cooling to prevent thawing. We used the innuPREP Plant RNA kit (Analytik Jena GmbH) for RNA extraction according to the manufactureŕs instructions. The resulting RNA was eluted with RNase-free water in a volume of 30µL. Total RNA yield was measured spectrophotometrically using NanoDrop^TM^ One spectrophotometer (Thermo Fisher Scientific, Wilmington, Delaware, US). One µg of RNA was used to synthesize cDNA with RevertAid First Strand cDNA Synthesis kit (Thermo Fisher Scientific), as described by the manufacturer’s protocol.

The transcript level quantification was performed using RT-qPCR with innuMIX qPCR DSGreen Standard (Analytik Jena GmbH), following the manufacturer’s guidelines with the following PCR conditions: initial denaturation at 95°C for 2 min, denaturation at 95°C for 10 sec, annealing temperature specific to the primer (Additional Table S2 for annealing temperature of primer pair) for 10 sec, elongation at 72°C for 20 sec (fluorescence read at this step), 45 cycles of denaturation to elongation. The transcript abundances were analyzed with the ddCt method (as described by Pfaffl, 2001) with the qSOFT program (version: 4.0, Analytik Jena GmbH). Two house-keeping genes *ACTIN* and *UBIQUITIN* were used for the normalization. All primer pairs were tested for their efficiency (Pfaffl, 2001). The primers are specified in Additional Table S2.

### 2.6 Wounding experiments

Rooted WT poplars (see 2.1) were potted in “N-type soil” (Hawita Gruppe GmbH, Vechta, Germany) and acclimated to greenhouse conditions (16 h light, 21 to 28 °C, 150 µE m^-2^ s^-1^ light conditions, approximately 30 % relative air humidity, light source: 163 15L34, Adolf Schuch, Worms, Germany) as described by Müller et al. (2013). Plants were irrigated with tap water every alternate day and were randomized weekly. For foliar mechanical wounding experiments, the third fully developed leaf from the top of eight-week-old poplars was used. Wounding was performed with a micro-tissue tweezer, 1 x 2 teeth (Prestige 7-102) by punching 15 to 20 holes, distributed homogenously on the leaf. After three and eight hours, the wounded leaf was excised and flash-frozen in liquid nitrogen. Control samples were the third fully developed unwounded leaf from the top of independent plants (n = 3 to 4 individual plants per treatment). To avoid bias by stress volatiles, wounded and control plants were separated in independent greenhouse cabinets for the experiments. The leaves were used to determine the transcript abundances of target genes as described above (see 2.5).

### 2.7 Phytohormone treatments

For exogenous phytohormone application, ten-week-old greenhouse-grown WT poplars were divided into separate greenhouse cabinets and exposed to one of the following treatments: meJA (methyl jasmonic acid, 200 µM dissolved in demineralized water, Merck KGaA, Darmstadt, Germany), ACC (aminocyclopropane-1-carboxylic acid, 100 µM, dissolved in demineralized water; Merck KGaA), meJA- and ACC-mock solution (demineralized water), BTH (Benzothiadiazole 1000 µM, dissolved in 10 % methanol; Merck KGaA) and BTH mock solution (10 % methanol). All phytohormone and mock solutions were Additionaled with 0.1 % Tween-20 (Merck KGaA) to improve adhesion to the leaves. The plants were individually sprayed on the leaves’ abaxial and adaxial surfaces until drip-off. After spraying, plants were immediately enclosed in a polypropylene bag (400 x 780 mm, Labsolute, Th. Geyer GmbH and Co. KG., Höxter, Germany) for 4 h. Subsequently, the plants were grown for an additional 4 h and 20 h without polypropylene bags before tissue sampling. This resulted in sampling time points of 8 h and 24 h. The first fully developed leaf from the top was harvested, shock-frozen in liquid nitrogen and used to determine the transcript abundances of target genes as described above (see 2.5). Each treatment was conducted with n = 3 to 4 individual plants.

### 2.8 Physiological and phenotypic characterization of transgenic poplars

Rooted plants of the WT, the CRISPR/Cas12a knock-out lines (= *kti* lines), the *p35S KTI* overexpressing lines (*KTIox* lines) and empty vector control lines for *KTIox* and *kti* were potted and acclimated to greenhouse conditions (20 – 28 °C, 150 µE m^-2^ s^-1^ PAR, 16 hr light supplied as ambient light Additionaled with 163 15L34, Adolf Schuch, Worms, Germany; approximately 30 % relative air humidity) as described above (see 2.6). The plants were daily irrigated with tap water and grown for two months. Gas exchange was determined four times in bi-weekly intervals on the third fully expanded leaf with a portable photosynthesis system device (LCpro+, ADC BioScientific Ltd, UK) under ambient light conditions. At harvest, leaves, stems and roots of all plants were weighed, dried (2 weeks at 60 °C) and used to determine whole plant biomass. We used 2 or 3 independent transgenic lines per transformation event (empty vector controls, the *KTIox* and the *kti* lines), each with four plants and the WT (n = 8).

### 2.9 Bioassay of transgenic poplars with *Helicoverpa armigera*

The eggs of the broad-range generalist insect *Helicoverpa armigera* (provided by Prof. Dr. M. Rostás, Agricultural Entomology, Department for Crop Sciences, University of Göttingen) were allowed to hatch in a plastic container (8 cm length × 13.5 cm width × 6 cm height) at 22°C and 16 hr light [60 μE m^−2^ s^−1^ PAR (provided by Alto 32 Watt, Philips, Amsterdam, Netherlands)], 60 % RH in a climate cabinet (AR-75L, Percival Scientific, Perry, USA), containing an artificial diet composed of alfalfa powder, rapeseed oil, baker’s yeast, Wesson Salt Mix, β-sistosterol, L-leucine, ascorbic acid, vitamin mix, sorbic acid, bean flour and 4-hydroxybenzene S (Gergs and Baden, 2021). The larvae were used when they reached their 1^st^ to 2^nd^ instar stage (determined after (Mironidis and Savopoulou-Soultani, 2008) and had lengths of approximately 1 to 3 mm. The starting weight was determined for pools of 10 larvae on an analytical balance (Cubis® MCA225S-2S00-I, Sartorius, Göttingen, Germany) and divided by 10.

Two- to 3-week-old rooted WT and transgenic poplar plants from stock cultures (see 2.1) were individually placed on solid ½ MS media in sterile squared Magenta jars (size: 76 x 76 x 102 mm Magenta, Merck KGaA). The plants were cultivated for one week in climatized cabinets (Percival Scientific, Perry, USA; 22°C, 60 μE m^−2^ s^−1^ PAR, 16 hr light, light source: Alto 32 Watt). A single *H. armigera* larvae was placed in each Magenta jar containing one poplar plantlet. After 12 days of feeding, the weight of each individual *H. armigera* was measured on the analytical balance (Cubis® MCA225S-2S00-I, Sartorius). Weight gain was determined as (Weight after feeding - Weight at the beginning). Two independent experiments were conducted, each included WT, empty vector controls, and 2 lines per *kti* and *KTIox*. The number of individual plants per experiment varied and is indicated in the figure legend.

### 2.10 Statistical analyses

All statistical analyses were performed in R (R Core Team, 2022) and were visualized in RStudio (RStudio Team, 2020) or using OriginPro2024b (Northampton, Massachusetts, USA). Data were checked for normal distribution and variance homogeneity (Levenés test, visual inspection of residuals). ANOVA was performed using general linear models with the package “multcomp” (Hothorn et al., 2008) followed by a post-hoc test (usually Tukey). When the data were not normal-distributed, we used the Kruskal-Wallis test for pairwise comparisons. When more than one experiment was analyzed, packages “car” (Fox and Weisberg, 2019) and “lme4” (Bates et al., 2015) were used to assign random effects to every experiment. Differences between means of treatments and controls were considered significant at *p* < 0.05.

## 3. Results

### 3.1 *In-silico* selection of potential candidate KTIs

We conducted multiple sequence alignment of the amino acid sequences inferred from the *P.* x *canescens* genome (i.e., the parent’s genomes *P. alba* and *P. tremula*) together with the *P. trichocarpa* genome and identified a total of 28, respectively 29 *KTI* sequences in the haploid parent genomes of *P.* x *canescens* (Additional Fig. S1). In this study, we used the Potri IDs of the closest *P.* x *canescens* homologs for annotation to ease the comparison among studies. The Potra IDs are shown in Additional Table S3.

The putative KTIs clustered in three large clades with 8 to 10 members (Additional Fig. S1). Two KTIs (Potri.019G088200 for both parents and Potri.003G097900 only for *P. alba*) did not cluster with any of the larger groups (Additional Fig. S1).

Twenty-two of the *KTI*s in *P.* x *canescens* showed significant upregulation of transcript abundances in response to poplar leaf beetle feeding (Fig. 1). Genes with the greatest increases in transcript abundances in response to herbivory were KTI_400 and KTI_600 (both in clade I of the phylogeny, orthologue to TI3 in *P. trichocarpa* x *deltoides*, Major & Constabel, 2008). Further relatively strong transcriptional responses were observed for a group of five genes, including members of clade I and clade II in addition to the single *KTI,* Potri.019G088200 (Fig. 1). *KTI,* Potri.019G088200 is an orthologue to *P. nigra* PnD1 (Eberl et al., 2021). The remaining genes showed moderate or low transcriptional regulation and comprised members of clade I, II, and III of the phylogeny (Fig. 1).

**Figure 1:**
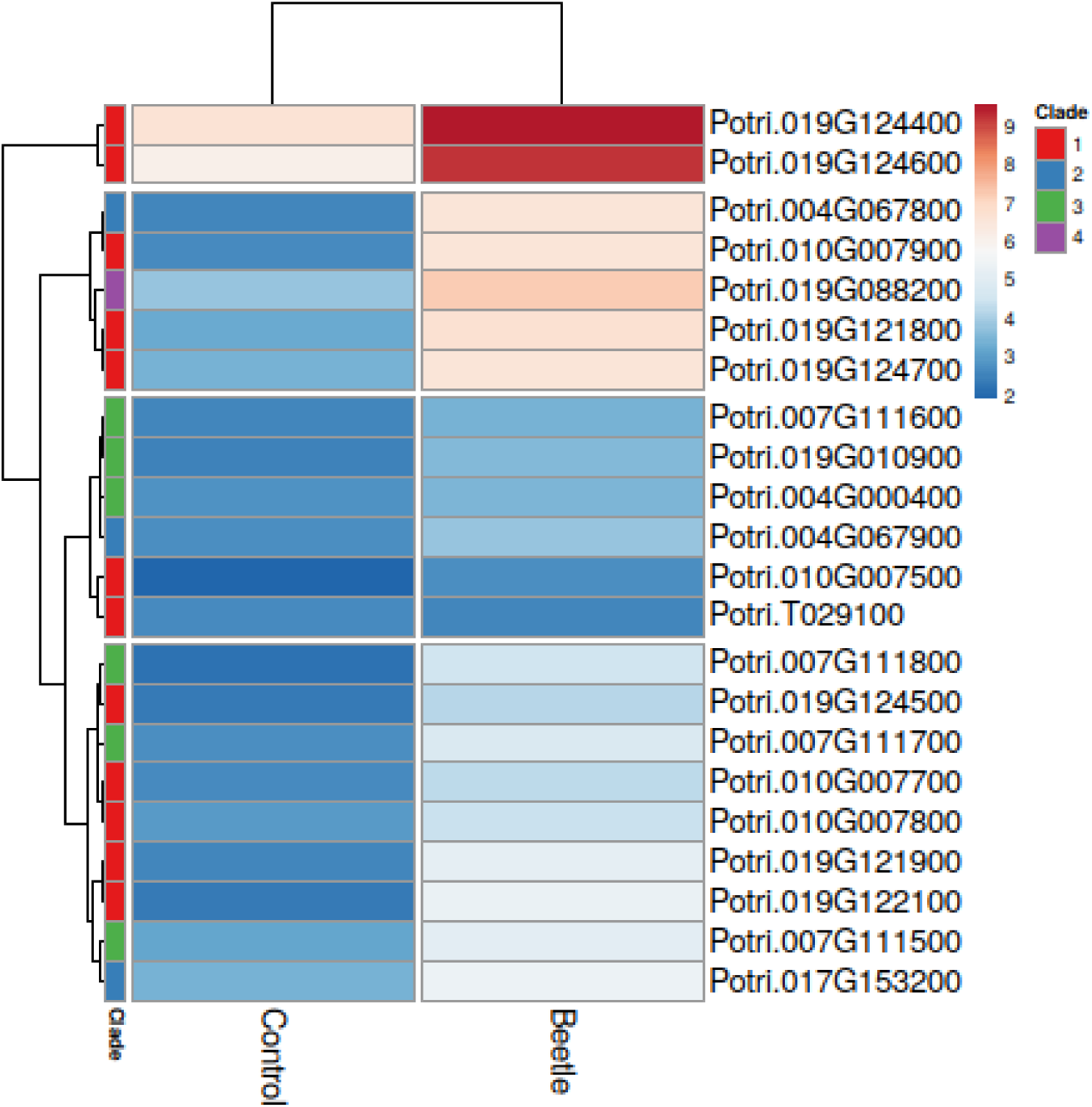
Differentially expressed *KTI* genes (*p_adj_* < 0.05) after exposure of *P*. x *canescens* to *C. populi* in outdoor cage areas. Data were extracted from the Additionalal material of RNAseq results in Kaling et al. (2018). Means were clustered in rows. Left panel “clade” indicates the phylogenetic clade of the genes.

For further analyses, we selected *KTI_400* and *KTI_600* (clade I) because of their massive response to herbivory. Furthermore, *KTI_53200* (clade II, orthologue to *KTI5* in *P. alba,* Bradshaw *et al*., 1990) with a moderate herbivory response (Fig. 1) was chosen because the protein was present in the xylem sap of *P.* x *canescens*, where it might play a role in poplar immune responses, while KTI_400 and KTI_600 proteins were not present this compartment (Kasper et al., 2022).

### 3.2 Molecular features of potential candidate KTIs

Among the selected candidates, KTI_400 and KTI_600 exhibited 92% identity of the amino acid sequences and both had approximately 70% identity with their closest neighbours Potri.019G121900, Potri.019G124500, Potri.019G124700 and Potri.019G122100 (Additional Fig. S1, Fig. S2). With the exception of Potri.019G124700, the close neighbours showed only very low responsiveness to herbivory (Fig. 1). KTI_53200 shared 67% amino acid sequence identity with its closest neighbour Potri.004G067900, a gene also showing only low induction by herbivory (Fig. 1). Owing to a relatively large dissimilarity within the amino acid sequences, KTI_53200 shared only 19% identity with KTI_400 and 36% with KTI_600 (Additional Fig. S2). KTI_400 and KTI_600 showed 47% amino acid identity with their closest *Arabidopsis thaliana* ortholog At*KTI3* (At1g73325) and KTI_53200 51% with At*KTI5* (At1g17860).

Each of the three candidate KTIs, KTI_400, KTI_600 and KTI_53200 was characterized by a signal peptide (Fig. 2, Additional Fig. 2 for the amino acid sequence), which predicted extracellular localization with high probability (Additional Table S4). However, a note of caution is warranted because no consistent prediction for subcellular localization was observed when other prediction programs were used (Additional Table S4). Furthermore, the candidate KTIs contained the characteristic Kunitz motif ([L, I, V, M]-X-D-X2- G-X2-[L, I, V, M]-X5-Y-X-[L, I, V, M]) and six cysteine residues, of which four are predicted to form disulphide bridges (Fig. 2, Additional Fig. 2). The P1-P1’ motif in the reactive sites, which interact with the proteases, were “E-S” (glutamic acid-serine) residues at 86-87 aa position of KTI_400 and KTI_600, and “D-D” (aspartic acid-aspartic acid) residues at 89 aa position of KTI_53200 (Additional Fig. S2).

**Figure 2:**
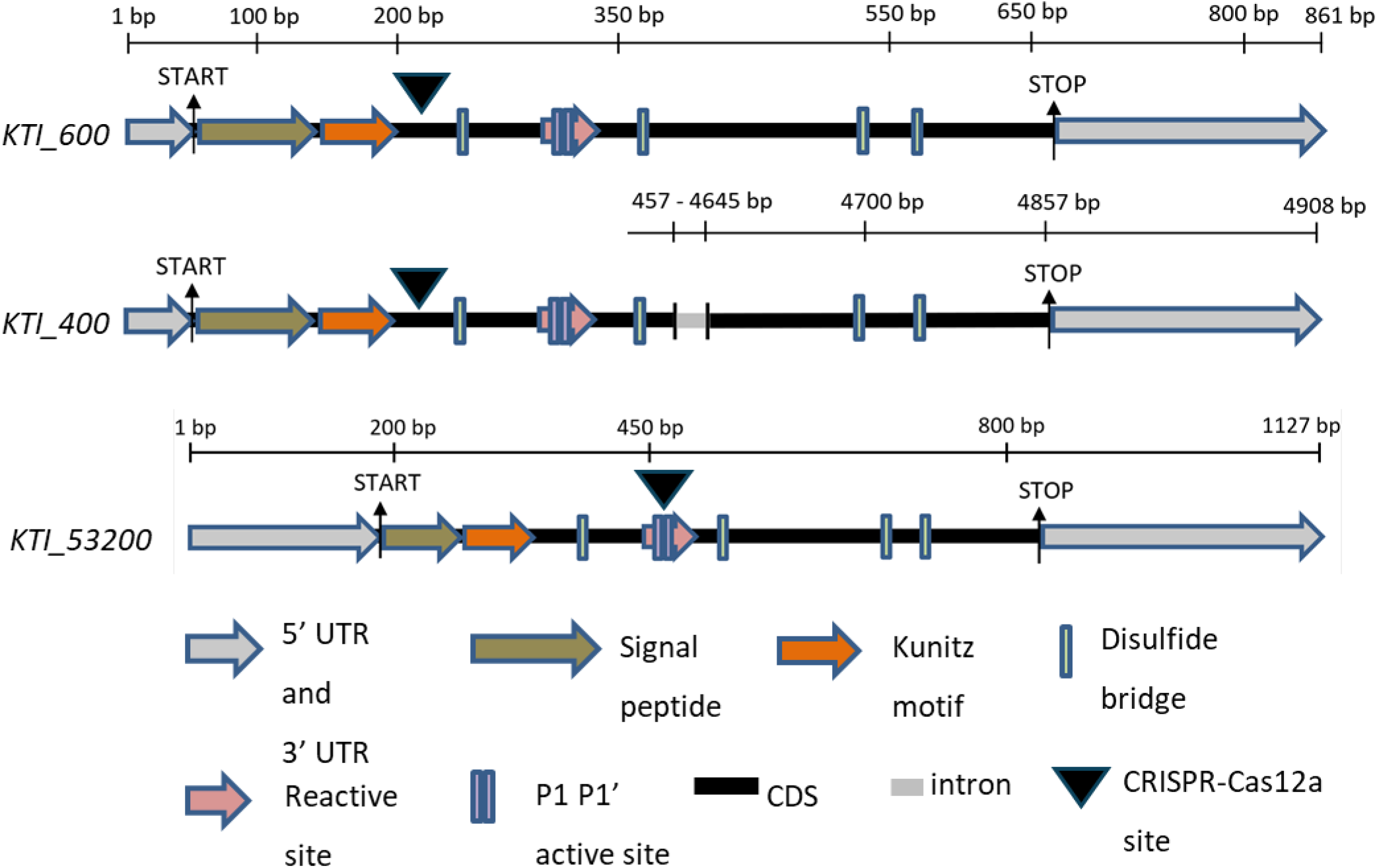
Gene models of candidate *KTI_400*, *KTI_600* and *KTI_53200*. Motifs of Kunitz Trypsin Inhibitor marked with colored arrows and boxes. The length of genomic DNA is stated above the respective sequence. The cDNA/amino acid lengths are *KTI_400:* 932bp/203 aa, *KTI_600:* 861bp/203aa and *KTI_53200:* 1122bp/ 210aa. Start and stop codons are marked with black arrows. All sequences have been retrieved from sPta717 v2 (https://www.aspendb.org/databases; date accessed: 21^st^ August, 2020).

### 3.3 *In-planta* response to wounding and phytohormone treatments of the candidate *KTI*s

The candidate *KTI*s showed significant increases in transcript abundances 3h and 8h after mechanical wounding of leaves (Fig. 3a). The initial wounding response was greatest for *KTI_400* with up to 200-fold increases in transcript levels but declined afterwards, whereas *KTI_600* was less induced (approximately 40-fold) but remained stably increased after 8h (Fig. 3a). *KTI_53200* showed less responsiveness to wounding than *KTI_400* and *KTI_600* (Fig. 3a).

**Figure 3:**
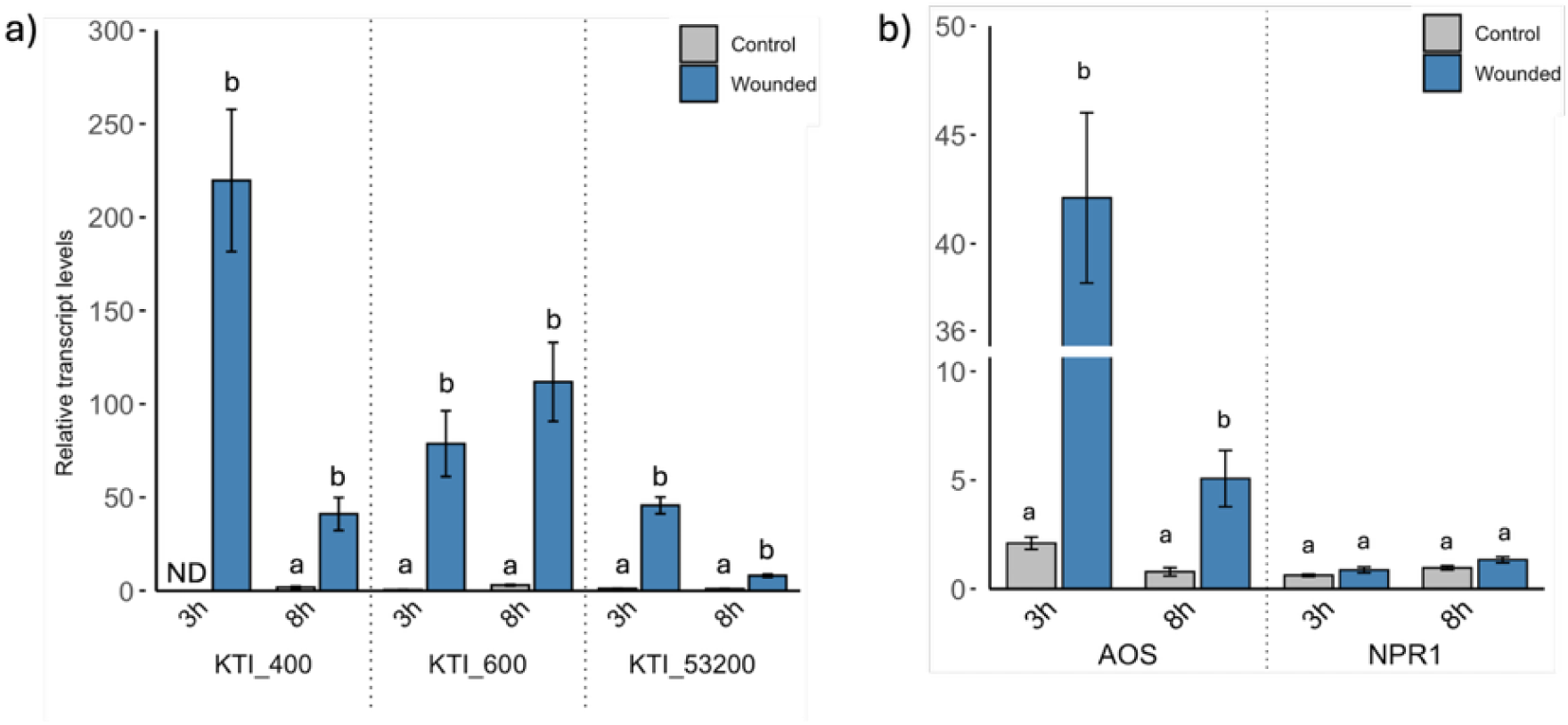
Transcript abundances of candidate *Kunitz Trypsin Inhibitors,* jasmonic acid, and salicylic acid marker genes in response to wounding of *Populus* x *canescens* leaves. Relative transcript abundances are shown of a) *KTI_400, KTI_600, KTI_53200* and b) *AOS* and *NPR1* at 3 and 8 hr post wounding. The third leaf of 8-week-old poplars grown in soil in the greenhouse was used for wounding and RT-qPCR. Leaves from non-wounded plants sampled at the same time points were used as controls. Transcript abundances of the target genes were expressed relative to the reference genes *ACTIN* and *UBIQUITIN*. Bars show means of n = 3 to 4 ± SD biological replicates. Each biological replicate is an independent plant with its third leaf from the stem apex wounded. Different letters in the panel represent statistical significance calculated between the control and wounded tissues at an individual time-point using ANOVA with *p* < 0.05 (Tukey post-hoc test). ND, non-detectable.

Since wounding responses may imply regulation by phytohormones (Erb et al., 2012), we studied marker genes for the JA and SA pathway, *ALLENE OXIDE SYNTHASE* (*AOS*) and *NON-EXPRESSOR OF PATHOGENESIS-RELATED GENES 1* (*NPR1*), respectively. We observed divergent responses of *AOS* and *NPR1* to wounding (Fig. 3b). *AOS* transcript abundances were increased after 3h and 8h, similar to the pattern observed for *KTI_400*, whereas *NPR1* was unaffected (Fig. 3b).

To obtain further evidence for phytohormone responsiveness of the candidate KTIs, we treated *P*. x *canescens* exogenously with meJA (JAs), BTH (SA analogue) or ACC (ethylene precursor). *KTI_400* and *KTI_600* were affected by meJA treatment, while *KTI_53200* did not show significant increases in transcript abundances (Fig. 4a). The highest up-regulation was noted for *KTI_600* with approximately 10- fold and 35-fold increases in expression levels at 8h and 24h post meJA treatment. *KTI_400* showed significantly increased transcript levels (5-fold) at 8h post meJA treatment and declined to the levels of the mock-treated controls at 24h. BTH and ACC exposure of poplar did not cause significant effects on the candidate *KTIs* (Fig. 4 b,c).

**Figure 4:**
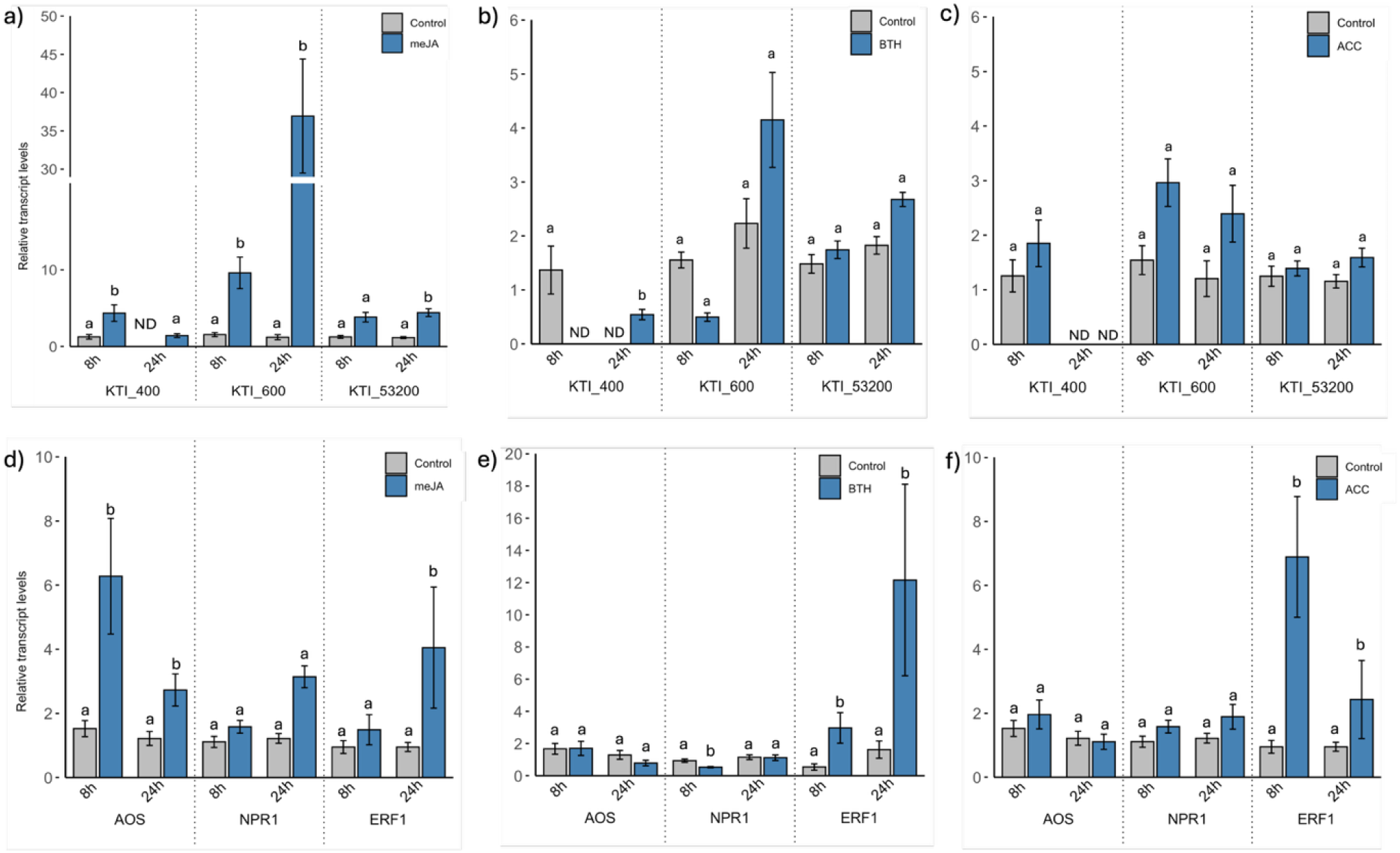
Response of the candidate *Kunitz Trypsin Inhibitors*, jasmonic acid, salicylic acid and ethylene marker genes 8h and 24h after exogenous phytohormone spraying. a,d) meJA (200 uM); b,e) BTH (1000 µM); c,f) ACC (100 µM) were sprayed to *Populus* x *canescens*. Bars represent the mean transcript abundance of *KTI_400, KTI_600, KTI_53200, AOS,* and *NPR1* normalized to the reference genes *ACTIN, UBIQUITIN,* and mock-treated controls, n = 3 or 4 (biological replicates). Plants were separated in different chambers, and meJA, BTH, and mock solution were sprayed on the leaves until dripping. Transcript abundance was quantified via RT-qPCR. Error bar represents the standard deviation. Different letters in the panel represent statistical significance calculated using ANOVA with *p* < 0.05 (Tukey post-hoc test) between control and treatment. ND, non-detectable. meJA, methyl-jasmonic acid; BTH, Benzothiadiazole; ACC, 1-Aminocyclopropane-1-carboxylic acid.

Initially, we also included *KTI*_Potri.019G088200 in our experiments. *KTI*_Potri.019G088200 did not show a rapid wounding response, like our candidate *KTI*s (Additional Fig. S3). However, *KTI*_Potri.019G088200 responded to meJA treatments with increased transcript levels (significant only after 8h) and was not induced by ACC or BTH (Additional Fig. S3).

We further tested marker gene expression for the exogenously applied phytohormones to validate their *in planta* activity (Fig. 4d-f). *AOS* transcript levels were responsive to meJA at both tested time points, showing approximately 6- and 2-fold increases after 8h and 24h, respectively (Figure 4d). For *NPR1,* a significant decrease at 8h post BTH exposure was observed, whereas AOS was unresponsive to BTH and ACC treatments (Fig. 4e). ACC caused an increase in *ETHYLENE RESPONSIVE FACTOR 1* (*ERF1*) transcript abundance (Fig. 4f).

### 3.4 Phenotypes of poplar overexpression and knock-out *KTI* lines

For poplar transformation, we chose a double knock-out strategy, simultaneously targeting *KTI_400* and *KTI_600* (assigned as *kti4+600*) and a single knock-out strategy for *KTI_53200* (*kti_53200*), using CRISPR/Cas12a. Positive CRISPR-Cas12a transgenic lines were tested by PCR (see 2.4), followed by Sanger sequencing of the CRISPR-Cas12a target region. The CRISPR-Cas12a construct targeting *KTI4+600* showed gene-editing patterns of 50, 14, 11, 10, 9, 8, 6 and 2 bp deletions, whereas the CRISPR- Cas12a construct targeting *KTI_53200,* showed deletions of 10, 8, 7, 6 and 5 bp (relevant mutant lines in Additional Table S5). The CRISPR-Cas12a mutant lines *kti4+600_1_28*, *kti4+600_1_52* and *kti53200_2_18*, and *kti53200_2_29* were chosen for further functional characterization due to their prominent frame-shift mutations and their putative pre-mature stop-codon (Additional Table S6).

Overexpression lines (*KTIox*) were generated by expression of each *KTI* individually under the *p35S* promoter. The transcript levels of the target candidate *KTI*s were quantified in 26 *KTIox* lines (four *KTIox_400* lines, eight *KTIox_600* and fourteen *KTIox_53200*, Additional Fig. S4). Compared with WT levels, elevated *KTI* expression levels ranged from 10- to 60-fold in most of the tested *KTIox_400* and *KTIox_600* lines (Additional Fig. S4a,b). The *KTIox_53200* lines showed very high overexpression levels ranging from approximately 500 to 12000 above the WT levels (Additional Figure S4c). We selected *KTIox_400_4_14*, *KTIox_400_4_15*, *KTIox_600_5_9*, *KTIox_600_5_9, KTIox_53200_6_9* and *KTIox_53200_6_15* for further analyses.

We cultivated WT, empty vector lines, *KTIox*, and *kti* lines under greenhouse conditions (Additional Fig. S5). There was no obvious visual phenotype (Additional Fig. S5). We did not observe significant differences in photosynthesis (Fig. 5a). Other physiological parameters (stomatal conductance, transpiration) and morphological parameters (Plant height, stem diameter, biomass of leaves, stem and roots, root-to-shoot ratio) either showed no or small but significant variations among the lines (Additional Table S7). However, the whole-plant biomass (sum of leaf, stem and root) of the *kti4+600* lines was significantly greater than that of the other lines (Fig. 5b).

**Figure 5:**
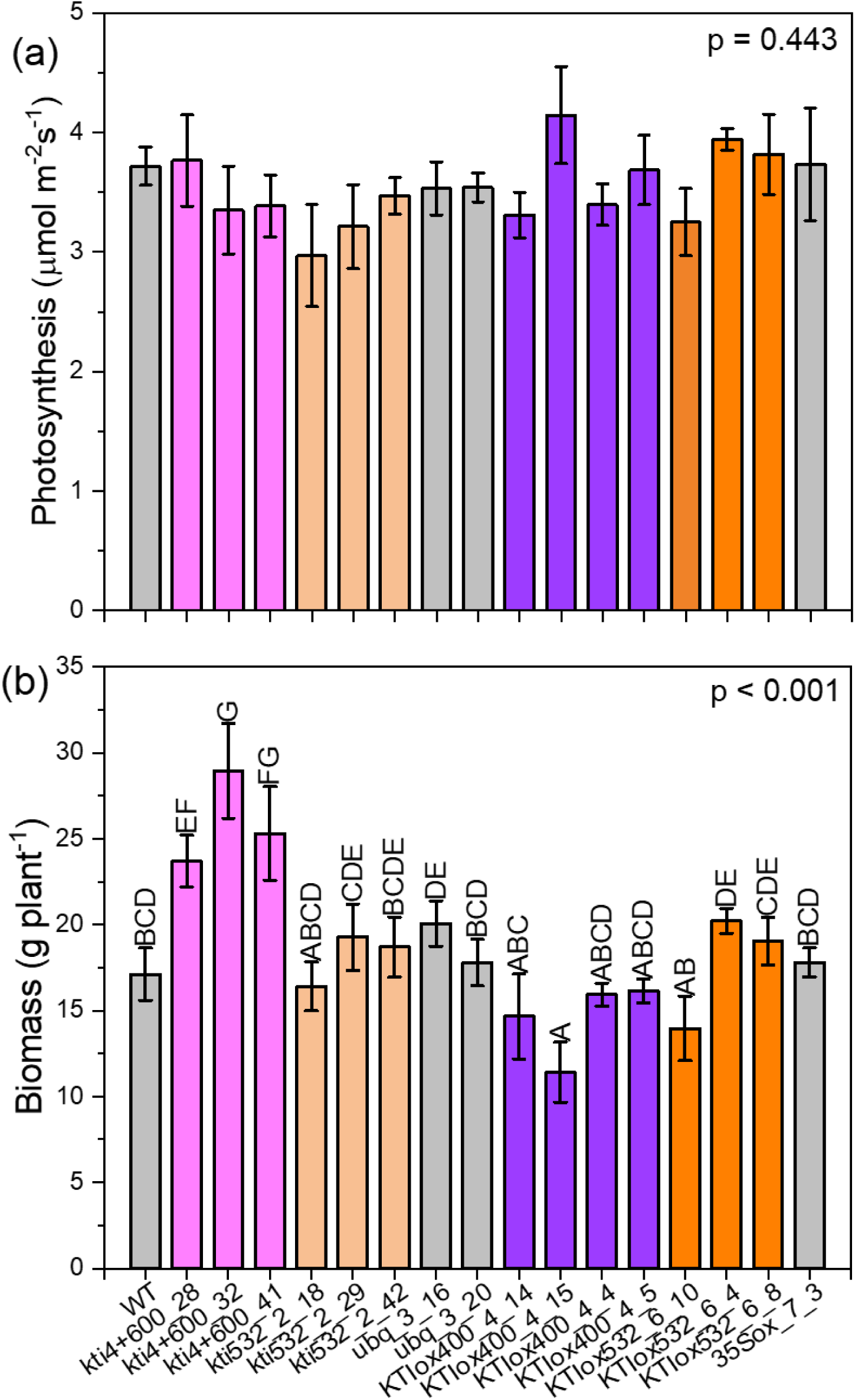
Photosynthesis and biomass of poplar WT and transgenic lines. The plants were grown for 2 months in greenhouse cabinets. n = 4 plants per line, WT: n = 8. Data show means ± SE. Different letters indicate significant differences of means at *p* < 0.05 (Tukey test).

### 3.5 Differences in herbivory on *KTI* overexpressing and knock out poplar lines

Initial attempts to grow the generalist *H. armigera* on *P.* x *canescens* leaves of greenhouse-cultivated plants as the sole diet were not successful due to high mortality of the larvae, whereas larvae on an artificial diet showed massive weight gain from approximately 0.4 mg to approximately 19 mg within 12d. After testing various conditions, we found that exposing sterile-grown, “naive” poplar plantlets individually in Magenta jars to *H. armigera* resulted in reproducible results. Significant damage by herbivory was visually noted for all lines (Additional Fig. S6).

*H. armigera* larvae revealed significantly higher weight gain after feeding on *kti4+600_1_28* and *kti4+600_1_52* than on WT or on empty vector plants *ubq::3_16,* and *ubq::3_20* (Fig. 6). The weight gain of the larvae feeding on these mutant lines was approximately three-fold higher than on WT plants. In contrast to the *kti4+600* poplars, there were no significant differences among *kti_53200* lines to the WT or empty vector lines (Fig. 6).

**Figure 6:**
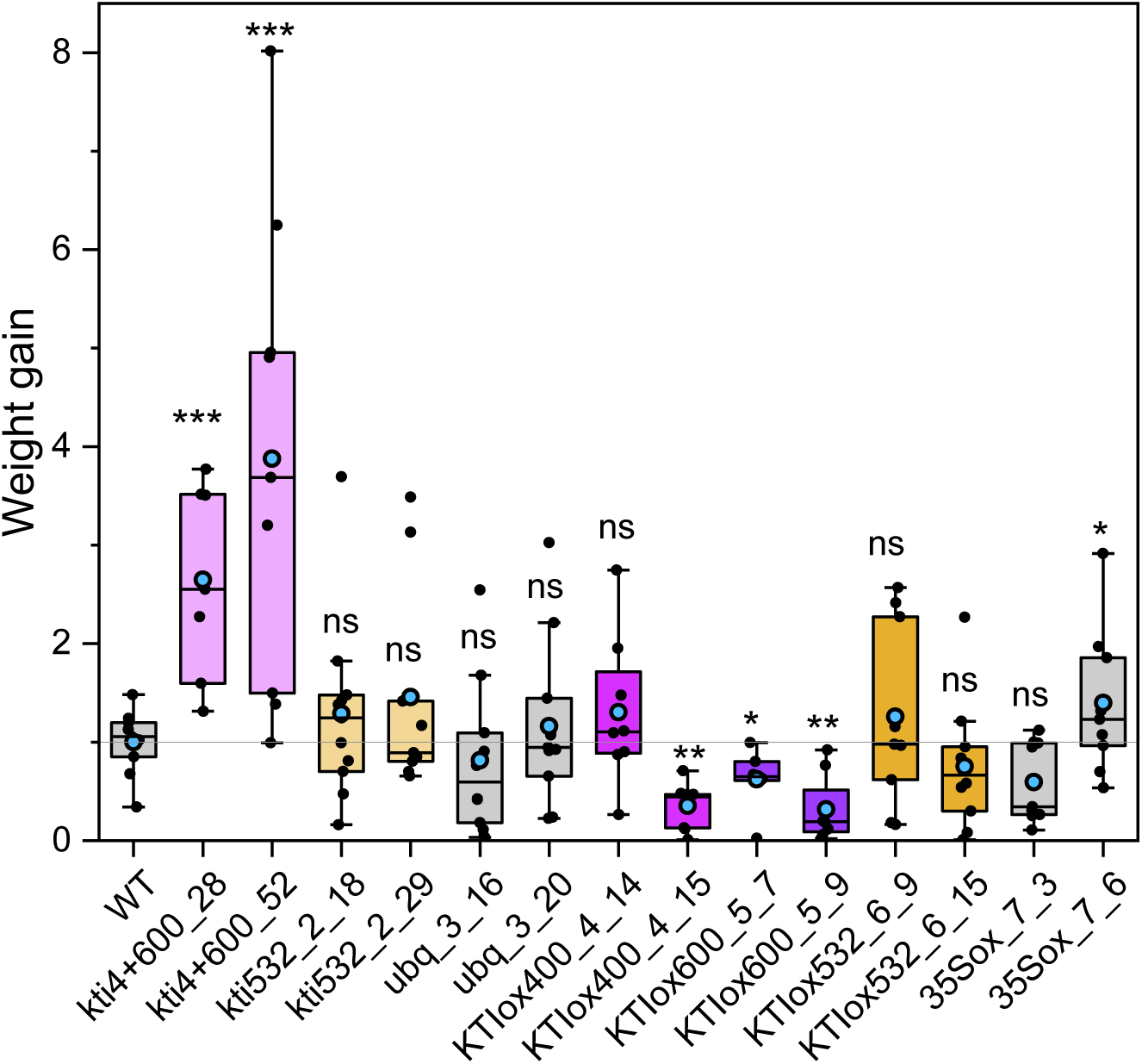
Weight gain of *Helicoverpa armigera* larvae on *Kunitz Trypsin Inhibitor* transgenic poplars. One II-instar stage larva was applied per plant to CRISPR-Cas12a loss-of-function *kti4+600_1_28*, *kti4+600_1_52*, *kti53200_2_18*, *kti53200_2_29*, empty control vector *ubq*::*3_16* and *ubq*::*3_20* lines; *35S* over-expressing *KTI400_4_14*, *KTI400_4_15*, *KTI600_5_9*, *KTI600_5_9, KTI53200_6_9* and *KTI53200_6_15,* empty vector control *35S::7_3* and *35S::7_6* and WT (Wild-type) growing individually in sterile jars. Six to 12 biological replicates per line from two separate experiments replicates were used. The sterile jars containing one poplar plantlet and one larva were placed in growth cabinets for 12 days. In each experiment, the data were normalized to the mean weight gain of the wildtype. Individual data = black circles, horizontal line = median, blue circle = mean. Box extends from the 25^th^ to 75^th^ percentile, and whiskers encompass the minimum and maximum values. Statistical analysis was performed with ANOVA and Kruskal-Wallis post hoc test). Stars indicate significant differences compared to the controls: * p < 0.05, ** p < 0.01, ns = not significant.

In the *p35S-*overexpression lines, we observed about 40% less weight gain for larvae feeding on *KTIox_600_5_7* and about 70% less weight gain for larvae feeding on *KTIox_400_4_15* and *KTIox_600_5_9* but no difference to the WT for *KTIox400_4_14* (Fig. 6). No significant weight differences were observed for *H. armigera* larvae feeding on *KTIox_53200* lines compared with the WT or empty vector lines (Fig. 6).

## 4. Discussion

### 4.1 Phylogeny and expression patterns of poplar *KTI*s

In this study, we determined the phylogeny of KTIs in *P*. x *canescens* based on full-length amino sequences, used an *in-sili*co strategy to identify *KTI*s responsive to herbivore feeding, tested the selected genes for their induction by wounding and phytohormones and used them to dissect their functions as herbivore protectant by forward and reverse genetic engineering. We identified 28 *KTI*s in the haploid genome of *P.* x *canescens,* which compares well with the number of KTI genes reported for other *Populus* species or hybrids [31 KTIs in *P. trichocarpa* x *deltoides* (Philippe et al., 2009), 31 KTIs in *P. trichocarpa* (Ma et al., 2011), 32 KTIs in *P. nigra* (Eberl et al., 2021), 29 KTIs in *P. yunnanensis* (Guo et al., 2025)] and underpins massive expansion of this gene family compared with other species [e.g., seven KTIs in the herbaceous model plant *Arabidopsis thaliana* (Arnaiz et al., 2018), three KTIs in *Camellia sinensis* (Zhu et al., 2019), and four in *Vitis vinifera* (Guo et al., 2025)]. In line with previous assessments (Bendre et al., 2018), we observed the typical structural characteristics for the KTIs (signal peptide, P1-P1’site, cysteine residues, Kunitz motif) in *P.* x *canescens* and confirmed that the KTIs clustered in three major clades and a small group IV, with only one member (Eberl et al., 2021; Guo et al., 2025; Ma et al., 2011).

Several studies suggested that poplar *KTI* genes in clade I are involved in herbivory defence (Constabel and L. Lindroth, 2010; Eberl et al., 2021; Major and Constabel, 2008), that clade II is mixed containing KTIs for defence and development [female catkins, apical shoot, flowers (Clemente et al., 2019; Ma et al., 2011; Major and Constabel, 2008)], whereas functions of KTIs in clade III remain enigmatic. We observed transcriptional responses to *C. populi* feeding for KTIs in all clades; however, distinct members in clade I showed stronger transcriptional induction than those from clade II and III. *KTI_8820* (clade IV) also showed notable upregulation under herbivory by *C. populi* (our study) and other lepidopterans (Eberl et al., 2021) but we observed less massive meJA and wounding responses than for *KTI_400* or *KTI_600*.

Whether genes with low basal transcript levels and low induction found in our study for many clade II and III genes contribute markedly to poplar defense is questionable since the production of KTI proteins is transcriptionally regulated (Ma et al., 2011). Accordingly, increased *KTI* expression levels correlate with higher *in-vitro* trypsin inhibitor activity (Eberl et al., 2021).

Phytohormones, especially JAs, are important for sensing and inducing defences against herbivores (Ali et al., 2024; Erb et al., 2012). The formation of JAs is induced by elicitors and triggered by wounding, starting with lipid oxidation by lipoxygenases (LOX) (Feussner and Wasternack, 2002). Oxidized lipid precursors are then transformed by AOS into the unstable intermediate 12,13-epoxyoctadecatrienoic acid, which marks the entry into the JAs biosynthetic pathway (Feussner and Wasternack, 2002). In poplar leaves damaged by *C. populi*, increased levels of octadecatrienoic acid were observed, implying enhanced JA signaling (Sivaprakasam Padmanaban et al., 2022). In line with previous studies in other poplar species (Christopher et al., 2004; Haruta et al., 2001; Major and Constabel, 2008, 2006; Philippe et al., 2009), we observed that wounding and meJA caused increased transcript levels of *KTI*s. The response intensities and time courses of the selected *KTI*s varied in our study, wounding stimulating *KTI*s in the order *KTI_400* > *KTI_600* > *KTI_53200,* whereas *KTI_600* showing greater induction by meJA than *KTI_400* and *KTI_53200.* In parallel, under both conditions wounding and meJA exposure *AOS*, a key enzyme for JA biosynthesis, was also induced, whereas BTH or ACC exposure did neither influence *AOS* nor our candidatés *KTI* expression. At first glance, the induction of *AOS* by meJA is surprising; however, JAs can increase their own biosynthesis by a feedback mechanism; this was found in *Arabidopsis thaliana* (Bate et al., 1998), *Camellia sinensis* (tea) (Shi et al., 2015) and *Taxus chinensis* (Chinese yew) (Li et al., 2012).

In contrast to meJA, exogenous ACC or BTH did not trigger KTI expression, although marker gene analyses suggested intracellular responses to these compounds. *ETR1*, a marker for ethylene (Dong et al., 2008), was upregulated after ACC (precursor for ethylene) application. The responses to BTH (as SA analogue) were complex. A central function of SA is the activation of defenses against biotrophic pathogens via the receptor NPR1 (Withers and Dong, 2016). We observed a 2-fold decrease in *NPR1* upon BTH treatment. Similarly, Ullah et al. (Ullah et al., 2019) found only moderate variations in *NPR1* upon exposure to rust (*Melampsora larici-populina),* despite massive SA accumulation, presumably because inactive forms of NPR1 present in the cytosol are post-translationally activated by SA (Lorenzo and Solano, 2005; Zhou et al., 2023). Whether the increase in *ERF1* observed in our study under high BTH reflects the well-known antagonistic regulation of the SA and ethylene signaling pathways (Spoel et al., 2007) is speculative, especially, since these pathways are independently regulated in poplar (Ullah et al., 2022). Importantly, the application of BTH and ACC did not cause up-regulation of the transcript levels of any candidate KTI gene, which supports that SA and ET pathways were not involved in KTI induction. Hence, our results underpin a distinctive role of JAs for defense activation, likely downstream of JA production since the stimulation of *KTI*s was achieved in undamaged plants. In future studies, it will be interesting to dissect the molecular mechanisms resulting in divergent responses of distinct KTIs.

### 4.2 Functional analysis of KTIs and ecological aspects

In previous studies, the functions of poplar KTIs were investigated *in-vitro* or ectopically and revealed diversified patterns. For example, biochemical analyses of five recombinant KTIs from poplar showed different inhibitory profiles for commercial proteases as well as for proteases in midgut extracts from forest tent caterpillar (*Malacosoma disstria*) and armyworm (*Mamestra configurata*) (Major and Constabel, 2008). However, the greatest activity in the biochemical assays was exerted by a soybean (*Glycine max*) protease inhibitor (GmKTI) (Major and Constabel, 2008). Overexpression of *GmKTI* in poplar (*P. nigra*) resulted in inhibitory activity in transgenic leaf extracts; yet this transformation did not affect larval growth of polyphagous lepidopterans, *Lymantria dispar* and *Clostera anastomosis* feeding on the transgenic plants (Confalonieri et al., 1998). Ectopic overexpression of poplar KTIs (*PtdKTI2*, *PtdPOP3*, *PtdWIN4*, *PtKTI5*) in *Arabidopsis* resulted only in moderate reductions of larval growth but hampered proper larval development (Hu et al., 2012). In contrast to those preceding studies, we used *P.* x *canescens* as gene donor and host, employing overexpression under the constitutive *pS35* promoter and CRISPR/Cas12a for gene editing. The CRISPR/Cas12a system for plants was developed in *Arabidopsis thaliana*, where 21 % of editing efficiency was noted (Schindele and Puchta, 2020). We obtained similar editing efficiencies. The deletions ranged from 2 bp to 50 bp, and occurred at the 3’ distal side of the PAM as observed in rice and Arabidopsis (Bernabé-Orts *et al*., 2019; Malzahn *et al*., 2019; Schindele & Puchta, 2020).

Using this novel approach, we clearly showed that KTI_400 and KTI_600 are central in regulating the fitness of a generalist herbivore. Increased expression levels similar to those induced by leaf wounding or meJA exposure caused significant reductions in the weight gain of *H. armigera*, whereas *kti4+600* lines made leaves more palatable for the larvae. The greatest weight gain compared to controls occurred when the larvae fed on CRISPR/Cas12a mutant lines with the largest deletions. We did not attempt to generate single knock-out lines of *KTI_400* and *KTI_600* since their responses to meJA, wounding and *C. populi* feeding were similar, suggesting gene redundancy (Cusack et al., 2021). This presumption has yet to be tested. In *Arabidopsis thaliana*, suppression of single *KTI*s (*AtKTI4* and *AtKTI5* T-DNA insertion lines) effectively increased fecundity of spider mites (*Tetranychus urticae*) feeding on the transgenic plants (Arnaiz et al., 2018). However, the *Arabidopsis thaliana KTI* family is relatively small. Therefore, redundancy effects may be more likely in species with expanded *KTI* families such as *Populus*, especially for genes that share high similarity such as *KTI_400* and *KTI_600*.

In contrast to *KTI_400* and *KTI_600*, neither overexpression of *KTI_53200* nor its suppression in CRISPR/Cas12a lines affected the growth of the leaf-feeding *H. armigera* larvae. Previous transcriptomic studies of poplar tissues (e.g., *P. trichocarp*a (Shi et al., 2017)) found the highest expression levels of *KTI_53200* in roots (Guo et al., 2025). Since the KTI_53200 protein is abundant in the xylem sap of *P.* x *canescens* (Kasper et al. 2022), it is possible that KTI_53200 is produced in roots and transported upward the stem together with a wealth of other proteins (Dafoe and Constabel, 2009; Plomion et al., 2006). A possible function of KTI_53200 could be the control of serine proteases, which are also present in poplar sap (Kasper et al., 2022). However, this idea is speculative and needs further studies. Other possibilities are functions in plant development (Boex-Fontvieille et al., 2015; Havé et al., 2017), abiotic stresses (Islam et al., 2015), pathogen defense (Chen et al., 2021) or herbivore specificity (Eberl et al., 2021).

## Conclusions

Here, we tested only the generalist herbivore, *H. armigera*. *H. armigera* is a devastating pest in the (sub-)tropics, especially in Africa and Asia (Liu et al., 2004) but is also spreading across Europe and America (https://gd.eppo.int/taxon/HELIAR/datasheet). Its invasion into northern latitudes is limited by low winter temperatures, which prevent its survival. However, with increasing global temperatures, it is expected that *H. armigera* will become a threat to plants at higher latitudes (Subedi et al., 2023). Our study demonstrates the importance of KTI_400 and KTI_600 as defense against this pest. In future studies, it will also be necessary to test the fitness of specialist pests of poplars (Brückmann et al., 2002; Rank et al., 1998) such as *Chrysomela populi* and *Phratora vitellinae* on plants with modified KTI levels.

An unexpected result of our study was that the *kti4+600* lines showed greater biomass production than overexpressing or control poplars. This suggests that KTI production incurs fitness costs. In a previous study, fitness costs were only observed when *Nicotiana attenuata* plants with different levels of KTI were grown in neighborhood to each other, typically resulting in higher seed production in plants with lower KTI levels (Zavala et al., 2004). Zavala *et al*. (2004) suggested that KTI production is intrinsically costly when plants compete for belowground resources. However, other explanations than the classical idea of optimization of resource use efficiency, i.e., for defense or growth, are also possible. Recently, ecological theories on growth-defense allocational tradeoffs were interpreted in the light of molecular growth regulation, considering coordination by phytohormones and signaling cascades (Monson et al., 2022). Thereby, growth and defense may be partly uncoupled, opening new ways for a mechanistic understanding (Monson et al., 2022). The poplars generated in this study will be an ideal tool to dig deeper into the molecular mechanisms linking intrinsic KTI production with plant and insect fitness traits. Further investigation into the specific mechanisms and efficacy of KTIs in poplar defense systems could pave the way for developing novel biocontrol strategies against herbivorous pests.

## Conclusions

*KTIs* are differentially regulated in response to feeding of the poplar specialist *C. populii*. Overexpression of the *KTI*s with the strongest transcriptional response had negative effects on the fitness on larvae of the generalist *H. armigera*, while knock-out mutants showed increased larval growth. These results indicate a broad protective potential of KTI_400 and KTI_600 against herbivore insects and open perspectives for novel biocontrol measures.

## Declarations

### Ethics approval and consent to participate

Not applicable

### Consent for publication

Not applicable

### Availability of data and materials

Data generated or analysed during this study are included in this published article and its Additional information files. Materials are available upon request.

### Authors’ contributions

ISD: conceived study, conducted transformation and laboratory experiments, phytohormone treatment, data analysis, first draft, QQS: conducted greenhouse experiments, SD: conducted phytohormone treatments and laboratory experiments, AP: conceived study, secured funding, contributed to data analysis, supervision, revised drafts, all: commented and agreed on the final version.

## Acknowledgements

We are grateful to Prof. Dr. M. Rostás (Agricultural Entomology, Department for Crop Sciences, University of Göttingen, Göttingen) for provision of larvae for the herbivore experiments and to Prof. Dr. H. Puchta (Karlsruher Institut für Technologie (KIT), Botanisches Institut, Karlsruhe) for the provision of vectors for the CRISPR-Cas12a experiments. We thank M. Fastenrath and C. Leibecke (Forest Botany and Tree Physiology, University of Göttingen) for help with plant cultivation.

## Additional materials

### Additional Figures

Additional Figure S1: Phylogenetic tree of putative Kunitz Trypsin Inhibitors in *Populus trichocarpa* and the haplotype sequences of *P. tremula* and *P. alba*.

Additional Figure S2: Multiple sequence alignment of amino acids of the three candidate Kunitz Trypsin Inhibitor proteins

Additional Figure S3: a) *KTI* transcript levels of *P.* x *canescens* plantlets grown and wounded under sterile conditions, b) *KTI* (Potri019G08220, KTI_8220) transcript levels of greenhouse grown poplars after exposure to methyl-jasmonate (meJA), ACC or BTH.

Additional Figure S4: Transcript abundances of *KTI_400, KTI_600* and *KTI_53200* over-expressed under the *35S* promoter.

Additional Figure S5: Twelve-week-old Kunitz Trypsin Inhibitor mutant lines grown under greenhouse conditions.

Additional Figure S6: Representative photographs of Kunitz Trypsin Inhibitor poplar mutant lines under constant feeding of *Helicoverpa armigera* larvae.

### Additional Tables

Additional Table S1: List of all primer sets used for cloning and standard PCR

Additional Table S2: List of primers for RT-qPCR

Additional Table S3: Gene identity numbers for *P. trichocarpa* (Potri) and *P. tremula* (Potra)

Additional Table S4: Subcellular localization of the candidate KTIs

Additional Table S5: Description of transformed and surviving mutant lines of *Kunitz Trypsin Inhibitor* in *Populus* x *canescens*

Additional Table S6: Consequences of CRISPR-Cas12a editing events observed in mutant lines.

Additional Table S7: Gas exchange and growth of wildtype and transgenic poplar lines

## Additional Methods

## References

Abdul-Hussain S, Paulsen GM. Role of proteinaceous .alpha.-amylase enzyme inhibitors in preharvest sprouting of wheat grain. J Agric Food Chem 1989;37:295–9. 10.1021/jf00086a004.

Agback P, Agback T. Direct evidence of a low barrier hydrogen bond in the catalytic triad of a Serine protease. Sci Rep 2018;8:10078. 10.1038/s41598-018-28441-7.

Alborn HT, Turlings TCJ, Jones TH, Stenhagen G, Loughrin JH, Tumlinson JH. An elicitor of plant volatiles from beet armyworm oral secretion. Science 1997;276:945–9. 10.1126/science.276.5314.945.

Ali J, Mukarram M, Ojo J, Dawam N, Riyazuddin R, Ghramh HA, et al. Harnessing Phytohormones: Advancing Plant Growth and Defence Strategies for Sustainable Agriculture. Physiologia Plantarum 2024;176:e14307. 10.1111/ppl.14307.

do Amaral M, Freitas ACO, Santos AS, dos Santos EC, Ferreira MM, da Silva Gesteira A, et al. TcTI, a Kunitz-type trypsin inhibitor from cocoa associated with defense against pathogens. Sci Rep 2022;12:698. 10.1038/s41598-021-04700-y.

Amirkhosravi A, Strijkstra G-J, Keyl A, Häffner F, Lipka U, Herrfurth C, et al. Overexpression of jojoba wax ester synthase in poplar increases foliar lipid accumulation, alters stomatal conductance, and increases water use efficiency. Plant Biology 2025:doi: 10.1111/plb.70056. 10.1111/plb.70056.

Arnaiz A, Talavera-Mateo L, Gonzalez-Melendi P, Martinez M, Diaz I, Santamaria ME. Arabidopsis Kunitz trypsin inhibitors in defense against spider mites. Frontiers in Plant Science 2018;9. 10.3389/fpls.2018.00986.

Babst BA, Sjödin A, Jansson S, Orians CM. Local and systemic transcriptome responses to herbivory and jasmonic acid in Populus. Tree Genetics & Genomes 2009;5:459–74. 10.1007/s11295-009-0200-6.

Bate NJ, Sivasankar S, Moxon C, Riley JMC, Thompson JE, Rothstein SJ. Molecular Characterization of an Arabidopsis Gene Encoding Hydroperoxide Lyase, a Cytochrome P-450 That Is Wound Inducible1. Plant Physiology 1998;117:1393–400. 10.1104/pp.117.4.1393.

Bates D, Mächler M, Bolker B, Walker S. Fitting Linear Mixed-Effects Models Using lme4. Journal of Statistical Software 2015;67:1–48. 10.18637/jss.v067.i01.

Bendre AD, Ramasamy S, Suresh CG. Analysis of Kunitz inhibitors from plants for comprehensive structural and functional insights. International Journal of Biological Macromolecules 2018;113:933–43. 10.1016/j.ijbiomac.2018.02.148.

Bernabé-Orts JM, Casas-Rodrigo I, Minguet EG, Landolfi V, Garcia-Carpintero V, Gianoglio S, et al. Assessment of Cas12a-mediated gene editing efficiency in plants. Plant Biotechnology Journal 2019;17:1971–84. 10.1111/pbi.13113.

Boex-Fontvieille E, Rustgi S, Reinbothe S, Reinbothe C. A Kunitz-type protease inhibitor regulates programmed cell death during flower development in Arabidopsis thaliana. Journal of Experimental Botany 2015;66:6119–35. 10.1093/jxb/erv327.

Bonturi CR, Silva Teixeira AB, Rocha VM, Valente PF, Oliveira JR, Filho CMB, et al. Plant Kunitz inhibitors and their interaction with proteases: current and potential pharmacological targets. International Journal of Molecular Sciences 2022;23:4742. 10.3390/ijms23094742.

Bradshaw HD, Hollick JB, Parsons TJ, Clarke HRG, Gordon MP. Systemically wound-responsive genes in poplar trees encode proteins similar to sweet potato sporamins and legume Kunitz trypsin inhibitors. Plant Mol Biol 1990;14:51–9. 10.1007/BF00015654.

Brückmann M, Termonia A, Pasteels JM, Hartmann T. Characterization of an extracellular salicyl alcohol oxidase from larval defensive secretions of *Chrysomela populi* and *Phratora vitellinae* (Chrysomelina). Insect Biochemistry and Molecular Biology 2002;32:1517–23. 10.1016/S0965-1748(02)00072-3.

Bruegmann T, Polak O, Deecke K, Nietsch J, Fladung M. Poplar Transformation. Methods Mol Biol 2019;1864:165–77. 10.1007/978-1-4939-8778-8_12.

Chen Q, Zhang R, Li D, Wang F. Integrating transcriptome and coexpression network analyses to characterize salicylic acid- and jasmonic acid-related genes in tolerant poplars infected with rust. International Journal of Molecular Sciences 2021;22:5001. 10.3390/ijms22095001.

Christopher ME, Miranda M, Major IT, Constabel CP. Gene expression profiling of systemically wound-induced defenses in hybrid poplar. Planta 2004;219:936–47. 10.1007/s00425-004-1297-3.

Clemente M, Corigliano MG, Pariani SA, Sánchez-López EF, Sander VA, Ramos-Duarte VA. Plant serine protease inhibitors: biotechnology application in agriculture and molecular farming. Int J Mol Sci 2019;20:1345. 10.3390/ijms20061345.

Conconi A, Miquel M, Browse JA, Ryan CA. Intracellular Levels of Free Linolenic and Linoleic Acids Increase in Tomato Leaves in Response to Wounding. Plant Physiol 1996;111:797–803.

Confalonieri M, Allegro G, Balestrazzi A, Fogher C, Delledonne M. Regeneration of Populus nigra transgenic plants expressing a Kunitz proteinase inhibitor (KTi3) gene. Molecular Breeding 1998;4:137–45. 10.1023/A:1009640204314.

Constabel CP, L. Lindroth R. The Impact of Genomics on Advances in Herbivore Defense and Secondary Metabolism in Populus. In: Jansson S, Bhalerao R, Groover A, editors. Genetics and Genomics of Populus, New York, NY: Springer; 2010, p. 279–305. 10.1007/978-1-4419-1541-2_13.

Cusack SA, Wang P, Lotreck SG, Moore BM, Meng F, Conner JK, et al. Predictive models of genetic redundancy in *Arabidopsis thaliana*. Molecular Biology and Evolution 2021;38:3397–414. 10.1093/molbev/msab111.

Dafoe NJ, Constabel CP. Proteomic analysis of hybrid poplar xylem sap. Phytochemistry 2009;70:856–63. 10.1016/j.phytochem.2009.04.016.

Divekar PA, Narayana S, Divekar BA, Kumar R, Gadratagi BG, Ray A, et al. Plant Secondary Metabolites as Defense Tools against Herbivores for Sustainable Crop Protection. International Journal of Molecular Sciences 2022;23:2690. 10.3390/ijms23052690.

Dong C-H, Rivarola M, Resnick JS, Maggin BD, Chang C. Subcellular co-localization of Arabidopsis RTE1 and ETR1 supports a regulatory role for RTE1 in ETR1 ethylene signaling. The Plant Journal 2008;53:275–86. 10.1111/j.1365-313X.2007.03339.x.

Eberl F, Fabisch T, Luck K, Köllner TG, Vogel H, Gershenzon J, et al. Poplar protease inhibitor expression differs in an herbivore specific manner. BMC Plant Biology 2021;21:170. 10.1186/s12870-021-02936-4.

Erb M, Meldau S, Howe GA. Role of phytohormones in insect-specific plant reactions. Trends Plant Sci 2012;17:250–9. 10.1016/j.tplants.2012.01.003.

Fear G, Komarnytsky S, Raskin I. Protease inhibitors and their peptidomimetic derivatives as potential drugs. Pharmacol Ther 2007;113:354–68. 10.1016/j.pharmthera.2006.09.001.

Feussner I, Wasternack C. The lipoxygenase pathway. Annu Rev Plant Biol 2002;53:275–97. 10.1146/annurev.arplant.53.100301.135248.

Fox J, Weisberg S. An R Companion to Applied Regression,. Third Edition. Thousand Oaks CA: Sage; 2019.

Gergs A, Baden CU. A dynamic energy budget approach for the prediction of development times and variability in *Spodoptera frugiperda* rearing. Insects 2021;12:300. 10.3390/insects12040300.

Grosse-Holz FM, van der Hoorn RAL. Juggling jobs: roles and mechanisms of multifunctional protease inhibitors in plants. New Phytologist 2016;210:794–807. 10.1111/nph.13839.

Guo H, Ma S, Zhang X, Xu R, Wang C, Zhang S, et al. Identification of Kunitz-Type Inhibitor Gene Family of Populus yunnanensis Reveals a Stress Tolerance Function in Inverted Cuttings. International Journal of Molecular Sciences 2025;26:188. 10.3390/ijms26010188.

Haruta M, Major IT, Christopher ME, Patton JJ, Constabel CP. A Kunitz trypsin inhibitor gene family from trembling aspen (*Populus tremuloides* Michx.): cloning, functional expression, and induction by wounding and herbivory. Plant Mol Biol 2001;46:347–59. 10.1023/a:1010654711619.

Havé M, Marmagne A, Chardon F, Masclaux-Daubresse C. Nitrogen remobilization during leaf senescence: lessons from Arabidopsis to crops. Journal of Experimental Botany 2017;68:2513–29. 10.1093/jxb/erw365.

Hothorn T, Bretz F, Westfall P. Simultaneous Inference in General Parametric Models. Biometrical Journal 2008;50:346–63. 10.1002/bimj.200810425.

Hu R, Wang J, Ji Y, Song Y, Yang S. Overexpression of poplar wounding-inducible genes in Arabidopsis caused improved resistance against Helicoverpa armigera (Hübner) larvae. Breed Sci 2012;62:288–91. 10.1270/jsbbs.62.288.

Islam A, Leung S, Burgess EPJ, Laing WA, Richardson KA, Hofmann RW, et al. Knock-down of transcript abundance of a family of Kunitz proteinase inhibitor genes in white clover (*Trifolium repens*) reveals a redundancy and diversity of gene function. New Phytologist 2015;208:1188–201. 10.1111/nph.13543.

Joshi RS, Mishra M, Suresh CG, Gupta VS, Giri AP. Complementation of intramolecular interactions for structural– functional stability of plant serine proteinase inhibitors. Biochimica et Biophysica Acta (BBA) - General Subjects 2013;1830:5087–94. 10.1016/j.bbagen.2013.07.019.

Kaling M, Schmidt A, Moritz F, Rosenkranz M, Witting M, Kasper K, et al. Mycorrhiza-triggered transcriptomic and metabolomic networks impinge on herbivore fitness. Plant Physiology 2018;176:2639–56. 10.1104/pp.17.01810.

Karimi M, Inzé D, Depicker A. Gateway vectors for Agrobacterium-mediated plant transformation. Trends in Plant Science 2002;7:193–5. 10.1016/S1360-1385(02)02251-3.

Kasper K, Abreu IN, Feussner K, Zienkiewicz K, Herrfurth C, Ischebeck T, et al. Multi-omics analysis of xylem sap uncovers dynamic modulation of poplar defenses by ammonium and nitrate. PLANT JOURNAL 2022;111:282–303. 10.1111/tpj.15802.

Katz BA, Elrod K, Luong C, Rice MJ, Mackman RL, Sprengeler PA, et al. A novel serine protease inhibition motif involving a multi-centered short hydrogen bonding network at the active site1. Journal of Molecular Biology 2001;307:1451–86. 10.1006/jmbi.2001.4516.

Li S, Zhang P, Zhang M, Fu C, Zhao C, Dong Y, et al. Transcriptional profile of Taxus chinensis cells in response to methyl jasmonate. BMC Genomics 2012;13:295. 10.1186/1471-2164-13-295.

Liu Z, Li D, Gong P, Wu K. Life Table Studies of the Cotton Bollworm, Helicoverpa armigera (Hübner) (Lepidoptera: Noctuidae), on Different Host Plants. Environ Entomol 2004;33:1570–6. 10.1603/0046-225X-33.6.1570.

Lorenzo O, Solano R. Molecular players regulating the jasmonate signalling network. Curr Opin Plant Biol 2005;8:532–40. 10.1016/j.pbi.2005.07.003.

Lortzing T, Steppuhn A. Jasmonate signalling in plants shapes plant–insect interaction ecology. Current Opinion in Insect Science 2016;14:32–9. 10.1016/j.cois.2016.01.002.

Ma Y, Zhao Q, Lu M-Z, Wang J. Kunitz-type trypsin inhibitor gene family in Arabidopsis and Populus trichocarpa and its expression response to wounding and herbivore in Populus nigra. Tree Genetics & Genomes 2011;7:431–41. 10.1007/s11295-010-0345-3.

Major IT, Constabel CP. Functional Analysis of the Kunitz Trypsin Inhibitor Family in Poplar Reveals Biochemical Diversity and Multiplicity in Defense against Herbivores. Plant Physiology 2008;146:888–903. 10.1104/pp.107.106229.

Major IT, Constabel CP. Insect regurgitant and wounding elicit similar defense responses in poplar leaves: Not something to spit at? Plant Signaling & Behavior 2007;2:1–3. 10.4161/psb.2.1.3589.

Major IT, Constabel CP. Molecular analysis of poplar defense against herbivory: Comparison of wound- and insect elicitor-induced gene expression. The New Phytologist 2006;172:617–35. 10.1111/j.1469-8137.2006.01877.x.

Malzahn AA, Tang X, Lee K, Ren Q, Sretenovic S, Zhang Yingxiao, et al. Application of CRISPR-Cas12a temperature sensitivity for improved genome editing in rice, maize, and Arabidopsis. BMC Biology 2019;17:9. 10.1186/s12915-019-0629-5.

McCormick AC, Reinecke A, Gershenzon J, Unsicker SB. Feeding experience affects the behavioral response of polyphagous gypsy moth caterpillars to herbivore-induced poplar volatiles. J Chem Ecol 2016;42:382–93. 10.1007/s10886-016-0698-7.

Mehmood S, Thirup SS, Ahmed S, Bashir N, Saeed A, Rafiq M, et al. Crystal structure of Kunitz-type trypsin inhibitor: Entomotoxic effect of native and encapsulated protein targeting gut trypsin of *Tribolium castaneum* Herbst. Computational and Structural Biotechnology Journal 2024;23:3132–42. 10.1016/j.csbj.2024.07.023.

Merker L, Schindele P, Huang TK, Wolter F, Puchta H. Enhancing in planta gene targeting efficiencies in Arabidopsis using temperature-tolerant CRISPR/LbCas12a. Plant Biotechnology Journal 2020. 10.1111/pbi.13426.

Metsalu T, Vilo J. ClustVis: a web tool for visualizing clustering of multivariate data using Principal Component Analysis and heatmap. Nucleic Acids Res 2015;43:W566–570. 10.1093/nar/gkv468.

Mironidis GK, Savopoulou-Soultani M. Development, survivorship, and reproduction of *Helicoverpa armigera* (Lepidoptera: noctuidae) under constant and alternating temperatures. ENVIRONMENTAL ENTOMOLOGY 2008;37.

Monson RK, Trowbridge AM, Lindroth RL, Lerdau MT. Coordinated resource allocation to plant growth–defense tradeoffs. New Phytologist 2022;233:1051–66. 10.1111/nph.17773.

Müller A, Volmer K, Mishra-Knyrim M, Polle A. Growing poplars for research with and without mycorrhizas. Frontiers in Plant Science 2013;4:332. 10.3389/fpls.2013.00332.

Murashige T, Skoog F. A revised medium for rapid growth and bio assays with tobacco tissue cultures. Physiologia Plantarum 1962;15:473–97. 10.1111/j.1399-3054.1962.tb08052.x.

Oliva MLV, Silva MCC, Sallai RC, Brito MV, Sampaio MU. A novel subclassification for Kunitz proteinase inhibitors from leguminous seeds. Biochimie 2010;92:1667–73. 10.1016/j.biochi.2010.03.021.

Oliveira AS, Migliolo L, Aquino RO, Ribeiro JKC, Macedo LLP, Bemquerer MP, et al. Two Kunitz-type inhibitors with activity against trypsin and papain from Pithecellobium dumosum seeds: purification, characterization, and activity towards pest insect digestive enzyme. Protein Pept Lett 2009;16:1526–32. 10.2174/092986609789839403.

Patel S. A critical review on serine protease: Key immune manipulator and pathology mediator. Allergol Immunopathol (Madr) 2017;45:579–91. 10.1016/j.aller.2016.10.011.

Pekkarinen AI, Longstaff C, Jones BL. Kinetics of the Inhibition of Fusarium Serine Proteinases by Barley (Hordeum vulgare L.) Inhibitors. J Agric Food Chem 2007;55:2736–42. 10.1021/jf0631777.

Pfaffl MW. A new mathematical model for relative quantification in real-time RT-PCR. Nucleic Acids Research 2001;29:0. 10.1093/nar/29.9.e45.

Philippe RN, Bohlmann J. Poplar defense against insect herbivores. Canadian Journal of Botany 2007;85:1111–26. 10.1139/B07-109.

Philippe RN, Ralph SG, Külheim C, Jancsik SI, Bohlmann J. Poplar defense against insects: Genome analysis, full-length cDNA cloning, and transcriptome and protein analysis of the poplar Kunitz-type protease inhibitor family. New Phytologist 2009;184:865–84. 10.1111/j.1469-8137.2009.03028.x.

Plomion C, Lalanne C, Claverol S, Meddour H, Kohler A, Bogeat-Triboulot M-B, et al. Mapping the proteome of poplar and application to the discovery of drought-stress responsive proteins. Proteomics 2006;6:6509–27. 10.1002/pmic.200600362.

Pureswaran DS, Roques A, Battisti A. Forest insects and climate change. Curr Forestry Rep 2018;4:35–50. 10.1007/s40725-018-0075-6.

Qi J, Zhou G, Yang L, Erb M, Lu Y, Sun X, et al. The Chloroplast-Localized Phospholipases D α4 and α5 Regulate Herbivore-Induced Direct and Indirect Defenses in Rice1[C][W]. Plant Physiol 2011;157:1987–99. 10.1104/pp.111.183749.

Rank NE, Köpf A, Julkunen-Tiitto R, Tahvanainen J. Host Preference and Larval Performance of the Salicylate- Using Leaf Beetle Phratora Vi℡linae. Ecology 1998;79:618–31. 10.1890/0012-9658(1998)079%255B0618:HPALPO%255D2.0.CO;2.

Rogers PC, Pinno BD, Šebesta J, Albrectsen BR, Li G, Ivanova N, et al. A global view of aspen: Conservation science for widespread keystone systems. Global Ecology and Conservation 2020;21:e00828. 10.1016/j.gecco.2019.e00828.

Rustgi S, Boex-Fontvieille E, Reinbothe C, von Wettstein D, Reinbothe S. The complex world of plant protease inhibitors: Insights into a Kunitz-type cysteine protease inhibitor of *Arabidopsis thaliana*. Commun Integr Biol 2017;11:e1368599. 10.1080/19420889.2017.1368599.

Schäfer M, Fischer C, Meldau S, Seebald E, Oelmüller R, Baldwin IT. Lipase Activity in Insect Oral Secretions Mediates Defense Responses in Arabidopsis. Plant Physiology 2011;156:1520–34. 10.1104/pp.111.173567.

Schindele P, Puchta H. Engineering CRISPR/LbCas12a for highly efficient, temperature-tolerant plant gene editing. Plant Biotechnology Journal 2020;18:1118–20. 10.1111/pbi.13275.

Schmelz EA, Carroll MJ, LeClere S, Phipps SM, Meredith J, Chourey PS, et al. Fragments of ATP synthase mediate plant perception of insect attack. Proceedings of the National Academy of Sciences 2006;103:8894–9. 10.1073/pnas.0602328103.

Shi J, Ma C, Qi D, Lv H, Yang T, Peng Q, et al. Transcriptional responses and flavor volatiles biosynthesis in methyl jasmonate-treated tea leaves. BMC Plant Biology 2015;15:233. 10.1186/s12870-015-0609-z.

Shi R, Wang JP, Lin Y-C, Li Q, Sun Y-H, Chen H, et al. Tissue and cell-type co-expression networks of transcription factors and wood component genes in Populus trichocarpa. Planta 2017;245:927–38. 10.1007/s00425-016-2640-1.

Sivaprakasam Padmanaban PB, Rosenkranz M, Zhu P, Kaling M, Schmidt A, Schmitt-Kopplin P, et al. Mycorrhiza- Tree-Herbivore Interactions: Alterations in Poplar Metabolome and Volatilome. METABOLITES 2022;12. 10.3390/metabo12020093.

Spoel SH, Johnson JS, Dong X. Regulation of tradeoffs between plant defenses against pathogens with different lifestyles. Proceedings of the National Academy of Sciences 2007;104:18842–7. 10.1073/pnas.0708139104.

Stanton BJ, Bourque A, Coleman M, Eisenbies M, Emerson RM, Espinoza J, et al. The practice and economics of hybrid poplar biomass production for biofuels and bioproducts in the Pacific Northwest. Bioenerg Res 2021;14:543–60. 10.1007/s12155-020-10164-1.

Subedi B, Poudel A, Aryal S. The impact of climate change on insect pest biology and ecology: Implications for pest management strategies, crop production, and food security. Journal of Agriculture and Food Research 2023;14:100733. 10.1016/j.jafr.2023.100733.

Ullah C, Schmidt A, Reichelt M, Tsai C-J, Gershenzon J. Lack of antagonism between salicylic acid and jasmonate signalling pathways in poplar. New Phytologist 2022;235:701–17. 10.1111/nph.18148.

Ullah C, Tsai C-J, Unsicker SB, Xue L, Reichelt M, Gershenzon J, et al. Salicylic acid activates poplar defense against the biotrophic rust fungus Melampsora larici-populina via increased biosynthesis of catechin and proanthocyanidins. New Phytologist 2019;221:960–75. 10.1111/nph.15396.

Vick BA, Zimmerman DC. The biosynthesis of jasmonic acid: A physiological role for plant lipoxygenase. Biochemical and Biophysical Research Communications 1983;111:470–7. 10.1016/0006-291X(83)90330-3.

Withers J, Dong X. Posttranslational Modifications of NPR1: A Single Protein Playing Multiple Roles in Plant Immunity and Physiology. PLOS Pathogens 2016;12:e1005707. 10.1371/journal.ppat.1005707.

Zavala JA, Patankar AG, Gase K, Baldwin IT. Constitutive and inducible trypsin proteinase inhibitor production incurs large fitness costs in *Nicotiana attenuata*. Proc Natl Acad Sci USA 2004;101:1607–12. 10.1073/pnas.0305096101.

Zhou P, Zavaliev R, Xiang Y, Dong X. Seeing is believing: Understanding functions of NPR1 and its paralogs in plant immunity through cellular and structural analyses. Current Opinion in Plant Biology 2023;73:102352. 10.1016/j.pbi.2023.102352.

Zhu J, He Y, Yan X, Liu L, Guo R, Xia X, et al. Duplication and transcriptional divergence of three Kunitz protease inhibitor genes that modulate insect and pathogen defenses in tea plant (Camellia sinensis). Hortic Res 2019;6:126. 10.1038/s41438-019-0208-5.

