## Additional_materials for "Divergent functions of three Kunitz trypsin inhibitor (KTI) proteins in herbivore defense in poplar": Additional_Figures_and_Tables.pdf

for

**Additional Figures**

Additional Figure S1: Phylogenetic tree of putative Kunitz Trypsin Inhibitors in *Populus trichocarpa* and the haplotype sequences of *P. tremula* and *P. alba*.
Additional Figure S2: Multiple sequence alignment of amino acids of the three candidate Kunitz Trypsin Inhibitor proteins
Additional Figure S3: a) *KTI* transcript levels of *P.* x *canescens* plantlets grown and wounded under sterile conditions, b) *KTI* (Potri019G08220, KTI_8220) transcript levels of greenhouse grown poplars after exposure to meJA, ACC or BTH.
Additional Figure S4: Transcript abundances of *KTI_400, KTI_600* and *KTI_53200* over-expressed under the p*35S* promoter.
Additional Figure S5: Twelve-week-old Kunitz Trypsin Inhibitor mutant lines grown under greenhouse conditions.
Additional Figure S6: Representative photographs of Kunitz Trypsin Inhibitor poplar mutant lines under constant feeding of *Helicoverpa armigera* larvae.

**Additional Tables**

Additional Table S1: List of all primer sets used for cloning and standard PCR
Additional Table S2: List of primers for RT-qPCR
Additional Table S3: Gene identity numbers for *P. trichocarpa* (Potri) and *P. tremula* (Potra)
Additional Table S4: Subcellular localization of the candidate KTIs
Additional Table S5: Description of transformed and surviving mutant lines of *Kunitz Trypsin Inhibitor* in *Populus* x *canescens*
Additional Table S6: Consequences of CRISPR-Cas12a editing events observed in mutant lines.
Additional Table S7: Gas exchange and growth of wildtype and transgenic poplar lines

**Additional Methods**

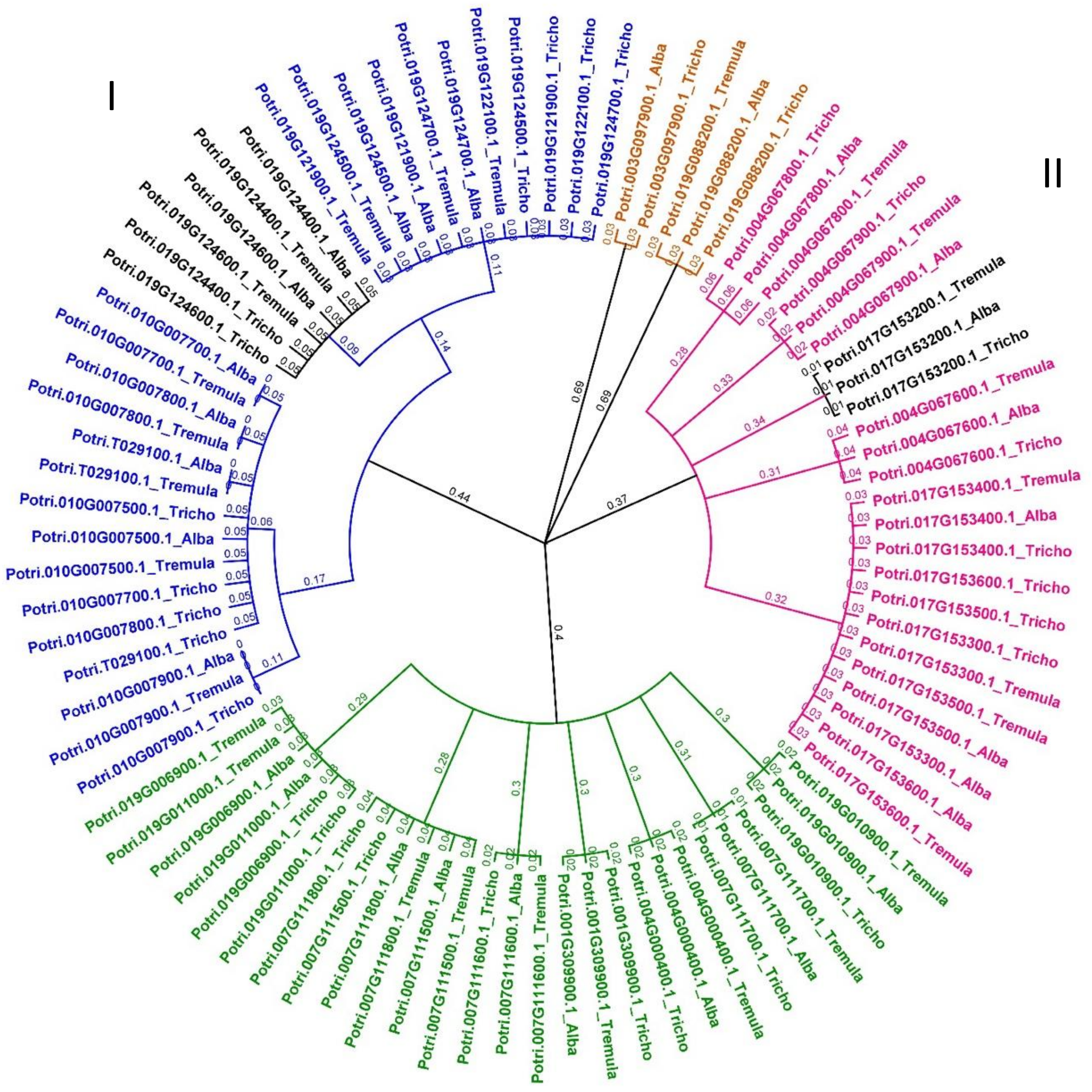


**Additional Figure S1:** Phylogenetic tree of putative Kunitz Trypsin Inhibitors in *Populus trichocarpa* and the haplotype sequences of *P. tremula* and *P. alba*. Unrooted UPGMA tree was generated with full-length amino acid sequences*.* sPta717. *P.* x *canescens* is a hybrid of *Populus tremula* and *Populus alba;* thus, the whole-genome sequence of *P.* x *canescens* comprises the haplotype sequences to its parents. Suffix "_Alba" or "_Tremula" denoted respective sequences from the crossing parents and "_Tricho" for the *P. trichocarpa* genome. Jukes-Cantor genetic distance model was applied with the consensus method: strict greedy clustering (Bootstrap n = 1000) in Geneious Prime (Biomatters, Ltd., Auckland, New Zealand). Colored branches denote specific clades represented as I-III. Nodes colored orange represent the KTI with the least similarity to the other KTI, and black nodes denote candidate genes in this study. Numbers denote substitutions per site used for the pairwise alignment.

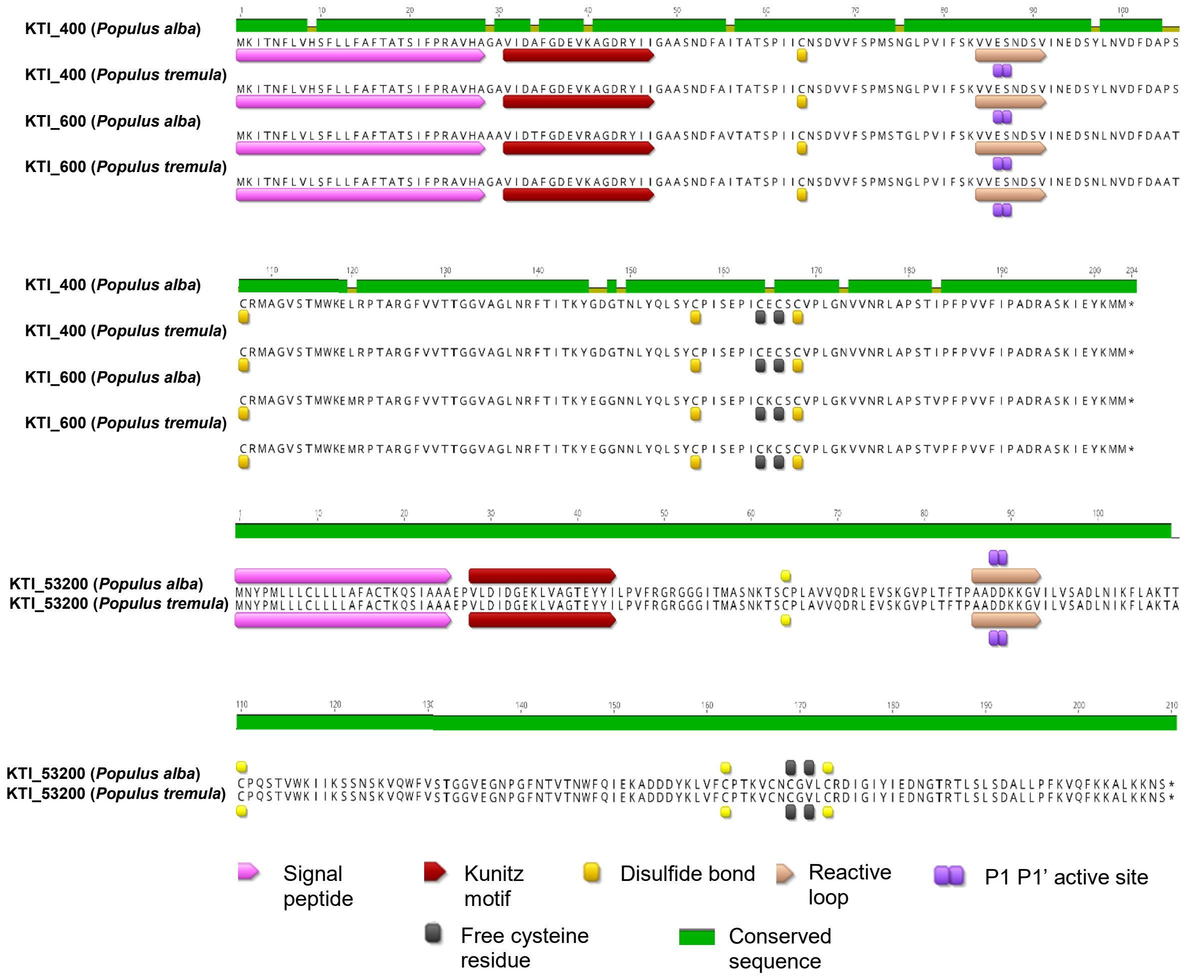


**Additional Figure S2:** Multiple sequence alignment of amino acids of the three candidate Kunitz Trypsin Inhibitor proteins. *Populus* x *canescens* is a hybrid of *Populus tremula* and *Populus alba;* thus, version 2 of sPta717 *P.* x *canescens* comprises the haplotype whole-genome sequences of both parents. Multiple sequence alignment performed in Clustal Omega 1.2.2 embedded in Geneious Prime (Biomatters, Ltd., Auckland, New Zealand). Kunitz motif and cysteine residues along with signal peptide, reactive loop, P1 P1' active sites, and free cysteine residues characteristic to KTIs were annotated.

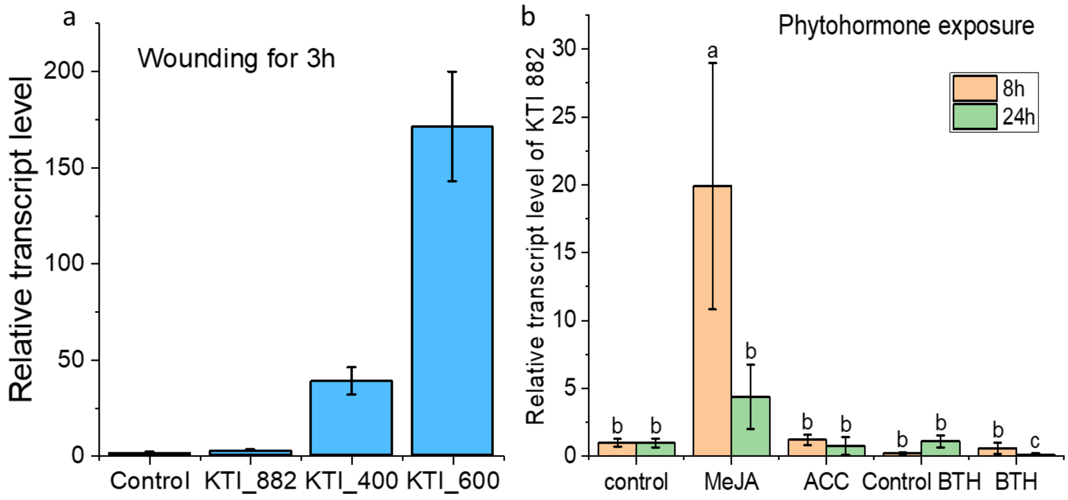


**Additional Figure S3**: a) KTI transcript levels of P x canescens plantlets grown and wounded under sterile conditions, b) KTI_Potri019G08220 (KTI_8220) transcript levels of greenhouse grown poplars after exposure to methyl-jasmonate (meJA), ACC or BTH. Leaves were harvested after 3h in the wounding experiments and after 8h and 24h in the phytohormone experiment post treatment. N = 4 ± SD biological replicates per treatment.

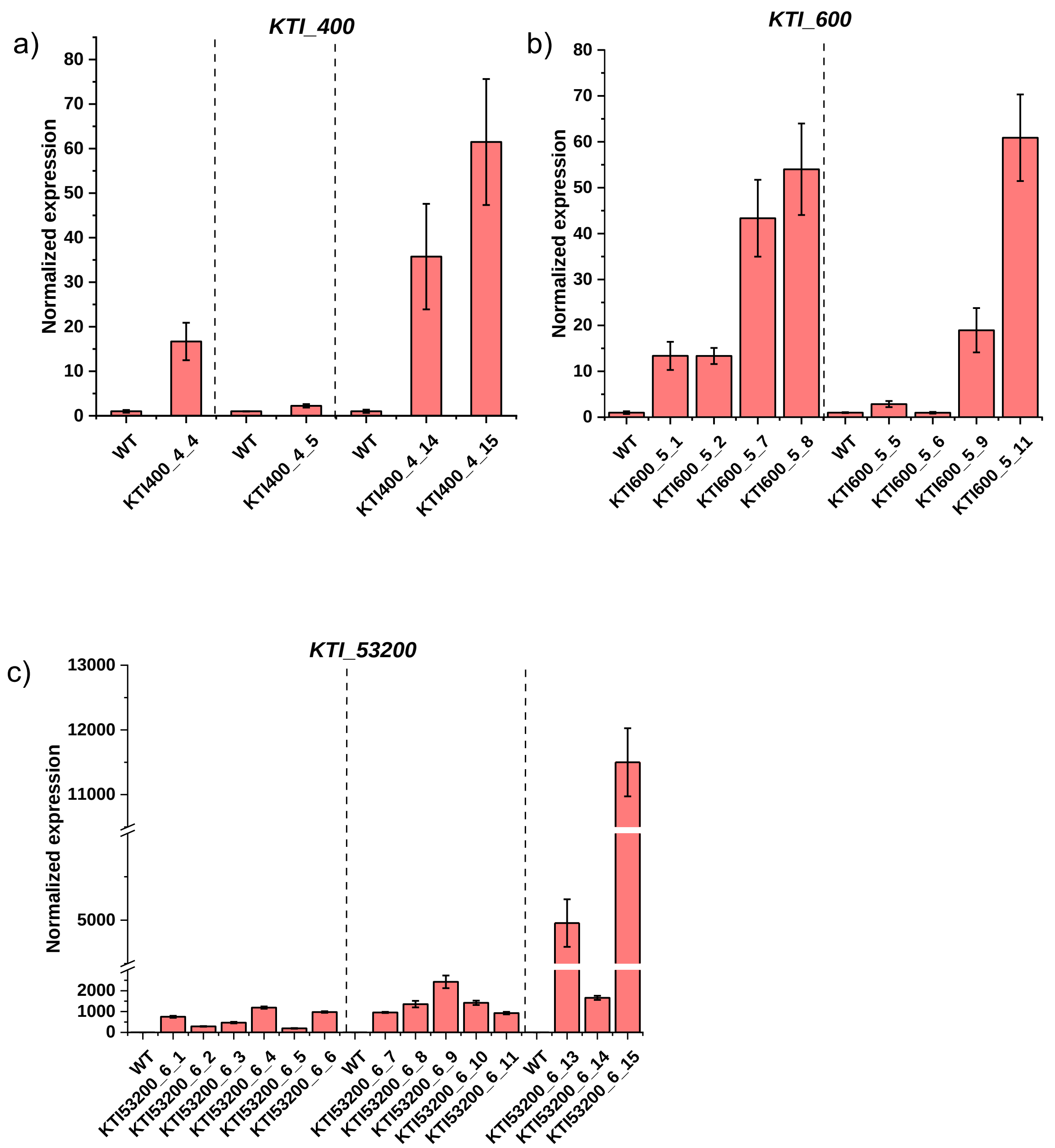


**Additional Figure S4:** Candidate *Kunitz Trypsin Inhibitor* genes over-expressed under the *35S* promoter. Transcript abundances of the independently transformed lines with the genes a) *KTI_400,* b) *KTI_600* and c) *KTI_53200* were tested. Transcript abundances were normalized to the reference genes, *ACTIN* and *UBIQUITIN* and were tested in comparison to the wild-type (WT) plants. Dotted lines refer to distinct experimental sets of tested plants and thus, the corresponding WT from the same experiment was compared to the mutant lines within these sets. Genotyping was performed on a single plant from each line. Error bars denote the standard deviation across three technical replicates used for RT-qPCR.

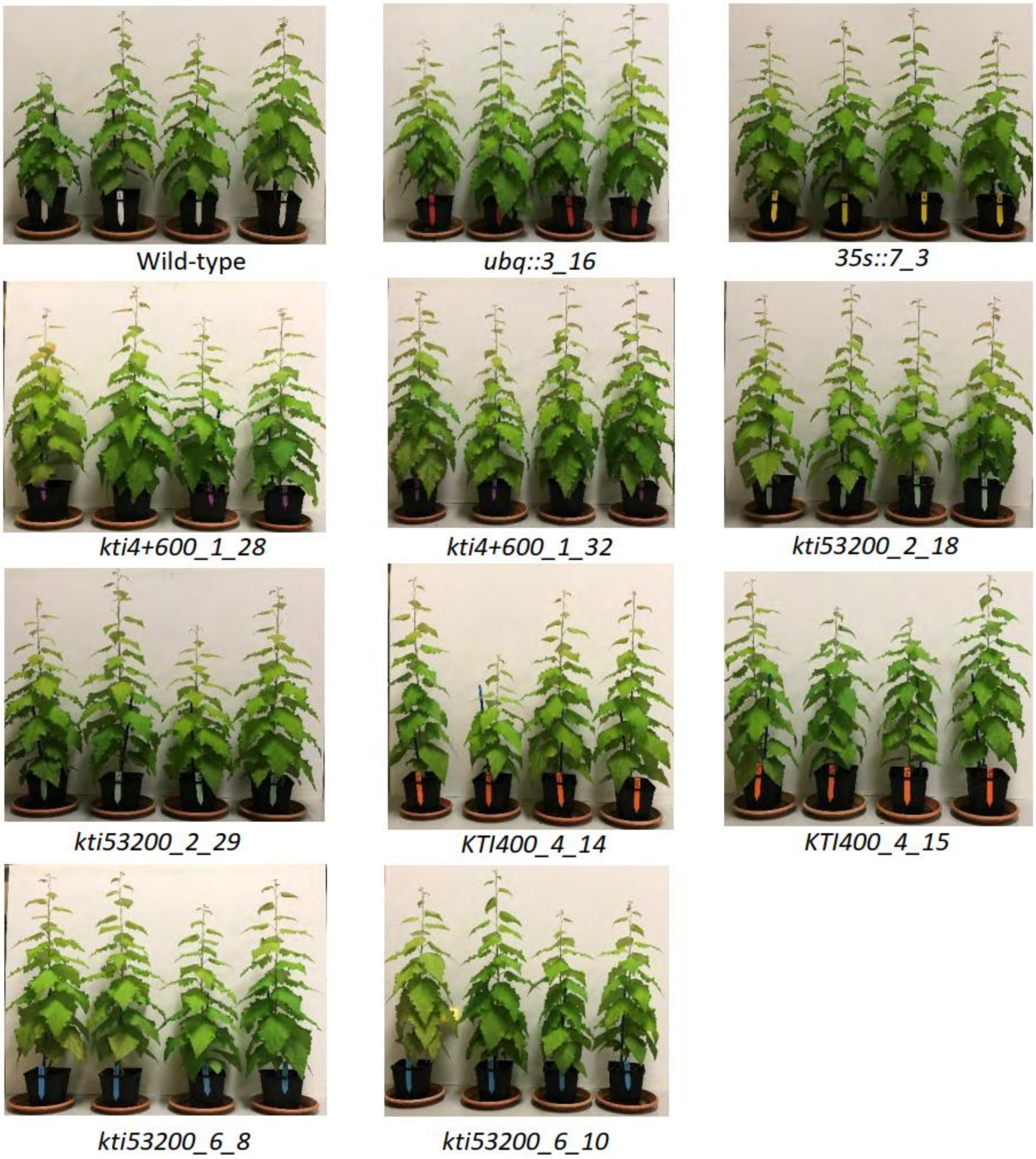


**Additional Figure S5:** Twelve-week-old Kunitz Trypsin Inhibitor mutant lines grown under greenhouse conditions. The panels show four biological replicates. WT, Wild type.

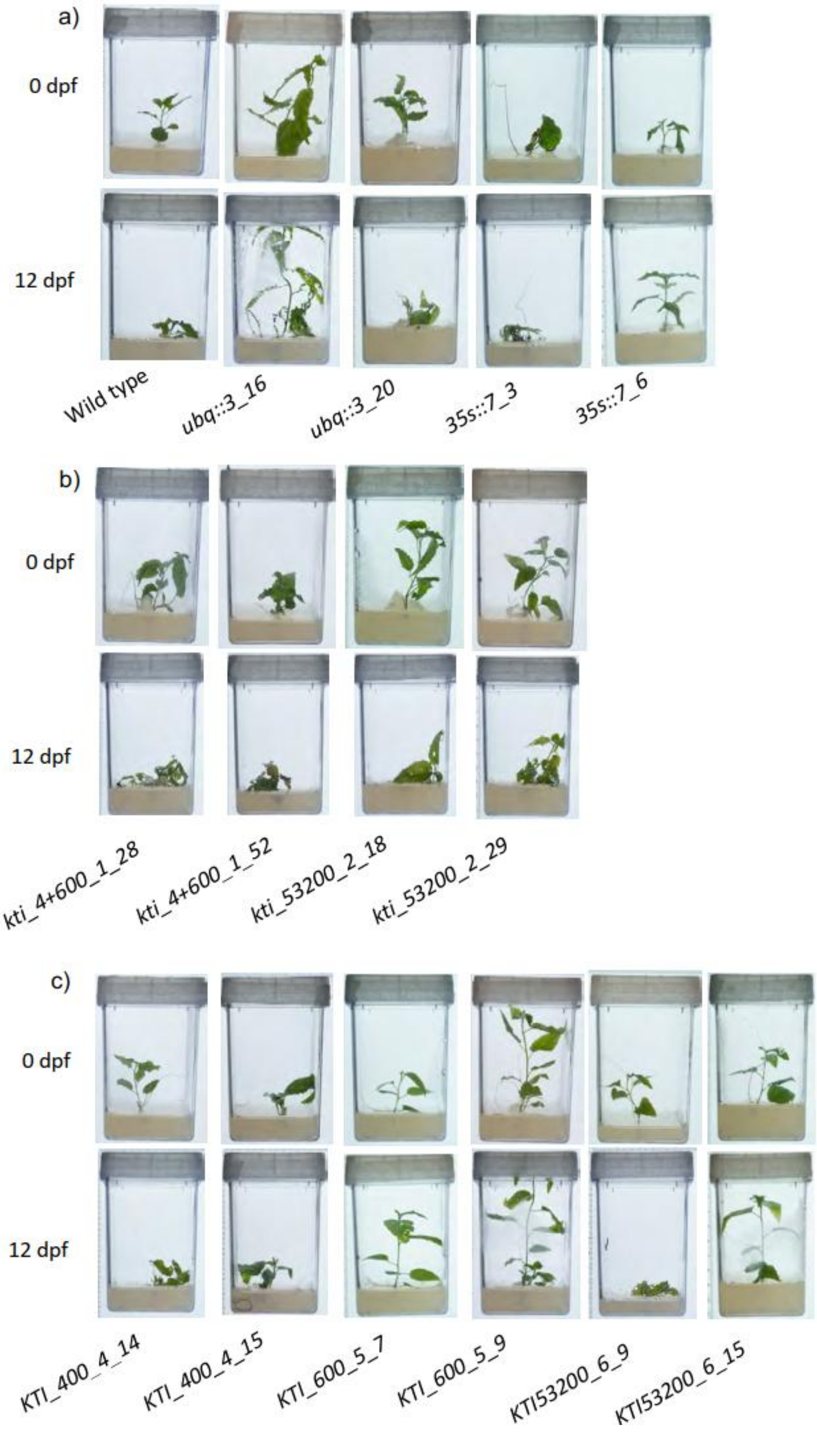


**Additional Figure S6**: Representative photographs of Kunitz Trypsin Inhibitor poplar mutant lines under constant feeding of *Helicoverpa armigera* larvae. Photos were taken at 0 dpf (left photograph in the panel) and 12 dpf (right photograph in the panel). dpf, days post-feeding; WT, Wildtype

**Additional Table S1:** List of all primer sets used for cloning and standard PCR. *Potri.019G124400* corresponds to *KTI_400, Potri.019G124600* to *KTI_600* and *Potri.017G153200* to *KTI_53200.*

| Primer Name Forward | Sequence (5′→3′) | Primer Name Reverse | Sequence (5′→3′) | Amplicon size (bp) | Annealing Temperature (°C) | Purpose |
|---|---|---|---|---|---|---|
| KTI_6_F_2 | ATATTTCCTCGTGCC GTTCATGCTG | KTI_6_R | GCATGAGCATTCACATAT GGGTTCG | 444 | 66 | *Potri.019G124400* gene and Target_400_600 specific |
| KTI_6_F_2 | ATATTTCCTCGTGCC GTTCATGCTG | KTI_4_R_11 | GGAATTATCATGTTTTAAT GCTGTGAGAG | 739 | 64 | *Potri.019G124600* (Nested PCR outer amplicon) |
| KTI_6_F_2 | ATATTTCCTCGTGCC GTTCATGCTG | KTI_4_R_5 | TTGATACGCCCGCCATTCT ACATGT | 280 | 66 | *Potri.019G124600* (Nested PCR inner amplicon) and Target_400_600_specific |
| KTI_5_F | GATGGTGAAAAGCTT GTAGCAGG | KTI_5_R | CCATACAGTTGATTGGGG ACATG | 255 | 55 | *Potri.017G153200* specific and Target_53200 specific |
| SS42 | TCCCAGGATTAGAAT GATTAGG | SS143 | CAAGAAAGCTGGGTCCT CAG | 441 | 55 | pChimera vector specific |
| SS42 | TCCCAGGATTAGAAT GATTAGG | 400_600_Target_R | GGCCTGATAGGGCTAGTC GCAGTGATCG | 348 | 56 | Target_400_600_integration to pChimera vector |
| SS42 | TCCCAGGATTAGAAT GATTAGG | 53200_Target_R | GGCCTTTCTTGTCGTCGG CAGCTGGTGT | 348 | 56 | Target_53200_integration to pChimera vector |
| SS42 | TCCCAGGATTAGAAT GATTAGG | SS102 | CACCATGTTATCACATCA ATCC | 918 bp | 56 | Integration of crRNA to pDettLb-Cas12a vector; successful Gateway reaction |
| LbCas12a_F_2 | GAAGAGAGCTGAAG ATTACAAGGG | LbCas12a_R_2 | TGATCACGTTCCACTCGC CG | 953 | 55 | pDett_LbCas12a T-DNA specific |
| Spec_F_2 | CCAGTCGGGCGGCG AGTTCC | Spec_R_2 | ACGCAGCAGGGCAGTCG CCC | 576 | 60 | pDett_LbCas12a vector backbone specific |

| | | | | | | |
|---|---|---|---|---|---|---|
| pDONR201_F | TCGCGTTAACGCTAGCATGGATCTC | pDONR201_R | GTAACATCAGAGATTTTGAGACAC | 2,490 | 58 | Amplification of pDONR vector flanking the attachment sites |
| 29-Pro35S_Seq | GATGTGATATCTCCACTGACG | 28-pK7WG_R (T35S) | TTGCGGACTCTAGCATGG | 1943 | 55 | pK7WG2 T-DNA specific |
| 400_600_OE_F | GGGGACAAGTTTGTACAAAAAAGCAGGCTTAAAAATGAAGATCACTAACTTTCTAGTGC | 400_600_OE_R | GGGGACCACTTTGTACAAGAAAGCTGGGTATTAAGCGTAATCTGGAACATCGTATGGGTACATCATTTTATACTCGATTTTAGAAGC | 600 | 58 | *Potri.019G124400* and *Potri.019G124600* cDNA amplification along with an adapter for pDONR201 vector |
| 53200_OE_F | GGGGACAAGTTTGTACAAAAAAGCAGGCTTAAAAATGAATTATCCAATGTTATTGYTGTGC | 53200_OE_R | GGGGACCACTTTGTACAAGAAAGCTGGGTATTAAGCGTAATCTGGAACATCGTATGGGTACGAGTTTTTCTTCAAAGCTTTCTTG | 773 | 58 | *Potri.017G153200* cDNA amplification along with an adapter for pDONR201 vector |

**Additional Table S2:** List of primers, their annealing temperature, and their primer efficiencies for RT-qPCR.

| Primer Name Forward | Sequence (5' → 3') | Primer Name Reverse | Sequence (5' → 3') | Amplicon size (bp) | Annealing Temp(°C) | Primer Efficiency (%) | Dilution for qRT-PCR[1] | Targeted gene ID |
|---|---|---|---|---|---|---|---|---|
| Actin9_F | TGGTGGTTCCACTATGT TCC | Actin9_R | TGGAAATCCACATCTGCT GG | 176 | 50 | 99 | 1 to 90 | *Potri.001G309500* |
| AlOxS_2_F | GTCTAAAAGAAAACATG TCC | AlOxS_2_R | AAATAAAACAGAAACCA CAC | 68 | 50 | 102 | 1 to 90 | *Potra002586g19438* |
| UBQ_L_F | TGAGGCTTAGGGGAGG AACT | UBQ_L_R | TGAGGCTTAGGGGAGGA ACT | 195 | 55 | 98 | 1 to 30 | *Potri.005G198700* |
| NPR1_2_F | AGCCTTTTACAAACCCA GAAATTGC | NPR1_2_R | TAAAATGCATCTGTGAAC TGGAACC | 152 | 55 | 94 | 1 to 30 | *Potri.006G148100* |
| KTI_4_F_2 | GACACCTTCGGTGATGA GGTTAAAG | KTI_4_R_4 | GTCCACATTCAGATTACT GTCTTCA | 204 | 55 | 93 | 1 to 30 | *Potri.019G124600* |
| KTI_3_F | ATGAAGATGACTAACTT TCTAGTGCA | KTI_3_R_2 | GTCCACATTAAGATTACT GTCTTCG | 300 | 50 | 102 | 1 to 90 | *Potri.019G124400* |
| KTI_5_F | GATGGTGAAAAGCTTGT AGCAGG | KTI_5_R_2 | CCTTTCTTGTCGTCGGCA | 50 | 50 | 103 | 1 to 9 | *Potri.017G153200* |
| A_Ref2_F | ATCGTTCCAAGTCAAGT ATGTG | A_Ref2_R | TCAAGGGAGCAACTTTAC AG | 93 | 50 | 101 | 1 to 9 | *Potri.015G001600* |
| C_Ref1_F | GCAATGTGAGGAGTTTA GGG | C_Ref1_R | TATTAAATGTCTGTGCTGT AGTGTG | 86 | 50 | 95 | 1 to 9 | *Potri.012G141400* |
| PtKTIup1_RT_F | GTGTCCAACTACAAGCT CCT | PtKTIup1_RT_R | TCAAAGCCAGGTATCTAA TCCC | 117 | 50 | 98 | 1 to 9 | *Potri.019G088200* |
| PtERF1_RT_F | TTCCAGAGCAATTCACT TCCT | PtERF1_RT_R | CACTTCCTCTTCCTTAATT CCAC | 136 | 50 | 90 | 1 to 9 | *Potri.T131500* |

[1]The dilution of cDNA to be performed on stock cDNA synthesized by the RevertAid first strand cDNA synthesis kit (Thermo Fisher Scientific).

**Additional Table S3**: Gene identity numbers for *P. trichocarpa* (Potri) and *P. tremula* (Potra). The annotations were downloaded from Plantgenie (https://plantgenie.org/, accessed: 25.05.2025). The gene IDs are associated with GO:0004866: endopeptidase inhibitor activity and have the pfam_description: PF00197-Trypsin and protease inhibitor.

| Potri_ID | chromosome name | atg_id | Potra_gene_id | description |
|---|---|---|---|---|
| Potri.001G309900 | Chr01 | AT1G17860 | Potra2n1c2691 | similar to truncated Kunitz trypsin inhibitor. [ORG:Glycine max]; [ co-ortholog (1of6) of AAK20290] |
| Potri.003G097900 | Chr03 | | Potra2n3c7636 | |
| Potri.004G000400 | Chr04 | AT1G73260 | Potra2n4c8413 | similar to truncated Kunitz trypsin inhibitor. [ORG:Glycine max]; [ co-ortholog (6of6) of AAK20290] |
| Potri.004G067600 | Chr04 | AT1G17860 | Potra2n4c9034 | |
| Potri.004G067800 | Chr04 | AT1G17860 | Potra2n4c9037 | similar to trypsin protein inhibitor 3. [ORG:Cicer arietinum]; [ co-ortholog (1of3) of S19190, BAD04942, BAD04939, JC7311, AAB21123, B45588, P09941, P25272, P83594, JQ0968, AAB23464, 1AVX_B, TISYB, AAB26177, CAH61462, P32733, P83051, AAA32618, AAT45474] |
| Potri.004G067900 | Chr04 | AT1G17860 | Potra2n4c9039 | similar to trypsin protein inhibitor 3. [ORG:Cicer arietinum]; [ co-ortholog (2of3) of S19190, BAD04942, BAD04939, JC7311, AAB21123, B45588, P09941, P25272, P83594, JQ0968, AAB23464, 1AVX_B, TISYB, AAB26177, CAH61462, P32733, P83051, AAA32618, AAT45474] |
| Potri.007G111500 | Chr07 | AT1G73260 | Potra2n416s35592 | |
| Potri.007G111600 | Chr07 | AT1G17860 | Potra2n416s35593 | |
| Potri.007G111700 | Chr07 | | Potra2n416s35593 | similar to truncated Kunitz trypsin inhibitor. [ORG:Glycine max]; [ co-ortholog (4of6) of AAK20290] |
| Potri.007G111800 | Chr07 | AT1G73260 | Potra2n416s35592 | |
| Potri.010G007500 | Chr10 | | Potra2n156s34746 | |
| Potri.010G007700 | Chr10 | | Potra2n156s34746 | |
| Potri.010G007800 | Chr10 | | Potra2n156s34746 | |
| Potri.010G007900 | Chr10 | | Potra2n156s34746 | |
| Potri.017G153200 | Chr17 | AT1G17860 | Potra2n17c30597 | similar to trypsin protein inhibitor 3. [ORG:Cicer arietinum]; [ co-ortholog (3of3) of S19190, BAD04942, BAD04939, JC7311, AAB21123, B45588, P09941, P25272, P83594, JQ0968, AAB23464, 1AVX_B, TISYB, AAB26177, CAH61462, P32733, P83051, AAA32618, AAT45474] |
| Potri.017G153300 | Chr17 | AT1G17860 | Potra2n17c30594 | |

| | | | | |
|---|---|---|---|---|
| Potri.017G153400 | Chr17 | AT1G17860 | Potra2n17c30594 | |
| Potri.017G153500 | Chr17 | AT1G17860 | Potra2n17c30594 | |
| Potri.017G153600 | Chr17 | AT1G17860 | Potra2n17c30594 | |
| Potri.019G006900 | Chr19 | AT1G73260 | Potra2n19c34329 | |
| Potri.019G010900 | Chr19 | AT1G17860 | Potra2n19c34330 | |
| Potri.019G011000 | Chr19 | AT1G73260 | Potra2n19c34325 | similar to truncated Kunitz trypsin inhibitor. [ORG:Glycine max]; [ co-ortholog (2of6) of AAK20290] |
| Potri.019G088200 | Chr19 | AT1G17860 | Potra2n19c33577 | |
| Potri.019G121900 | Chr19 | | Potra2n19c33329 | |
| Potri.019G122100 | Chr19 | | Potra2n19c33329 | |
| Potri.019G124400 | Chr19 | | Potra2n19c33329 | |
| Potri.019G124500 | Chr19 | | Potra2n19c33329 | |
| Potri.019G124600 | Chr19 | | Potra2n19c33329 | |
| Potri.019G124700 | Chr19 | | Potra2n19c33329 | |
| Potri.T029100 | scaffold_31 | | Potra2n156s34746 | |

**Additional Table S4**: *In-silico* prediction of the subcellular localization of the candidate KTIs according to different bioinformatics tools. The highest probabilities are highlighted in green.

| | Potri.019G124600 | Potri.019G124400 | Potri.017G153200 | Potri.019G124600 | Potri.019G124400 | Potri.017G153200 |
|---|---|---|---|---|---|---|
| **Membrane association** | **Probability** | **Probability** | **Probability** | | | |
| **Peripheral** | 0.238 | 0.243 | 0.124 | | | |
| **Transmembrane** | 0.042 | 0.04 | 0.048 | | | |
| **Lipid anchor** | 0.039 | 0.039 | 0.052 | | | |
| **Soluble** | 0.891 | 0.895 | 0.898 | | | |
| **Localization** | **Probability** | **Probability** | **Probability** | **Score** | **Score** | **Score** |
| **Predicted signals** | Signal peptide | Signal peptide | Signal peptide | | | |
| **Cytoplasm** | 0.0706 | 0.0605 | 0.1039 | | | 1 |
| **Nucleus** | 0.1476 | 0.1699 | 0.1532 | | | |
| **Extracellular** | 0.7673 | 0.7797 | 0.8694 | 4 | 4 | 11 |
| **Cell membrane** | 0.0951 | 0.0838 | 0.1277 | | | |
| **Mitochondrion** | 0.0098 | 0.0102 | 0.0075 | | 1 | 1 |
| **Plastid** | 0.1589 | 0.2342 | 0.015 | 8 | 5 | 1 |
| **Endoplasmic reticulum** | 0.3958 | 0.3046 | 0.2826 | 1.5 | | |
| **Lysosome/Vacuole** | 0.2562 | 0.2135 | 0.244 | 1.5 | 4 | |
| **Golgi apparatus** | 0.0709 | 0.0569 | 0.0706 | | 1 | |
| **Peroxisome** | 0.0023 | 0.0038 | 0.0031 | | | |

https://services.healthtech.dtu.dk/services/DeepLoc-2.1/

https://wolfpsort.hgc.jp/

**Additional Table S5:** Description of transformed and surviving mutant lines of *Kunitz Trypsin Inhibitor* in *Populus* x *canescens.*

| Line number | Vector used | Purpose | Gene target | Deletion (bp) [1] |
|---|---|---|---|---|
| 1_8 | pDett-Lb-Cas12a_400_600 | loss-of-function | *Potri.019G124400* and *Potri.019G124600* | 8 |
| 1_22 | pDett-Lb-Cas12a_400_600 | loss-of-function | *Potri.019G124400* and *Potri.019G124600* | 6 |
| 1_28 | pDett-Lb-Cas12a_400_600 | loss-of-function | *Potri.019G124400* and *Potri.019G124600* | 50 |
| 1_32 | pDett-Lb-Cas12a_400_600 | loss-of-function | *Potri.019G124400* and *Potri.019G124600* | 14 |
| 1_37 | pDett-Lb-Cas12a_400_600 | loss-of-function | *Potri.019G124400* and *Potri.019G124600* | 10 |
| 1_41 | pDett-Lb-Cas12a_400_600 | loss-of-function | *Potri.019G124400* and *Potri.019G124600* | 9 |
| 1_52 | pDett-Lb-Cas12a_400_600 | loss-of-function | *Potri.019G124400* and *Potri.019G124600* | 50 |
| 1_54 | pDett-Lb-Cas12a_400_600 | loss-of-function | *Potri.019G124400* and *Potri.019G124600* | 50 |
| 2_18 | pDett-Lb-Cas12a_53200 | loss-of-function | *Potri.017G153200* | 7 |
| 2_29 | pDett-Lb-Cas12a_53200 | loss-of-function | *Potri.017G153200* | 10 |
| 2_43 | pDett-Lb-Cas12a_53200 | loss-of-function | *Potri.017G153200* | 5 |
| 2_45 | pDett-Lb-Cas12a_53200 | loss-of-function | *Potri.017G153200* | 5 |
| 3_7 | pDett_LbCas12a_pEn-RZ-Lb_chimera | Ubiquitin_Promoter_EmptyVector | Empty vector control | N/A |
| 3_15 | pDett_LbCas12a_pEn-RZ-Lb_chimera | Ubiquitin_Promoter_EmptyVector | Empty vector control | N/A |
| 3_16 | pDett_LbCas12a_pEn-RZ-Lb_chimera | Ubiquitin_Promoter_EmptyVector | Empty vector control | N/A |
| 3_20 | pDett_LbCas12a_pEn-RZ-Lb_chimera | Ubiquitin_Promoter_EmptyVector | Empty vector control | N/A |

[1] All deletions were induced in exons. Length of deletions determined via Sanger sequencing.

**Additional Table S6:** Consequences of CRISPR-Cas12a editing events observed in mutant lines.

| Plant ID | Gene | CRISPR-Cas12a system | Predicted truncated amino acid length | No. premature stop codons |
|---|---|---|---|---|
| WT KTI_400* | *KTI_400* | MKITNFLVHSFLLFAFTATSIFPRAV HAGAVIDAFGDEVKAGDRYIIGAAS NDFAITATSPIICNSDVVFSPMSNGL PVIFSKVVESNDSVINEDSYLNVDF DAPSCRMAGVSTMWKIELRPTARG FVVTTGGVAGLNRFTITKYGDGTN LYQLSYCPISEPICECSCVPLGNVVN RLAPSTIPFPVVFIPADRASKIEYKM M* | 204 (full-length) | |
| WT KTI_600* | *KTI_600* | MKITNFLVLSFLLFAFTATSIFPRAVH AAAVIDTFGDEVRAGDRYIIGAASN DFAVTATSPIICNSDVVFSPMSTGLP VIFSKVVESNDSVINEDSNLNVDFD AATCRMAGVSTMWKIEMRPTARG FVVTTGGVAGLNRFTITKYEGGNN LYQLSYCPISEPICKCSCVPLGKVVN RLAPSTVPFPVVFIPADRASKIEYK MM* | 204 (full-length) | |
| *kti4+600_1_52* | *KTI_400* | MKITNFLVHSFLLFAFTATSIFPRAV HAGAVIDAFGDEVKAGDRYIIGARC CVFSDEQWTPCNIFKSCRIQRQCHQ RRQLSECGL*CTLM*DGGRLNHVE D*IEANSARIRCDHRRCCWIESVYD HQVWRWYQFVSAFLLSNFRTHM* MLMRPTRQRCQSLGSQYHPFSCCV YTSR*SF*NRV*NDV | 86; 91; 102; 149; 178; 181; 185 | 7 |
| | *KTI_600* | MKITNFLVLSFLLFAFTATSIFPRAVH AAAVIDTFGDEVRAGDRYIIGARCC VFSDEHWTPSNIFKSCRIQRQCHQR RQ*SECGL*CSHM*NGGRINHVED* NEANSARIRCDHRRCCWIESVYDH QV*RW**FVSAFLLSNFGTHM*ML MRPTRQSCQSLGSQYRPFSSCVYTS R*SF*NRV*NDV | 81; 87; 92; 103; 130; 133; 134; 149; 178; 181; 185 | 11 |

| | | | | |
|---|---|---|---|---|
| kti4+600_1_28 | *KTI_400* | MKITNFLVHSFLLFAFTATSIFPRAVHAGAVIDAFGDEVKAGDRYIIGARCCVFSDEQWTPCNIFKSCRIQRQCHQRRQLSECGL*CTLM*DGGRLNHVED*IEANSARIRCDHRRCCWIESVYDHQVWRWYQFVSAFLLSNFRTHM*MLMRPTRQRCQSLGSQYHPFSCCVYTSR*SF*NRV*NDV | 86; 91; 102; 149; 178; 181; 185 | 7 |
| | *KTI_600* | MKITNFLVLSFLLFAFTATSIFPRAVHAAAVIDTFGDEVRAGDRYIIGARCCVFSDEHWTPSNIFKSCRIQRQCHQRRQ*SECGL*CSHM*NGGRINHVED*NEANSARIRCDHRRCCWIESVYDHQV*RW**FVSAFLLSNFGTHM*MLMRPTRQSCQSLGSQYRPFSSCVYTSR*SF*NRV*NDV | 81; 87; 92; 103; 130; 133; 134; 149; 178; 181; 185 | 11 |
| WT KTI_53200* | *KTI_53200* | MNYPMLLLCLLLLAFACTKQSIAAAEPVLDIDGEKLVAGTEYYILPVFRGRGGGITMASNKTSCPLAVVQDRLEVSKGVPLTFTPAADDKKGVILVSADLNIKFLAKTTCPQSTVWKIIKSSNSKVQWFVSTGGVEGNPGFNTVTNWFQIEKADDDYKLVFCPTKVCNCGVLCRDIGIYIEDNGTRTLSLSDALLPFKVQFKKALKKNS* | 210 (full-length) | |
| kti53200_2_18 | *KTI_53200* | MNYPMLLLCLLLLAFACTKQSIAAAEPVLDIDGEKLVAGTEYYILPVFRGRGGGITMASNKTSCPLAVVQDRLEVSKGVPLTFTPAADKV*SLFLLILTSSF*RRQHVPNQLYGRL*SLRTRRYNGLCQLVGLKEILVLIR*PTGSRLRKLMMTTSLFSVLLKFVTVEFYAGILGFILRIMGLEHCLSVMHYYLSKSSSRKL*RKTR | 92; 104; 118; 143; 204 | 5 |
| kti53200_2_29 | *KTI_53200* | MNYPMLLLCLLLLAFACTKQSIAAAEPVLDIDGEKLVAGTEYYILPVFRGRGGGITMASNKTSCPLAVVQDRLEVSKGVPLTFTPAADE*SLFLLILTSSF*RRQHVPNQLYGRL*SLRTRRYNGLCQLVGLKEILVLIR*PTGSRLRKLMMTTSLFSVLLKFVTVEFYAGILGFILRIMGLEHCLSVMHYYLSKSSSRKL*RKTR | 91; 103; 117; 142; 203 | 5 |

*WT amino acid sequences from *KTI_400, KTI_600* and *KTI_53200.*
Red-colored amino acid residue denotes signal peptide; green denotes Kunitz. Motif; blue reactive loop and yellow disulfide bridge. *denotes the stop codon.

**Additional Table S7**: Gas exchange and growth of wildtype and transgenic poplar lines (*P*. x *canescens*). N = 4 ± SE, WT: n= 8. Different letters indicate significant differences of means ($p < 0.05$, Tukey test).

| Line | Count | Transpiration mol/m² | | SE | Gs mmol/m²s1 | SE | | Stem height cm | | SE mm | Stem diameter | | SE | | Root-to-Shoot | SE |
|---|---|---|---|---|---|---|---|---|---|---|---|---|---|---|---|---|
| WT | 8 | 4.77 | 0.18 | AB | 0.228 | 0.013 | A | 57.31 | 2.34 | A | 6.88 | 0.22 | A | 0.77 | 0.12 | DE |
| OXEV_3 | 4 | 4.74 | 0.14 | AB | 0.218 | 0.011 | A | 63.63 | 1.53 | AB | 6.92 | 0.18 | A | 0.48 | 0.02 | ABCDE |
| OX532_8 | 4 | 4.32 | 0.24 | AB | 0.198 | 0.008 | A | 68.68 | 2.69 | AB | 7.45 | 0.32 | A | 0.41 | 0.02 | ABC |
| OX532_4 | 4 | 4.25 | 0.19 | AB | 0.188 | 0.010 | A | 66.03 | 1.24 | AB | 7.80 | 0.24 | A | 0.45 | 0.02 | ABCD |
| OX532_10 | 4 | 4.36 | 0.08 | AB | 0.200 | 0.007 | A | 60.55 | 2.76 | AB | 6.98 | 0.67 | A | 0.37 | 0.04 | AB |
| OX400_5 | 4 | 4.18 | 0.43 | AB | 0.193 | 0.027 | A | 64.88 | 0.63 | AB | 6.55 | 0.29 | A | 0.52 | 0.03 | ABCDE |
| OX400_4 | 4 | 3.53 | 0.22 | A | 0.160 | 0.011 | A | 63.10 | 1.73 | AB | 6.80 | 0.16 | A | 0.45 | 0.04 | ABCD |
| OX400_15 | 4 | 5.21 | 0.38 | B | 0.243 | 0.023 | A | 55.43 | 2.75 | A | 6.48 | 0.31 | A | 0.33 | 0.04 | A |
| OX400_14 | 4 | 3.98 | 0.30 | AB | 0.178 | 0.019 | A | 59.80 | 5.52 | AB | 7.25 | 0.28 | A | 0.40 | 0.03 | ABC |
| kti532_43 | 4 | 4.00 | 0.18 | AB | 0.180 | 0.004 | A | 65.60 | 2.23 | AB | 7.16 | 0.36 | A | 0.44 | 0.04 | ABC |
| kti532_29 | 4 | 3.67 | 0.33 | A | 0.160 | 0.024 | A | 66.13 | 3.76 | AB | 7.31 | 0.35 | A | 0.47 | 0.03 | ABCDE |
| kti532_18 | 4 | 3.87 | 0.23 | AB | 0.168 | 0.015 | A | 64.98 | 0.80 | AB | 6.98 | 0.07 | A | 0.51 | 0.07 | ABCDE |
| kti400_41 | 4 | 3.68 | 0.37 | A | 0.158 | 0.022 | A | 68.18 | 2.86 | AB | 7.79 | 0.25 | A | 0.73 | 0.11 | BCDE |
| kti400_32 | 4 | 3.60 | 0.38 | A | 0.160 | 0.017 | A | 71.43 | 1.88 | B | 7.82 | 0.36 | A | 0.85 | 0.05 | E |
| kti400_28 | 4 | 4.33 | 0.29 | AB | 0.193 | 0.020 | A | 65.60 | 2.16 | AB | 7.91 | 0.32 | A | 0.78 | 0.04 | CDE |
| koev_16 | 4 | 3.94 | 0.33 | AB | 0.178 | 0.018 | A | 68.73 | 2.47 | AB | 7.99 | 0.41 | A | 0.48 | 0.03 | ABCDE |
| koev_15 | 4 | 3.82 | 0.29 | AB | 0.170 | 0.019 | A | 70.50 | 0.74 | B | 7.66 | 0.19 | A | 0.41 | 0.04 | ABC |
| Total | 72 | 4.13 | 0.11 | | 0.186 | 0.006 | | 64.74 | 1.07 | | 7.28 | 0.12 | | 0.52 | 0.04 | |
