## Additional_materials for "Divergent functions of three Kunitz trypsin inhibitor (KTI) proteins in herbivore defense in poplar": Additional_methods.pdf

**for**

### Transformation protocol for protease inhibitors in *Populus* x *canescens*.

Here we report all details of the transformation strategy to produce CRISPR-Cas12a knock mutants and *p35S* overexpression lines of *P.* x *canescens* for Kunitz Trypsin Inhibitor (KTIs) proteins (reproduced from the thesis of Das, 2024).

#### 1.1.1 Designing target sites for the CRISPR-Cas12a system

The PAM site (5'-TTTV-3') was searched within the gDNA of the candidate *KTI* sequences. The designing criteria of target sites (24-nucleotide sequence) were as follows: (1) the 24-nucleotide sequence must lie on the 3' end of the PAM (5'-TTTV-3') sequence; (2) the target site sequence must lie on a cDNA; (3) the target site sequence must not have any off-targets (ambiguous hit to any other region of the genome); (4) the sequence should not form any secondary structures; (5) the GC content should be higher than 50 % and (6) the target sequence should not have more than five consecutive T nucleotides. Following all these criteria, the target sites were identified and oligonucleotides were ordered from Microsynth Seqlab.

### 1.2 Molecular cloning

#### 1.2.1 Preparation of electro-competent *Escherichia coli* and *Agrobacterium tumefaciens* cells

A stock culture of *Escherichia coli,* DH5α (for all other transformations) and DB3.1 (for Gateway reactions) was used to prepare the competent cells. *E. coli* strain used for individual transformation is mentioned along with the protocol. This protocol was adapted from (Sharma and Schimke, 1996). A cryo-stock culture of *E. coli* was streaked on an LB-media plate without antibiotics (Table M1) and incubated overnight at 37 °C. With a sterile toothpick, a small culture of *E. coli* was picked and incubated further in glass tubes containing 5 mL of liquid LB (without antibiotics). Tubes were incubated overnight at 37 °C with constant shaking at 180 rpm in a shaker (ZWYR-240, Labwit Manufacturer, Shanghai ZHICHENG Analytical Instruments Manufacturing Co., Shanghai, China). One mL of the overnight culture was poured into 500 mL of fresh liquid LB medium in 2 L baffled flask. The flask was sealed with a cotton plug and was further incubated under constant shaking at 180 rpm at 37 °C (ZWYR-240 shaker Labwit Manufacturer) until the culture reached an $OD_{600}$ of 0.5 – 0.8. $OD_{600.}$ The OD values were measured in a photometer (Biophotometer 6131, Eppendorf AG) by pipetting 1 mL of bacterial culture to a UV semi-micro cuvette (Sarstedt AK & Co.KG, Nümbrecht, Germany) and fresh liquid LB medium as the blank. The bacterial culture was divided into two 250 mL capped sterile centrifuge bottles and was incubated in ice for 30 min, with occasional stirring. The cultures in centrifuge bottles were centrifuged (J2-HS, Beckman Coulter, California, Germany) at 5000 x *g*, and pre-cooled at 4 °C for 15 min. The supernatant was discarded, and the pellet was resuspended in ice-cold sterile demineralized water. This process was repeated three times, and the final pellet was resuspended in autoclaved (at 2.2 bar, 121 °C for 20 min) 10 % glycerol (dissolved in $ddH_2O$, v:v). The final culture was transferred into 15 mL Falcon tubes (Sarstedt AG & Co. KG) and further centrifuged (5000 x *g,* pre-cooled at 4 °C for 15 min). The supernatant was discarded, and the new pellet was dissolved in 700 µL of 10 % glycerol. Aliquots of 50 µL were frozen in liquid nitrogen and then stored at -80 °C. All containers and flasks were sterilized for the preparation of the competent cells. All processes, such as streaking, transfer of cultures, etc., were performed under the sterile bench (Thermo Fisher Scientific). No antibiotics were used for the culture of *E. coli* strain DH5α and DB3.1 since they are susceptible to ampicillin, spectinomycin, and kanamycin. The usage of specific antibiotics in specific media has been described along with the methodology.

**Table M1**: LB and YEB media preparation for *Escherichia coli* and *Agrobacterium tumefaciens* culture. All media were prepared in either 1 L or 500 mL glass bottles and autoclaved at 2.2 bar, 121 °C for 20 min (6x6x6 HST, Zirbus technology GmbH). Solid media were poured into 10 cm petri-dishes (Greiner Bio-One International GmbH) under the sterile bench. pH of all media was adjusted to 7.0 with NaOH (Carl Roth GmbH & Co. KG). All antibiotics were sterile-filtered through a filter (Filtropur S 0.2, size 0.2 µm; Sarstedt AG & Co. KG) with a syringe under a sterile bench. The filtrates were stored as aliquots at -20 °C. In any cloning steps or preparation of competent cells, any antibiotics used will be stated along with the described methodology.

| **LB Solid** | | |
|---|---|---|
| **Chemical** | **Final concentration (g/L)** | **Company** |
| Tryptone | 10 | Carl Roth GmbH & Co. KG |
| Bacto Yeast extract | 5 | Becton, Dickinson and Company, New Jersey, USA |
| NaCl | 10 | Carl Roth GmbH & Co. KG |

| | | |
|---|---|---|
| Micro-Agar | 20 | Duchefa Biochemie B.V. |
| **LB liquid** | | |
| **Chemical** | **Final concentration (g/L)** | **Company** |
| Tryptone | 10 | |
| Bacto Yeast extract | 5 | Same as above |
| NaCl | 10 | |
| **YEB solid** | | |
| **Chemical** | **Final concentration (g/L)** | **Company** |
| Bacto Beef extract | 5 | Becton, Dickinson and Company |
| Bacto Yeast extract | 1 | Becton, Dickinson and Company |
| Bacto Peptone | 5 | Becton, Dickinson and Company |
| Sucrose | 5 | Carl Roth GmbH & Co. KG |
| $MgSO_4.7H_2O$ | 0.3 | Carl Roth GmbH & Co. KG |
| Micro-Agar | 20 | Duchefa Biochemie B.V. |
| **YEB liquid** | | |
| **Chemical** | **Final concentration (g/L)** | **Company** |
| Bacto Beef extract | 5 | |
| Bacto Yeast extract | 1 | |
| Bacto Peptone | 5 | Same as above |
| Sucrose | 5 | |
| $MgSO_4.7H_2O$ | 0.3 | |
| **Antibiotics** | | |
| Ampicillin | 50 mg/L | Duchefa Biochemie B.V. |
| Spectinomycin | 50 mg/L | Duchefa Biochemie B.V. |
| Kanamycin | 50 mg/L | Duchefa Biochemie B.V. |
| Gentamicin | 50 mg/L | Duchefa Biochemie B.V. |
| Rifampicin | 50 mg/L | Duchefa Biochemie B.V. |

Similarly, competent *Agrobacterium tumefaciens* cells of the strain GV3101-pMP90 were prepared. Instead of the LB-based growth media, solid and liquid YEB media were used with the antibiotics gentamicin and rifampicin (Table M1). These antibiotics were used for the

*Agrobacterium* competent cell preparations since this strain already comprises of the gentamicin and rifampicin resistance cassettes. The incubation temperatures were changed to 28 °C (ZWYR-240 shaker Labwit Manufacturer).

To test the competency of the cells, a high copy plasmid, e.g., pUC19 (100 ng), was transformed into the freshly prepared competent cells separately to *E. coli* and *Agrobacterium* cells. Post transformation, 1 mL of culture was diluted to $10^{-5}$ (no $OD_{600}$ measurement is needed here since this process is only to multiply cell culture) using liquid LB medium without antibiotics (Table M1). Hundred μL of the $10^{-5}$ diluted cells were plated by gently distributing the culture with a sterile spatula onto LB solid media plates with ampicillin (50 mg/L) (Table M1). The plates were incubated overnight at 37 °C in darkness in an incubator (shaker ZWYR-240, Labwit Manufacturer). The total number of colonies was counted, and the transformation efficiency was calculated as:

$$\frac{\text{Transformation efficieny}}{\mu\text{g of pUC19 plasmid}} = \text{Counted colony number X 10 X } 10^{-5} \text{ X 10}$$

In the equation, the first ten was applied since only 100 µl of culture was taken from a 1000 µl. Second X $10^{-5}$ because of the dilution factor. Third X10 because 100 ng of plasmid was transformed and the final efficiency to be determined was calculated for 1 µg of the transformed cells.

For bacterial cells to be stated as competent, only an efficiency in the range of $10^7$ to $10^9$ was stated to be competent (Taketo, 1988). If the efficiency was not attained, a fresh batch of competent cells was again prepared.

#### 1.2.2 Transformation of plasmids into competent cells via electroporation

Competent *E. coli* cells were thawed for 5 min in ice. 100 ng of the plasmid of interest was added, and the suspension was further incubated on ice for 10 min. The total mixture was transferred to an ice-cold 1 mm gap electro-competent cuvette (Sarstedt GmbH) and incubated for 5 min inside the cuvette. For the transformation of *E. coli* cells, electroporation was performed with the Eporator (Eppendorf AG) at 1.7 kV for 5 ms according to the instructions of Eporator (Eppendorf AG) adapted from Dower *et al.*, (1988). One mL LB medium (without antibiotic) (Table M1) was immediately pipetted to the transformed cells within the cuvette. This culture was mixed gently and transferred to a 2 mL Eppendorf microtube and continuously shaken at 180 rpm for 1 h (shaker ZWYR-240, Labwit Manufacturer). 100 μL culture was plated onto LB solid media plates with the antibiotics as described in the specific cloning protocols.

The same transformation strategy was applied for *Agrobacterium* cells except for the electroporation step. For the *Agrobacterium* cells, 2 mm gap electro-competent cuvettes (Sarstedt GmbH) were used. The cells were transformed at 2.0 kV for 5 ms. Transformed cells were resuspended in 1 mL of YEB media. 100 µL culture was plated onto YEB media (Table M1) plates as described in the specific cloning protocol.

#### 1.2.3 Colony PCR

Colonies from transformed cells plated overnight on LB plates (see section 1.2.2) were used for colony PCR, adapted from the original protocol of Bergkessel and Guthrie (2013). A standard PCR master mixture was prepared as described by the manufacturer. Fifty µl master mixture was aliquoted in 8-strip PCR wells (Sarstedt AG & Co. KG). Under a sterile bench, single colonies were picked with sterile toothpicks and dipped separately into a well with the PCR mixtures for 2 min. The same colonies were also incubated individually in 3 mL of LB liquid culture (with vector-specific antibiotics, Figure M1, M2 and M3), by picking the respective colonies with fresh sterile toothpicks. The culture was shaken (shaker ZWYR-240, Labwit Manufacturer) overnight in ambient light conditions at 180 rpm. This culture was used for the extraction of the plasmid (see section 1.2.4 for the methodology).

#### 1.2.4 Plasmid extraction

The protocol for high copy plasmid extraction was followed as per the instructions of the innuPREP Plasmid kit (Analytik Jena). The overnight 3 mL LB liquid culture which was prepared in parallel with colony PCR (see section 1.2.3) was used for the extraction of the plasmid. The LB media comprised of the vector-specific antibiotics (Figure M1, M2 and M3). The steps of plasmid extraction were followed exactly as described by the manufacturer (Analytik Jena). The extracted plasmid was validated with Sanger sequencing. Sanger sequencing services were used from Microsynth Seqlab (Göttingen). Sample preparation for Sanger sequencing was performed according to the instruction of the company (https://www.microsynth.com/sample-requirements.html; Accessed on 28th August, 2023).

#### 1.2.5 Molecular cloning for the CRISPR-Cas12a system

The CRISPR-Cas12a system used in this study was based on the Cas protein of the *Lachnospiraceae bacterium ND2006* (Schindele and Puchta, 2020). The Gateway-compatible cloning plasmid sets, pDettLbCas12a and pEnRZ-Lb-Chimera (Figure M1a,b), were received as a gift from the lab of Prof. Dr. Holger Puchta (KIT, Karlsruhe, Germany). The cloning protocol developed by the Puchta lab (Merker et al., 2020) was applied in this study with minor modifications.

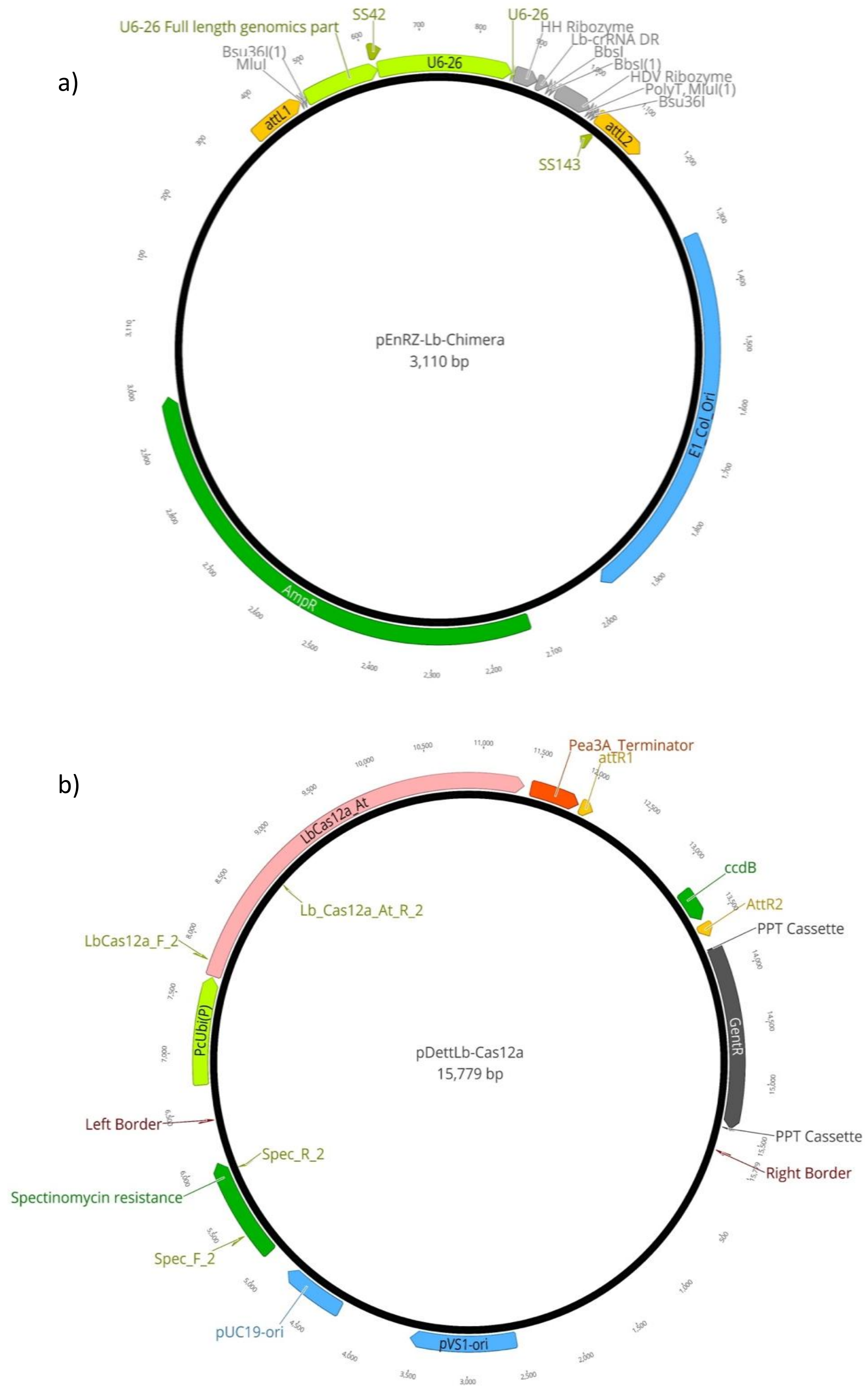


**Figure M1:** Binary vector systems of CRISPR-Cas12a. The vector sets a) pEnRZ-lb-Chimera and b) pDettLb-Cas12a were used for the CRISPR-Cas12a system-based knock-out system (Schindele and Puchta, 2020).

All major steps of cloning have been briefed through a flowchart for better understanding (Figure M4). The first step of cloning starts with the annealing of the forward and reverse oligonucleotides targeting the same gene. This was done by adding 1 µL of forward and reverse (100 µM) each to 48 µL of dd$H_2O$ and incubating the mixture at 95 °C for 5 min, followed by cooling down of the mixture for 20 min at room temperature. This process was applied to the ordered target oligonucleotides (from Microsynth Seqlab) “Target_400_600” and “Target_53200” individually. The target oligonucleotides were ordered with adapter sequences (shown in Table M2) for the ligation to the target oligonucleotides to vector- pEnRZ-Lb-Chimera.

**Table M2:** Designed target sites for CRISPR-Cas12a based knock-out system. Name of the target site, its GC content and poplar gene ID it is targeted to. The red nucleotides were overhangs introduced for the *BbsI* site ligation. All target sites were designed on the 3’ site of the PAM sequence “TTTV”.

| Target Name | Gene targeted | Sequence (5' → 3') | GC % | Position at gDNA (5' → 3') (bp) |
|---|---|---|---|---|
| Target_400_600_F | *Potri.019G124400* and *Potri.019G124600* (*KTI_400* and *KTI_600*) | AGATCGATCACTGCGACTAGCCCTATCA | 54 | 223 |
| Target_400_600_R | | GGCCTGATAGGGCTAGTCGCAGTGATCG | | 246 |
| Target_53200_F | *Potri.017G153200* (*KTI_53200*) | AGATACACCAGCTGCCGACGACAAGAAA | 54 | 440 |
| Target_53200_R | | GGCCTTTCTTGTCGTCGGCAGCTGGTGT | | 463 |

The annealed target oligonucleotides were individually integrated into the crRNA vector, pEnRZ-Lb-Chimera. For this, pEnRZ-Lb-Chimera was digested with the *BbSI* restriction enzyme (BbSI-HF) by adding 17 µL pEnRZ-Lb-Chimera (50 ng), 2 µL of 10 X CutSmart buffer and 20 U of *BbSI*- HF (New England Biolabs Ltd., Massachusetts, USA) and incubated for 2 h at 37 °C in the ThermoMixer® (Eppendorf AG). The mixture was purified with the innuPREP PCR Pure kit (Analytik Jena GmbH) according to the manufacturer's instructions. The concentration of the purified product was checked in a spectrophotometer (NanoDrop™ One spectrophotometer, Thermo Fisher Scientific, Waltham, USA) and was adjusted to 5 ng. The digested vector was ligated with the target oligonucleotides by mixing 1:1 (molar ratio of vector : “target”) ratio along with 2 µL 10X T4 DNA Ligase buffer, 1 U T4 DNA Ligase, and filled up to 20 µL ddH$_2$O (Thermo Fisher Scientific). The mixture was incubated at 22 °C in the ThermoMixer® (Eppendorf AG) for 1 h and transformed into *E. coli* strain DH5α (see section

1.2.2 for the methodology). One mL culture of transformed *E. coli* was plated onto LB with Ampicillin and incubated at 37 °C (shaker ZWYR-240, Labwit Manufacturer) in darkness. The next day, colony PCR was performed with the primer sets SS42+Target_400_600_R and SS42+Target_53200_ R (Additional Table S1) to control the respective target site integration into pEnRZ-Lb-Chimera. Positive PCR colonies were used for plasmid extraction with innuPREP plasmid kit (Analytik Jena GmbH). Target oligonucleotide integration to the pEnRZ-Lb-Chimera plasmids was verified by Sanger sequencing using primer SS42 as the reference (Microsynth seqlab). The final concentration of the plasmids harboring the target oligonucleotides and the crRNA i.e., pEnRZ-Lb-Chimera+Target_400_600 and pEnRZ-Lb-Chimera+Target_53200 was measured with the NanoDrop™ One spectrophotomer (Thermo Fisher Scientific) and was normalized to 100 ng by diluting in sterile demineralized water.

Integration of the crRNA complex with *U6-26* promoter and the ligated target sites (see Figure M2a,b) to pDettLbCas12a was performed by Gateway reaction (Thermo Fisher Scientific). 2 μL of pEnRZ-Lb-Chimera+Designed Target (adjusted to 100 ng), 3 μL of pDettLbCas12a (adjusted to 50 ng), 4 μL of TE buffer (10 mM TRIS-HCl (Tris-hydroxymethyl-aminomethane-hydrochloride); 1mM EDTA dissolved in $ddH_2O$, pH to 8.0; Carl Roth GmbH) and 1 µl LR Clonase II (Gateway™ LR Clonase™ II Enzyme mix, Thermo Fisher Scientific) were mixed and incubated at room temperature for 2 h. Proteinase K (Thermo Scientific) of 0.6 U was added and the mixture was incubated at 37 °C in the ThermoMixer (Thermo Fisher Scientific) for 10 min. Five µl of the reaction mixture was transformed into *E. coli* strain DB3.1 (see section 1.2.2 for the methodology) and plated onto LB solid medium containing spectinomycin (Table M1) for overnight incubation at 37 °C in an incubator (ZWYR-240, Labwit Manufacturer). Colonies of *E. coli* were tested for successful transformation by performing colony PCR with primer sets SS42+SS102 (Additional Table S1). Positive colonies were considered for plasmid extraction (see section 1.2.4) by incubating the positive colonies overnight in LB liquid culture with spectinomycin (Table M1). Standard PCR was performed with the primer sets SS42+SS102 (Additional Table S1) and Sanger sequencing with the same primers as reference. The generated vector maps with the crRNA, Cas12a and the target oligonucleotides Target_400_600 and Target_53200 have been shown in Figure M3.

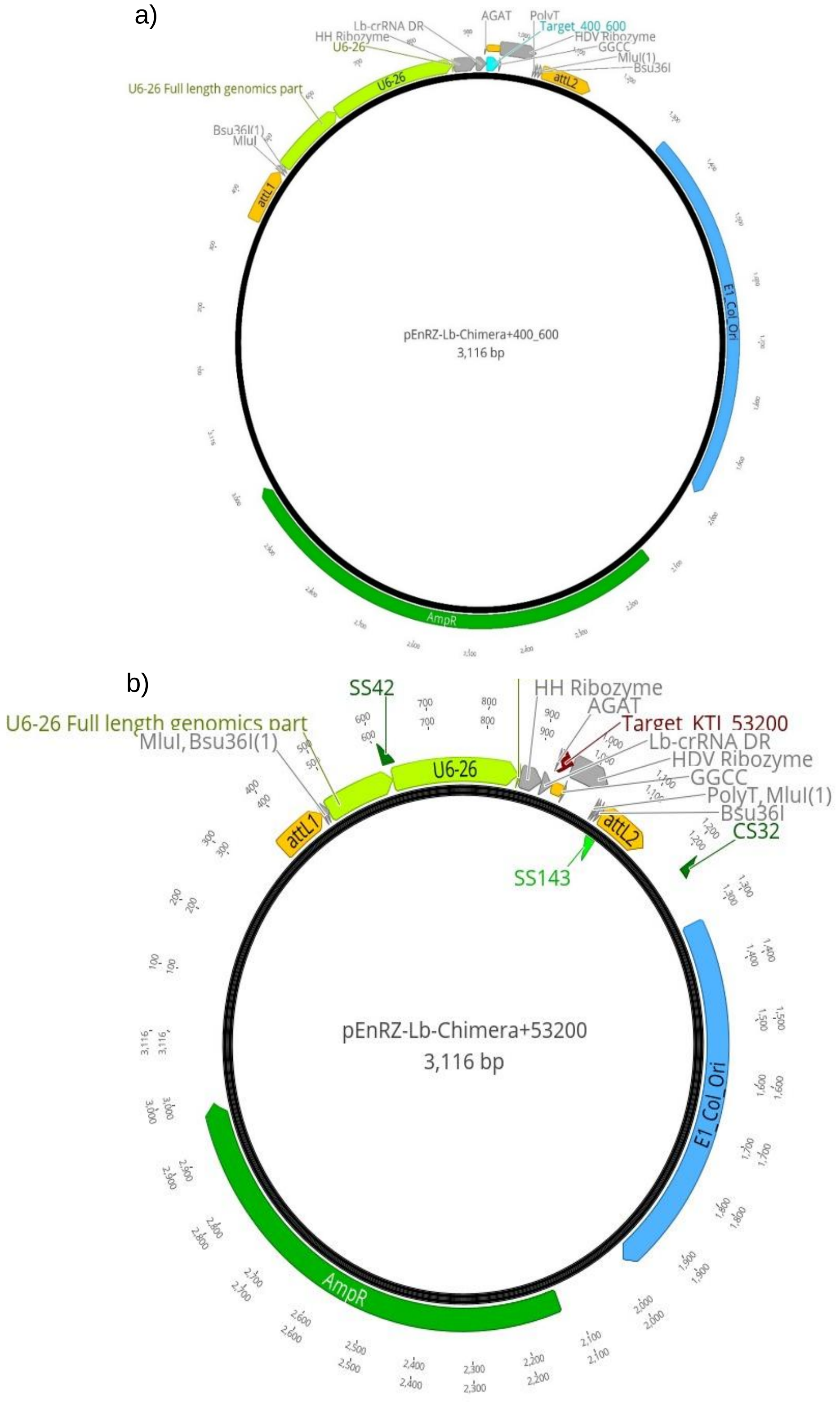


**Figure M2:** Vector maps of sgRNA vector- pEnRZ-Lb-Chimera with target sites. a) integration of Target_400_600 (blue arrow) and b) Target_53200 (red arrow) by restriction digestion with *BbsI* restriction enzyme. Further ligation of target sites with a T4 DNA ligase (Thermo Fisher Scientific, Waltham, USA). Vector maps were generated in Geneious Prime (Biomatters Ltd., New Zealand). Vector annotations on Additional Table S3.

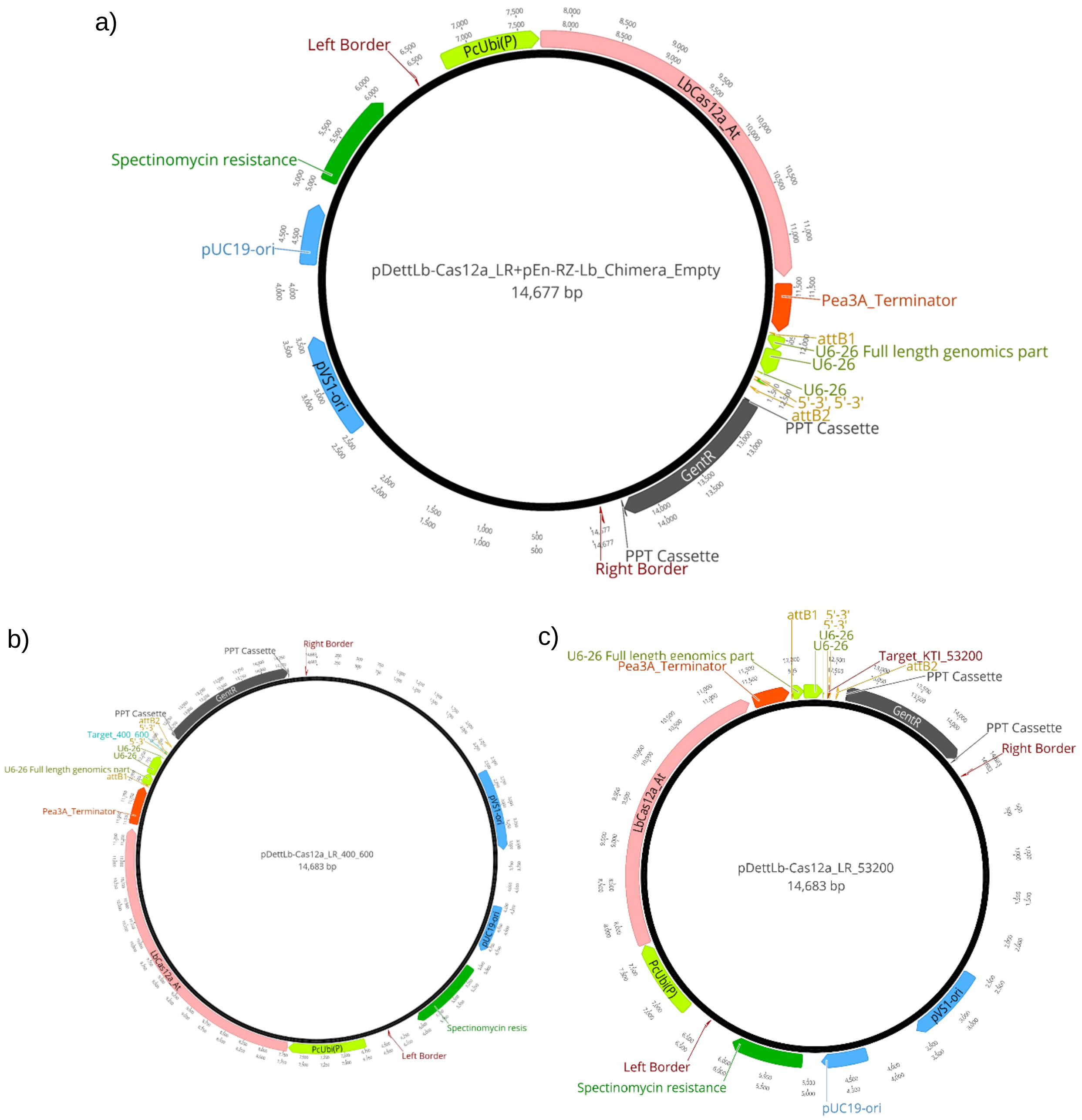


**Figure M3:** Vector maps of CRISPR-Cas12a system generated for the induced loss-of-function of *KTIs* in *Populus* x *canescens.* Final vectors were generated via the recombination of pDettLb-Cas12a with sgRNA vector- pEN-RZ-Lb_Chimera forming a) pDettLb-Cas12a+ pEN-RZ-Lb_Chimera, as empty vector control, b) Target_400_600 blue arrow) integrated into pDettLb-Cas12a targeting both *KTI_400* and *KTI_600* (double knock-out) and c) Target_53200 (red arrow) integrated into pDettLb-Cas12a targeting *KTI_53200*. Vector maps generated in Geneious Prime (Biomatters, Ltd., Auckland, New Zealand). Vector annotations on **Fehler! Verweisquelle konnte nicht gefunden werden.**.

For developing the empty vector controls, the vector sets pDett-LbCas12a+crRNA were used, but without the 24-nucleotide target sites. All transformation, plasmid isolation, and validation procedures were followed as described for the standard CRISPR-Cas12a cloning system

(Merker et al., 2020). The generated vector map pDettLb-Cas12a+ pEN-RZ-Lb_Chimera has been shown in Figure M3a.

The final vector constructs, pDettLbCas12a+Target_400_600, pDettLbCas12a+Target_53200 and empty vector control pDettLb-Cas12a+ pEN-RZ-Lb_Chimera (Figure M3) with the crRNA, target oligonucleotides, and the Cas12a protein expressed under the *UBIQUITIN* promoter were transformed into *Agrobacterium* cells (see section 1.2.2 for the methodology). The overnight colonies of the transformed *Agrobacterium* were generated on solid YEB medium containing rifampicin, gentamicin, and spectinomycin antibiotics which was incubated at 28 °C in darkness (incubator ZWYR-240, Labwit Manufacturer). The colonies were validated with colony PCR (see section 1.2.3 for the methodology) with primer sets SS42+SS102 (Additional Table S1). The positive colonies were also tested with Sanger sequencing for the validation of the transformation of the correct vector construct to the *Agrobacterium* cells. Sanger sequencing was performed using primers SS42 and SS102 as the reference (Microsynth Seqlab). The positively transformed *Agrobacterium* cells with the transformed pDettLbCas12a+Target_400_600, pDettLbCas12a+Target_53200 and empty vector control pDettLb-Cas12a+ pEN-RZ-Lb_Chimera were preserved as cryo-stocks. It was prepared by adding 500 µl of the *Agrobacterium* culture and 500 µl of 50 % sterile glycerol to the 2 mL Eppendorf reaction tubes and freezing the culture aliquots with liquid nitrogen and finally storage at -80 °C.

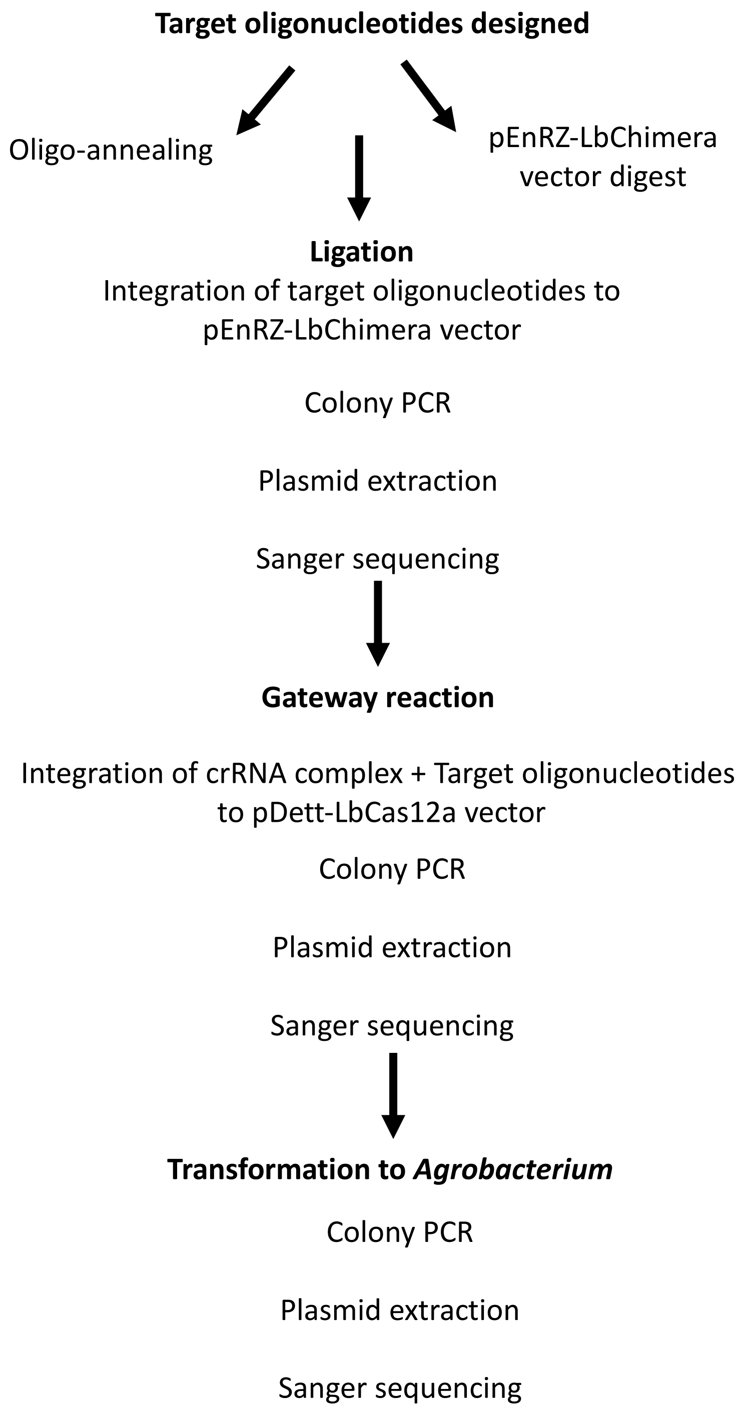


**Figure M4:** Schematic overview of the molecular cloning of CRISPR-Cas12a system.

#### 1.2.6 Molecular cloning of *p35S*-based over-expression system.

Over-expression of candidate *KTIs* was performed under the influence of a *p35S* promoter using the binary vector sets pDONR201 (entry vector; Invitrogen Life technologies) and pK7WG2 (destination vector; Karimi *et al.*, 2002) (Figure M5a,b). These vectors are Gateway compatible and were thus cloned and modified via the Gateway cloning system (Hartley 2000; Katzen 2007). The overview of the procedure of cloning has been shown in Figure M4.

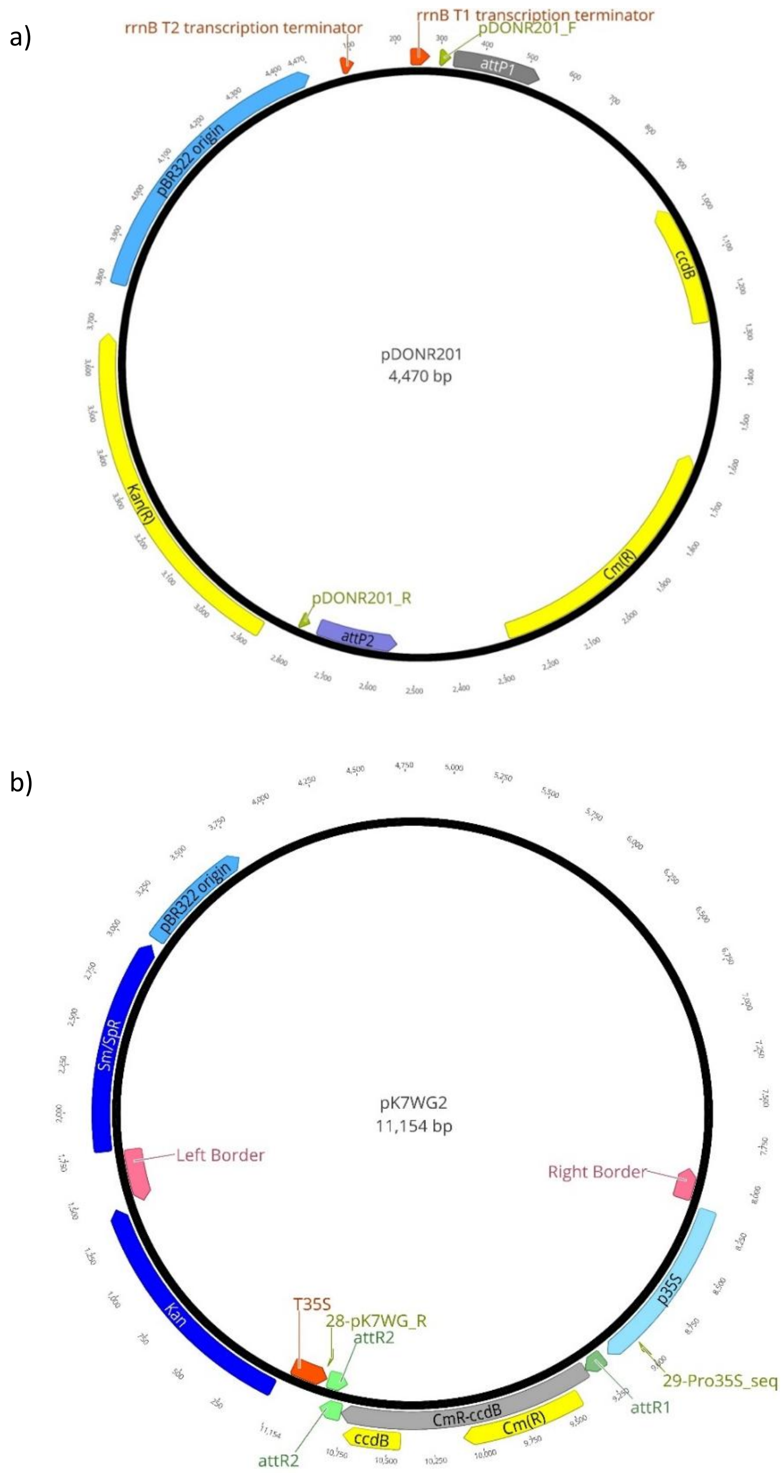


**Figure M5:** Binary vector sets of the *p35S* over-expression system. The vector sets a) pDONR201 (Invitrogen Life technologies) and b) pK7WG2 (Karimi *et al.*, 2002) were used for the over-expression.

The initial aim to generate the over-expression system was to amplify the CDS of the KTI candidates via standard PCR. The amplified CDS must also comprise of the attachment sites (att sites) necessary for the compatibility to the Gateway cloning system. Thus, the primer sets, forward and reverse of the CDS also contained 5’-

GGGGACAAGTTTGTACAAAAAAGCAGGCTTAAAA-3' and 5'-GGGGACCACTTTGTACAAGAAAGCTGGGTATTAAGCGTAATCTGGAACATCGTATGGGTAC-3' respectively, for the inclusion of the attachment sites to the amplified CDS. The standard PCR condition has been described in Additional Table S1. The primer sets were ordered from Microsynth Seqlab, Göttingen.

As mentioned above, the primer sets were used to amplify CDS via standard PCR but using Phusion™ DNA polymerase (Thermo Fisher Scientific) instead of *Taq* DNA polymerase (Thermo Fisher Scientific). The amplified sequences were integrated into the entry vector-pDONR201 by adding 5 µl of 100 – 200 ng of amplified CDS, 2.5 µl of pDONR201 (150 ng), 5 µl of 5X BP Reaction Buffer, 5 µl of BP Clonase™ enzyme mix (Thermo Fisher Scientific) and 7.5 µl of pH 8.0 TE buffer (preparation mentioned in section 1.2.5). These components were mixed well and left overnight at room temperature. 1.2 U of Proteinase K was added the next morning and was incubated at 37 °C (Thermomixer, Thermo Fisher Scientific) for 10 min. 5 µl of the newly transformed vector was transformed into *E. coli* strain DH5a (as described in section 1.2.2) and plated on LB solid medium with kanamycin at 37 °C (Table M1). Overnight colonies from the LB plates were tested with colony PCR as described in section 1.2.3 with the primers pDONR201_F+ pDONR201_R (Additional Table 1). Positive colonies were used for plasmid extraction after overnight culture at 37 °C in LB and kanamycin and were validated with Sanger sequencing using the primers pDONR201_F and pDONR201_R (primer details in Additional Table S1). The final concentration of the extracted plasmids was checked in a spectrophotometer (NanoDrop™ One , Thermo Fisher Scientific) and was normalized to 100 ng by diluting in autoclaved demineralized water.

The region containing the CDSs of the respective candidate *KTIs* in pDONR201 (Figure M5a), was transferred to the destination vector-pK7WG2, which contained the *p35S* promoter (Figure M5b). The Gateway™ LR Clonase™ II Enzyme mix (Thermo Fisher Scientific) was used for the integration of the CDS of the candidate gene into the destination vector pK7WG2. In a 1.5 mL reaction tube (Sarstedt AK & Co.KG), 2 µl of pDONR201 vector+CDS (100 ng), 3 µl of pK7WG2 (50 ng), 4 µl of TE buffer (pH 8.0, preparation mentioned in section 1.2.5) and 1 µl LR Clonase II (U/µl) were added and incubated at room temperature overnight. Proteinase K of 0.6 U was added the next day and incubated at 37 °C (ThermoMixer, Thermo Fisher Scientific) for 10 min. The constructs were transformed into *E. coli* cells strain DH5α (section 1.2.2 for transformation method) and tested with colony PCR (section 1.2.3) with the primers 29-Pro35S_Seq+28-pK7WG_R (T35S) (Additional Table S1). The positive colonies were incubated to 3 mL of LB liquid culture together with spectinomycin at 37 °C with continuous shaking at 180 rpm (shaker ZWYR-240, Labwit Manufacturer). Plasmid was extracted the next morning (section 1.2.4) and was tested further for the positive Gateway reaction with a Sanger sequencing with the primer 28-pK7WG_R (T35S) (Additional Table S1) from Microsynth Seqlab. The final vector construct generated with the introduction of the candidate *KTI* CDSs were, pK7WG2_400, pk7WG2_600 and pk7WG2_53200 (Figure M6).

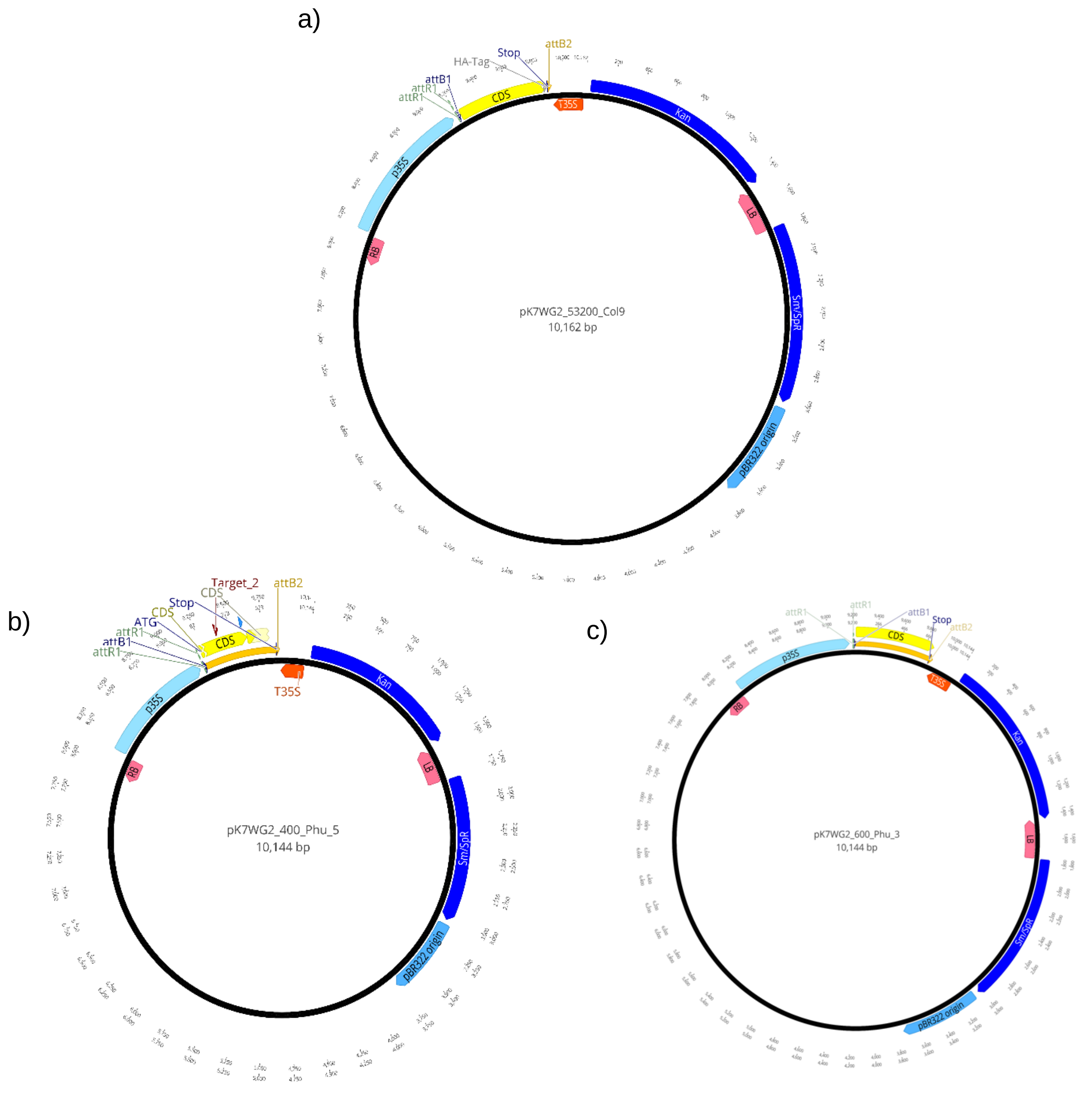


**Figure M6:** Vector maps generated for the over-expression of the candidate *KTIs* expressed under *p35S*. Vector sets were generated via the recombination of cDNA of *KTI_53200* forming a) pk7WG2_53200*;* cDNA of *KTI_400* forming b) pk7WG2_400 and cDNA of *KTI_600* forming c) pk7WG2_600*.* Vector maps generated in Geneious Prime (Biomatters, Ltd., Auckland, New Zealand).

The verified plasmid comprising CDS from the candidate *KTIs* expressed under the *p35S* promoter was transformed into *Agrobacterium* (section 1.2.2 for transformation). Colony PCR was performed with the primer sets 29-Pro35S_Seq+28-pK7WG_ R (Additional Table S1) to validate the transformation of the correct plasmid to *Agrobacterium.* A stock culture of *Agrobacterium* transformed with the desired plasmid was prepared by mixing 1:1 (v:v) *Agrobacterium* culture in YEB:50 % sterile glycerol. The culture was aliquoted, shock-frozen in liquid nitrogen, and stored as cryo-glycerol stock cultures at -80 °C.

The empty vector control was generated by transforming the vector pK7WG2 lacking any CDS from candidate *KTIs* into *Agrobacterium* (section 1.2.2 for transformation).

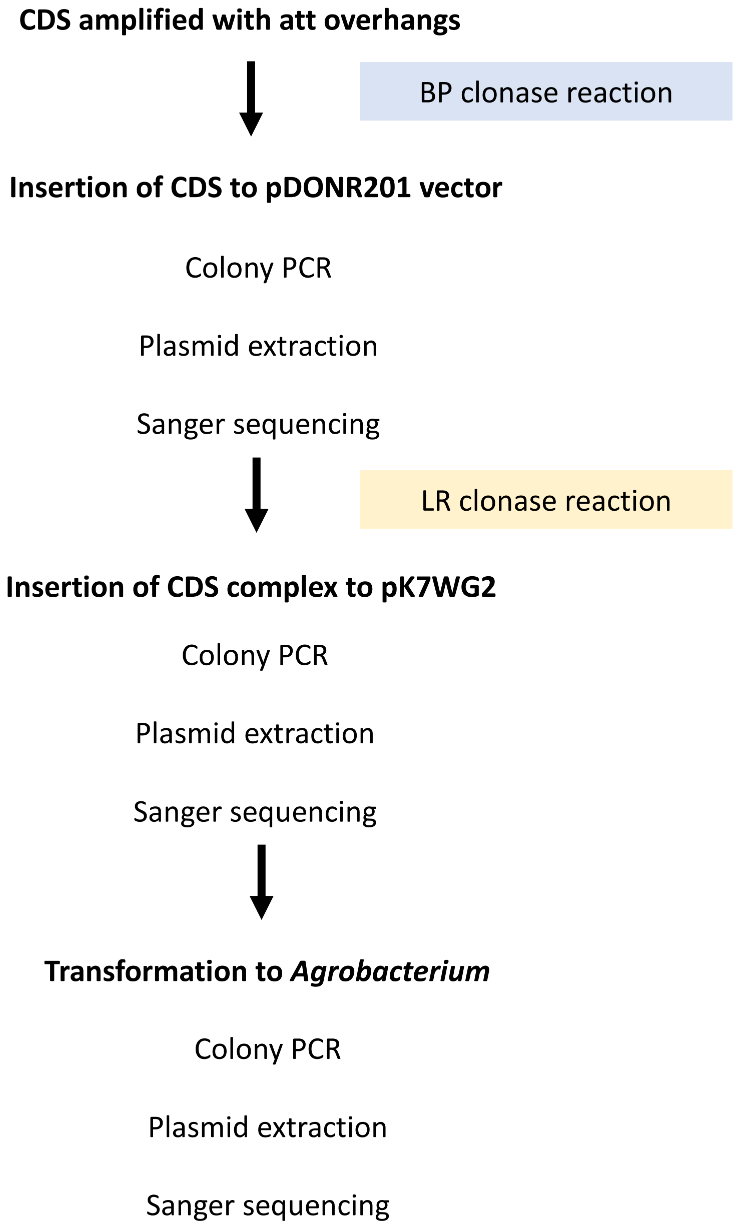


**Figure 7:** Schematic overview of the molecular cloning of *p35S* over-expression system.

#### 1.2.7 Transformation of poplars

The poplar transformation was adapted from Bruegmann *et al.* 2019. Several modifications were made during this study, as outlined below. The overview scheme of the transformation procedure is shown in Figure M8.

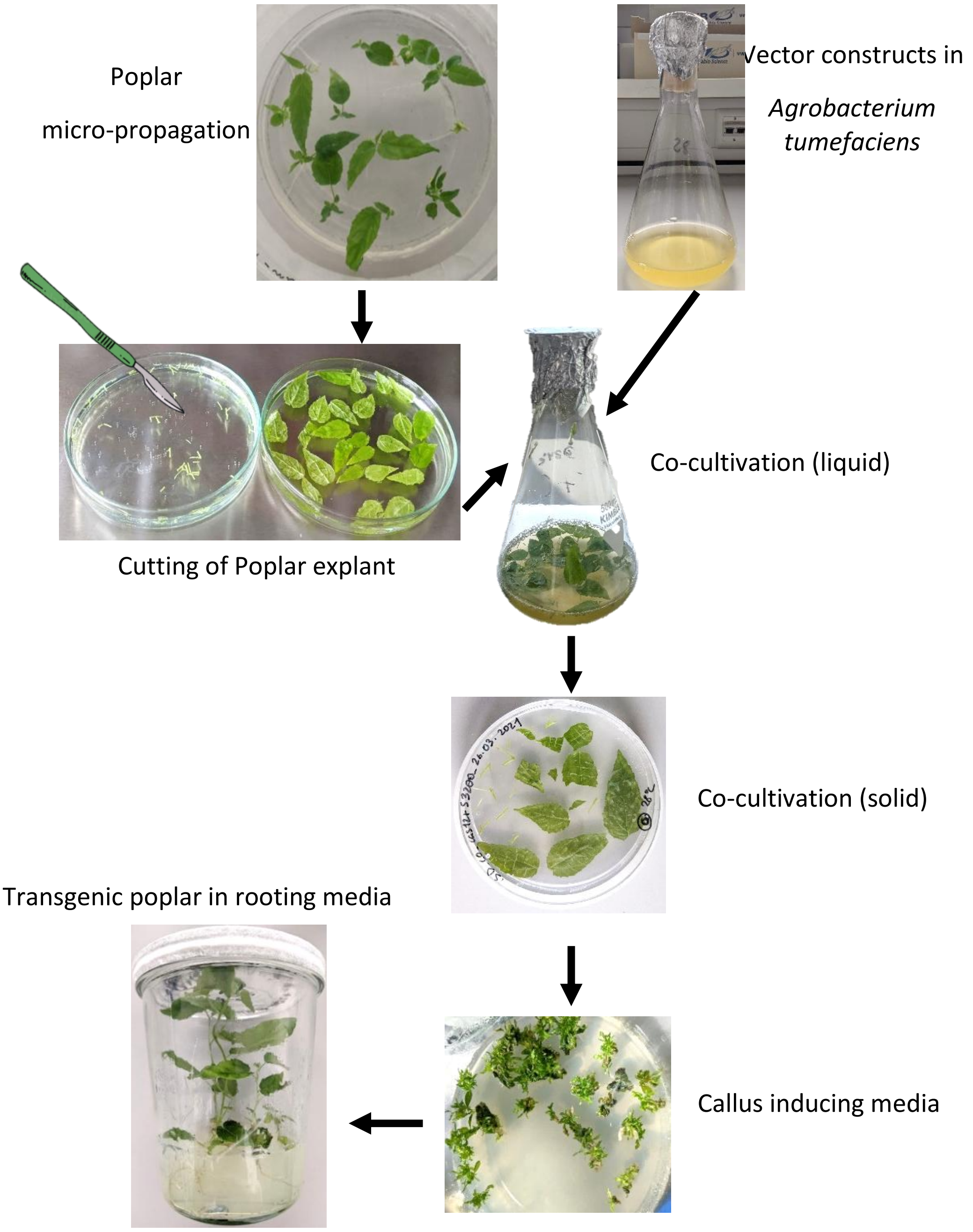


**Figure M8:** Scheme of the *in-vitro* transformation of *Agrobacterium tumefaciens* into *Populus* x *canescens.*

The desired *Agrobacterium* culture, stored as cryo-stocks, was used for *in-vitro* transformation into *P.* x *canescens.* Initially, a streak on a YEB plate, with rifampicin, gentamicin, and spectinomycin (Table M1), was performed and incubated overnight at 28 °C (incubator ZWYR-

240, Labwit Manufacturer). The next day, a portion of the streak was picked with a sterile toothpick and incubated with 4 mL of liquid YEB, containing rifampicin, gentamicin, and spectinomycin (Table M1). The culture was incubated at 28 °C in darkness with constant shaking at 160 rpm (ZWYR-240, Labwit Manufacturer) overnight. The overnight culture was poured into 100 mL of YEB (in a conical flask, pre-warmed at 28 °C; M1) with continuous shaking at 160 rpm (ZWYR-240, Labwit Manufacturer) at 28°C in ambient light conditions. $OD_{600}$ was checked regularly in a photometer (Eppendorf AG) until it reached values from 0.3 to 0.8. Then, 0.2 µl of acetosyringone from the stock solution (Carl Roth GmbH) was pipetted to the culture. Acetosyringone was prepared as a 200 mM stock dissolved in ethanol and sterile-filtered (Filtropur s 0.2 µm, Sarstedt GmbH). The culture was further shaken for 0.5 h before use.

Poplars to be transformed were prepared approximately three weeks before the transformation. They were grown in ½ MS media under plant cultivation room conditions (16 h light, 70-85 µE $m^{-2}$ $s^{-1}$ PAR, and 24 °C, 60 % RH, light source: L18W/840, Osram). For transformation, leaves were excised from the stem, and stems were further cut to approx. 0.5 cm in length on a sterile glass petri-dish under a clean bench (Thermo Fisher Scientific). Leaves of sterile poplars were used for transformation by making fine slits into the whole leaves (Figure M8) (Song et al., 2019). During the excision procedure, all tissues were soaked in sterile ½ MS media (Figure M8). The apical shoots of the plantlets were not used for transformation.

The slit leaves and cut stems ("transforming tissues") were separated from ½ MS liquid media using a sterilized stainless-steel sieve. The transforming tissues were placed in the previously prepared acetosyringone+YEB medium (having *Agrobacterium* culture, Table M1) and were shaken in darkness at 28 °C (shaker ZWYR-240, Labwit Manufacturer), 120 rpm for 30 min. The transforming tissues were sieved from the *Agrobacterium* culture and were dried with sterilized filter paper (Whatman paper, Merck KGaA) under a sterile bench (Thermo Fisher Scientific) and were transferred to the co-cultivation medium (½ MS media with sterile-filtered 200 mM acetosyringone, added after autoclaving). Co-cultivation plates having transforming tissues were incubated at 28 °C in darkness for 2 -3 days in the climate cabinet (AR-75L, Percival Scientific, 60 % RH).

Co-cultivated explants were washed two times, that is, in the beginning, and in the end, with autoclaved (21 °C, 2.2 bar for 20 minutes; 6x6x6 HST, Zirbus technology) tap-water, and in-between, three times with sterile water containing 400 mg/L ticarcillin (sterile-filtered, Duchefa Biochemie B.V.). All washing steps were performed for 2 min under sterile conditions with constant shaking. The final washed explants were placed in the callus-inducing medium (½

MS medium with cefotaxime (150 mg/L); ticarcillin (200 mg/L), and TDZ-Pluronic (Thiadiazuron from Duchefa Biochemie, dissolved in Poloxamer 188 solution from Sigma-Aldrich at 0.01 % w:v; sterile-filtered through 0.2 µm sieve). As selectable markers, for the positive selection of CRISPR-Cas12a plant transformants, callus-inducing media was supplemented with gentamicin (60 mg/L). Similarly, for the selection of *p35S*-based over-expression system plant transformants, kanamycin (50 mg/L) was used.

Callus-inducing media with explants were placed in climatized cabinets (AR-75L, Percival Scientific) at 28 ° C, 20 µE $m^{-2}$ $s^{-1}$ PAR, 60 % RH, 16 h of light, light source: Alto 32 Watt (Philips, Amsterdam, Netherlands) and LG4507.4 (Megaman, Langenselbold, Germany) for four weeks. Shoots were separated from the calli and transferred to the rooting media (same as callus-inducing media, with TDZ-Pluronic) and grown at 28 °C, 60 µE PAR with 16 h light for 4 weeks, light source: Alto 32 Watt and LG4507.4 (Figure M8).

#### 1.2.8 Plant growth tests for using gentamicin as a selectable marker

Callus-inducing and rooting media (see section 1.2.7 for the media recipe) were prepared with a concentration gradient of gentamicin of 0 mg/L, 30 mg/L, and 90 mg/mL. Gentamicin containing both media was prepared and poured into squared petri-dishes. Stems, petioles, and slit leaves from non-mutagenized wild-type plants were placed on the callus-inducing and rooting media. The rooting media were cut in half to clearly observe the root development, and single stems from the wild-type plants were placed vertically into the cut media. The rooting media with the sterile stems were placed vertically (to encourage gravitropism), whereas the callus-inducing media was placed horizontally to the surface. All media with the sterile explants were placed in a plant cultivation room at 16 h light, 70-85 µE $m^{-2}$ $s^{-1}$ PAR, and 24 °C, 60 % RH, light source: L18W/840, Osram) The growth of calli and roots were recorded for four weeks.

#### 1.2.9 Genotyping of putative mutant poplars

All rooted plants growing in the rooting media were used for genotyping. For this purpose, plantlets were potted to the N-type soil as a substrate in 12x12x12 cm pots and were transferred to greenhouse conditions. The third fully developed leaf from the top was collected from 10 weeks old mutant poplars, flash-frozen in liquid nitrogen, and used for DNA extraction. Samples were tested for T-DNA presence by performing standard PCR and using the primer sets LbCas12a_F_2+LbCas12a_R_2 for plants transformed with the CRISPR-Cas12a system and 29-Pro35S_Seq+28-pK7WG_R (T35S) for the *p35S* based over-expression system (Additional Table S1). The presence of vector backbone was also checked for the CRISPR-Cas12a system with primers Spec_F_2+Spec_R_2 (Additional Table S1).

To test for gene editing via the CRISPR-Cas12a system, gene-specific standard PCR was performed. Any difference of the DNA sequence of the transformed lines relative to WT was stated as a gene editing event.

To test for the transcript abundance of the p35S mutant lines, RNA was extracted with innuPREP RNA kit (Analytik Jena GmbH) from the leaf sample of a single plant. Then, 1 µg RNA was used for cDNA synthesis. RT-qPCR was performed with three technical replicates (as described in 1.2.9) with the primer sets, KTI_3_F+ KTI_3_R_2 for KTI_400, KTI_4_F_2+ KTI_4_R_4 for KTI_600 and KTI_5_F+ KTI_5_R_2 for KTI_53200 (Additional Table S1).

**References**

Bergkessel M, Guthrie C. Chapter twenty five - colony PCR. In: Lorsch J, editor. Methods in Enzymology, vol. 529, Academic Press; 2013, p. 299–309. https://doi.org/10.1016/B978-0-12-418687-3.00025-2.

Bruegmann T, Polak O, Deecke K, Nietsch J, Fladung M. Poplar transformation. Methods Mol Biol 2019;1864:165–77. https://doi.org/10.1007/978-1-4939-8778-8_12.

Das IS. Functional characterization of protease inhibitors involved in induced systemic defenses in poplar. Göttingen: 2024.

Dower WJ, Miller JF, Ragsdale CW. High efficiency transformation of *E. coli* by high voltage electroporation. Nucleic Acids Res 1988;16:6127–45.

Hartley JL, Temple GF, Brasch MA. DNA cloning using in vitro site-specific recombination. Genome Res 2000;10:1788–95. https://doi.org/10.1101/gr.143000.

Karimi M, Inzé D, Depicker A. Gateway vectors for Agrobacterium-mediated plant transformation. Trends in Plant Science 2002;7:193–5. https://doi.org/10.1016/S1360-1385(02)02251-3.

Katzen F. Gateway recombinational cloning: A biological operating system. Expert Opin Drug Discov 2007;2:571–89. https://doi.org/10.1517/17460441.2.4.571.

Merker L, Schindele P, Huang TK, Wolter F, Puchta H. Enhancing in planta gene targeting efficiencies in Arabidopsis using temperature-tolerant CRISPR/LbCas12a. Plant Biotechnology Journal 2020. https://doi.org/10.1111/pbi.13426.

Schindele P, Puchta H. Engineering CRISPR/LbCas12a for highly efficient, temperature-tolerant plant gene editing. Plant Biotechnol Journal 2020;18:1118. https://doi.org/10.1111/pbi.13275.

Sharma RC, Schimke RT. Preparation of electrocompetent *E. coli* using salt-free growth medium. Biotechniques 1996;20:42–4. https://doi.org/10.2144/96201bm08.

Song C, Lu L, Guo Y, Xu H, Li R. Efficient *Agrobacterium*-Mediated Transformation of the Commercial Hybrid Poplar *Populus alba* × *Populus glandulosa* Uyeki. Int J Mol Sci 2019;20:2594. https://doi.org/10.3390/ijms20102594.

Taketo A. DNA transfection of *Escherichia coli* by electroporation. Biochim Biophys Acta 1988;949:318–24. https://doi.org/10.1016/0167-4781(88)90158-3.
